# Red light-emitting short Mango-based system enables tracking a mycobacterial small noncoding RNA in infected macrophages

**DOI:** 10.1101/2022.09.07.506834

**Authors:** Oksana S. Bychenko, Alexei A. Khrulev, Julia I. Svetlova, Vladimir B. Tsvetkov, Polina N. Kamzeeva, Yulia V. Skvortsova, Boris S. Tupertsev, Igor A. Ivanov, Leonid V. Aseev, Yuriy M. Khodarovich, Evgeny S. Belyaev, Liubov I. Kozlovskaya, Timofei S. Zatsepin, Tatyana L. Azhikina, Anna M. Varizhuk, Andrey V. Aralov

## Abstract

Progress in the studies of RNA metabolism and function relies largely on molecular imaging systems, including those comprising a fluorogenic dye and an aptamer-based fluorescence-activating tag. G4 aptamers of Mango family, typically combined with a duplex/hairpin scaffold, activate fluorescence of a green-emitting dye TO1-biotin and hold great promise for intracellular RNA tracking. Here, we report a new Mango-based imaging platform. Its key advantages are tunability of spectral properties and applicability for visualization of small RNA molecules that require minimal tag size. The former advantage is due to an expanded (green-to-red-emitting) palette of TO1-inspired fluorogenic dyes, and the latter is ensured by the truncated duplex scaffold. To illustrate applicability of the improved platform, we tagged *Mycobacterium tuberculosis* sncRNA with the shortened aptamer-scaffold tag and visualized it in bacteria and bacteria-infected macrophages using the new red light-emitting Mango-activated dye.

## INTRODUCTION

Small noncoding RNAs (sncRNAs) have recently emerged as important modulators of transcription and translation. Many eukaryotic sncRNAs are differentially expressed in cancer^1, 2^, neurodegenerative diseases^3, 4^, and other pathologies, which makes them attractive drug targets or biomarkers. Prokaryotic sncRNAs govern stress response, adaptation, signaling, and virulence-related processes^5^. Of particular interest is their contribution to immune evasion^6^. The underlying mechanisms require clarification, hence the necessity for sncRNA imaging and tracking in living cells.

Despite recent advances in encoding fluorogenic RNA or intracellular RNA labeling for fluorescent microscopy imaging^7, 8^, available methods are hardly tuned for sncRNA. The key challenge stems from the size factor. Lengthy/bulky tags, such as arrays of fluorescent protein binding sites, may interfere with sncRNA interactions, traffic, or phase transitions. A perfect tag should be shorter than or at least comparable to a wild-type small RNA (40-500 nt). A more general problem to be solved is contrast enhancement. Systems based on genetically encoded RNA-binding fluorescent proteins suffer from substantial background signal, encouraging the development of light-up alternatives. Finally, for simultaneous tracking of two or more distinct RNAs, orthogonal dual-color or multiplex imaging systems are much needed.

Most of the above challenges and problems have been addressed to some extend in recent imaging systems based on an aptamer and a fluorogenic dye^9^. Such systems comprise a small molecule dye, which is supposed to be non-fluorescent in a free state, and a ‘turn-on’ RNA aptamer to this dye, which is fused with RNA of interest. In robust imaging systems, intracellular formation of the aptamer-dye complex causes several order of magnitude increase of fluorescence quantum yield. SELEX has provided a plethora of aptamers that meet the above ‘turn-on’ criterion and show high specificity for the respective dye, including Spinach, Broccoli, Corn, Chilli, Mango, Coral, Malachite Green, Riboglow, Pepper, and Peach^10, 11, 12, 13, 14, 15, 16, 17, 18^. However, each of these systems suffers from more or less pronounced drawbacks associated with difficult synthesis, low fluorescence enhancement, inability to penetrate cell membranes, low stability in live cells, long length of the tag, and limitations on detection channels.

In this paper, we report new Mango-specific fluorogenic dyes and the label length optimization. In dyes design, we focused on the following three problems: (1) simplification of fluorogenic dye synthesis; (2) enhancement of the fluorescence quantum yield in the bound state, and (3) adjustment of the emission range. The dyes were tested against two well-characterized Mango aptamers, namely Mango II and Mango IV^19^, and the effects of structure alterations on the fluorescence quantum yield were elucidated by molecular modeling. Additionally, to adjust the system for sncRNA visualization and tracking, the total length of the RNA imaging tag (F30-Mango II: 88 nt^19^) was shortened by truncating one of the stems of F30 folding scaffold. This resulted in the shortest terminal label for livecell RNA imaging to date (ds_Mango II: 52 nt). Applicability of the optimized tag and the new dye with improved characteristics for intracellular imaging and tracking of regulatory sncRNA was verified using an intracellular internalization of *Mycobacterium smegmatis* by a host macrophage.

## RESULTS

### Design and synthesis of the dyes

The design of the new dyes was inspired by the thiazole orange derivatives TO1-biotin, TO3-biotin and YO3-biotin (Fig. 1). These derivatives were first synthesized as partners for RNA Mango aptamer (later known as Mango I). TO1-Biotin was successfully used to purify Mango I-tagged transcription-regulating RNA 6S and visualize Mango I-based construct in *C. elegans*^20^, and YO3-biotin was combined with DFHBI-1T/Spinach pair to obtain a FRET-based system^21^. Despite the promise for bioimaging, application of TO1-biotin, TO3-biotin, and YO3-biotin is limited, partly because intermediates for their synthesis are reportedly obtained in very low yields ranging from 6 to 12%^20, 21^. The yields of the final amide bond-forming reaction are typically not presented, but one can expect moderately low efficiency of the proposed transformation or the difficulties with HPLC purification.

**Fig 1.**
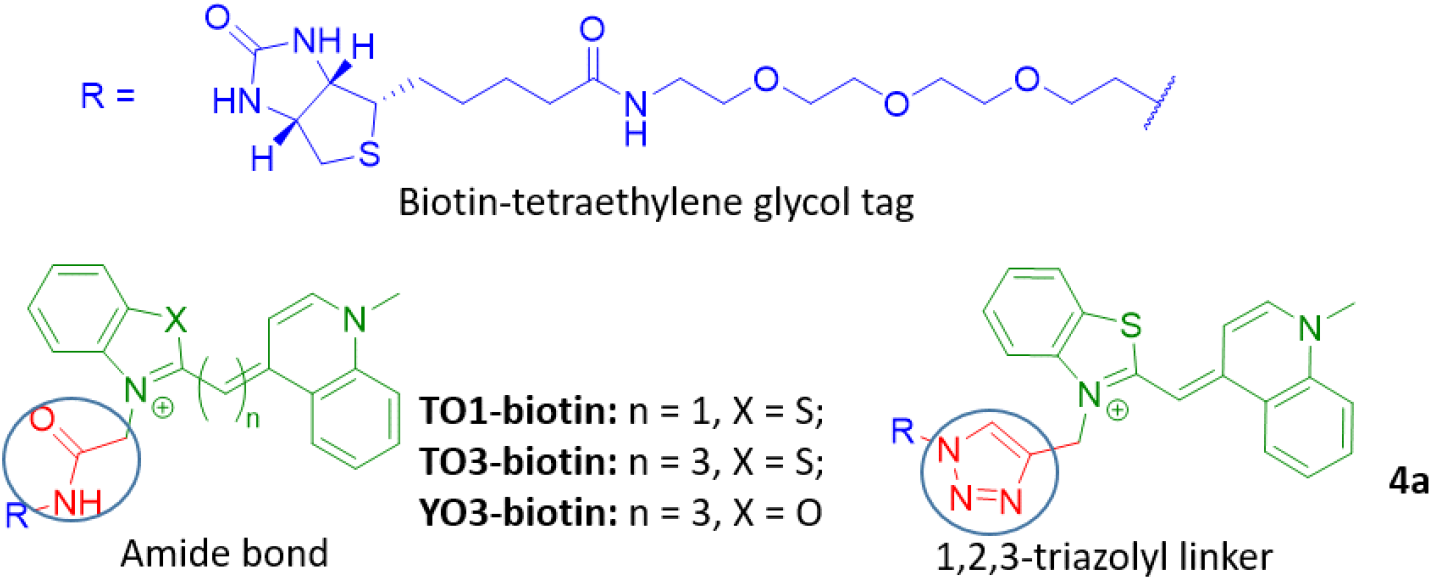
The structures of reported fluorogenic dyes TO1-biotin, TO3-biotin and YO3-biotin and triazolyl-linked TO1-biotin analog **4a**.

“Click” chemistry is a synthetic approach for efficient and highly selective conjugating molecules with a high yield^22^. The most popular variant of “click” chemistry seems to be a 1,3-dipolar cycloaddition under copper-(I) catalysis between terminal alkyne function and an azide moiety that leads to the formation of the 1,2,3-triazole cycle^23, 24^. The replacement of an amide bond with isosteric 1,2,3-triazolyl might simplify synthetic transformations, increase their efficiency and, in some cases, improve the properties of the biomolecules^25,26,27,28,29^.

The structure of TO1-biotin consists of two moieties, a biotin-tetraethylene glycol tag and a fluorophore, linked by an amide bond (Fig. 1). Here, we decided to replace this bond with an isosteric 1,2,3-triazolyl linker in order to increase the efficiency of synthetic transformations and evaluate its influence on the light-up effect upon binding with fluorogen-activating aptamers of Mango family.

TO1-biotin was prepared according to the literature^20^. For the synthesis of triazolyl-linked analog **4a**, benzothiazolium salt **1** was condensed with 1-methylquinolinium salt in the presence of TEA to afford alkyne-modified dye **2a** with a yield of 43%, which is ~7 times higher than the yield for the corresponding intermediate in the TO1-biotin synthesis^20^. Following the reported procedure for copper catalyzed azide— alkyne cycloaddition («click» reaction)^30^ (**Scheme 1, see Supplementary, chemistry**), alkyne-containing derivative **2a** and azido-containing biotinylated derivative **3**^31^ were condensed in the presence of copper (I) iodide, TBTA and DIPEA affording, after preparative HPLC purification, the target fluorogenic dye **4a** as trifluoroacetate salt. To verify the efficiency of Cu(I)-catalyzed azide–alkyne 1,3-dipolar cycloaddition (CuAAC) reaction compared to the amide bond-forming one, we analyzed crude reaction mixtures containing TO1-biotin and **4a** using LS-MS (Figs. S1 and S2). The purity assessment at absorption wavelengths of 260 and 500 nm (the latter wavelength is close to the maximum absorption wavelength of TO1-biotin and **4a**) gave 1.2 and 9.2% for TO1-biotin (R_f_ = 5.65 min) *versus* 22.4 and 76.5% for **4a** (R_f_ = 5.86 min). The reaction yields were 1.8 and 3.5% for TO1-biotin and **4a**, respectively, after a single HPLC purification, presumably due to the low solubility of both derivatives in the loading buffer (water with 0.1% TFA). However, after several iterations of HPLC purification overall yield of TO1-biotin and **4a** reached 6.8 and 58.1 %, respectively (see Supplementary).

**Scheme 1.**
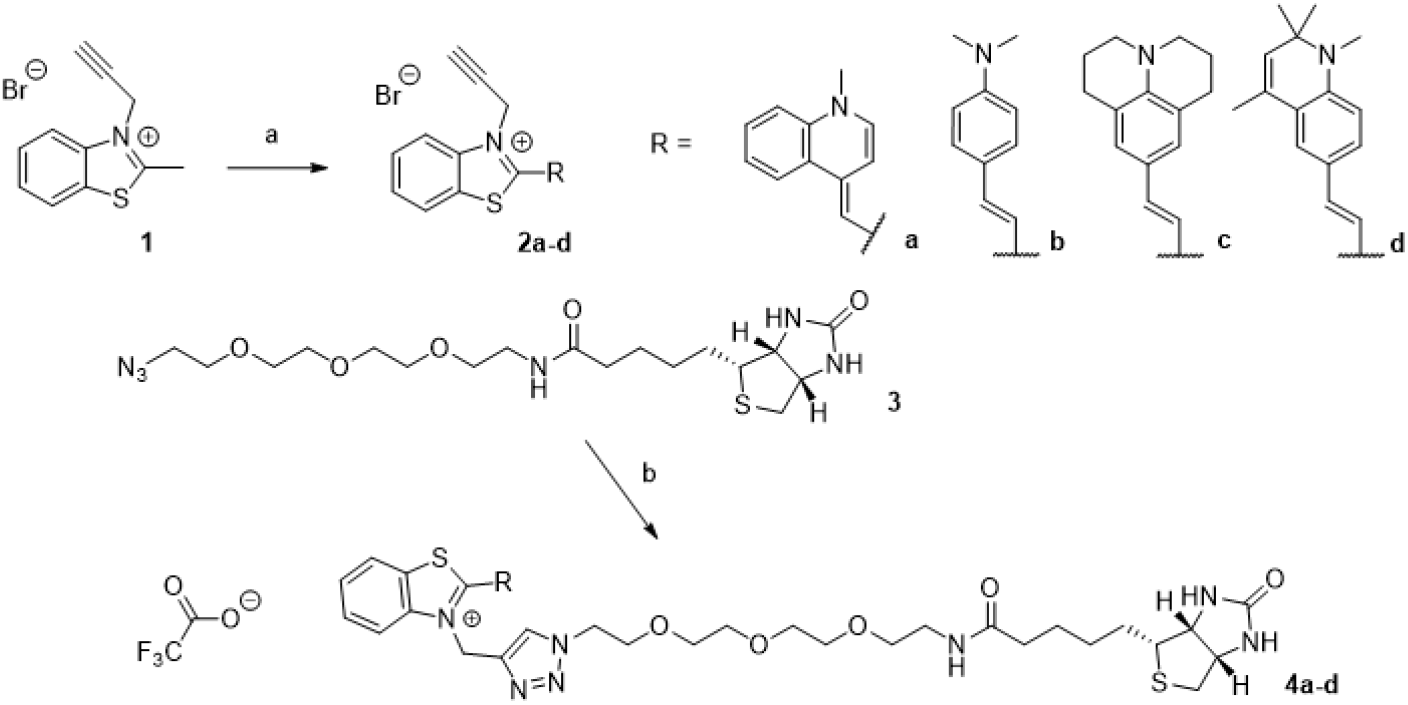
Synthesis of TO1-biotin analogs. Reagents and conditions: a) 1-methylquinolinium bromide, TEA, CH_2_Cl_2_ (for **2a**) or 4-(dimethylamino)benzaldehyde (for **2b**), 2,3,6,7-tetrahydro-1H,5H-pyrido[3,2,1-ij]quinoline-9-carbaldehyde (for **2c**), 1,2,2,4-tetramethyl-1,2-dihydroquinoline-6-carbaldehyde (for **2d**), Ac_2_O, reflux; b) **2**, CuI, TBTA, DIPEA, CH_3_OH/CH_3_CN, rt, then preparative HPLC.

Inspired by the chemical design of TO3-biotin and YO3-biotin with extended conjugated systems, which are significantly red-shifted but show drastically reduced fluorescence enhancement upon aptamer binding with respect to the parent TO1-biotin^20, 21^, and our finding on the utility of simple and effective «click» reaction, we next explored the possibility of using other cyanine derivatives as fluorogenic dyes for Mango-based RNA labeling systems. In the design of the dyes with the red-shifted emission spectra, D–π–A systems, where D and A are electron donor and acceptor, respectively, finds broad application^32^. Accordingly, we left the electron-withdrawing benzothiazolium ring^33^ in the cyanine push-pull system but replaced 1-methylquinoline ring with strong electron-donating moieties. We also replaced the methine bridge between the donor and acceptor fragments with an ethenyl linker. Thus, we synthesized the dyes containing *N*,*N*-dimethylaminophenyl (**4b**), 2,3,6,7-tetrahydro-1H,5H-pyrido[3,2,1-ij]quinolin-9-yl (**4c**), or 1,2,2,4-tetramethyl-1,2-dihydroquinolin-6-yl (**4d**) group as an electron donor. For this, benzothiazolium salt **1** was condensed with an appropriate aromatic aldehyde via an aldol-type reaction in acetic anhydride^34^ to afford the alkyne-modified derivatives **2b-d** (**Scheme 1, see Supplementary, chemistry**). Their condensation with azido-containing biotinylated derivative **3** in the presence of copper (I) iodide, TBTA and DIPEA yielded, after preparative HPLC purification, fluorogenic dyes **4b-d** as trifluoroacetate salts.

### Spectral properties, affinity and specificity

Spectral properties of the new dyes **4a-d** and the control dye TO1-biotin were investigated in a free state and in complexes with aptamers Mango II and Mango IV (for sequences see Table S1). The former one shows enhanced thermal stability compared to Mango I, is resistant to formaldehyde fixation, and has been comprehensively evaluated for bioimaging^19, 35, 36^. The latter one is the second actively used aptamer of the Mango family^37^.

First, we tested the dyes for disrupting or altering secondary structures of the aptamers. At a concentration of 5 μM, which is well above the reported Kd values for Mango complexes with TO1-biotin^19^, the dyes did not induce significant changes in absorbance or circular dichroism (CD) bands of Mango II and Mango IV, except for the minor decrease of CD amplitude (Fig. 2A). At the same time, the dyes showed red-shifted absorbance spectra in equimolar mixtures with the aptamers compared to the free state (Fig. 2B). Consistently with previous reports for TO1-biotin^19, 35^, these data support complex formation and the maintenance of the G4 core in the complexes.

**Fig 2.**
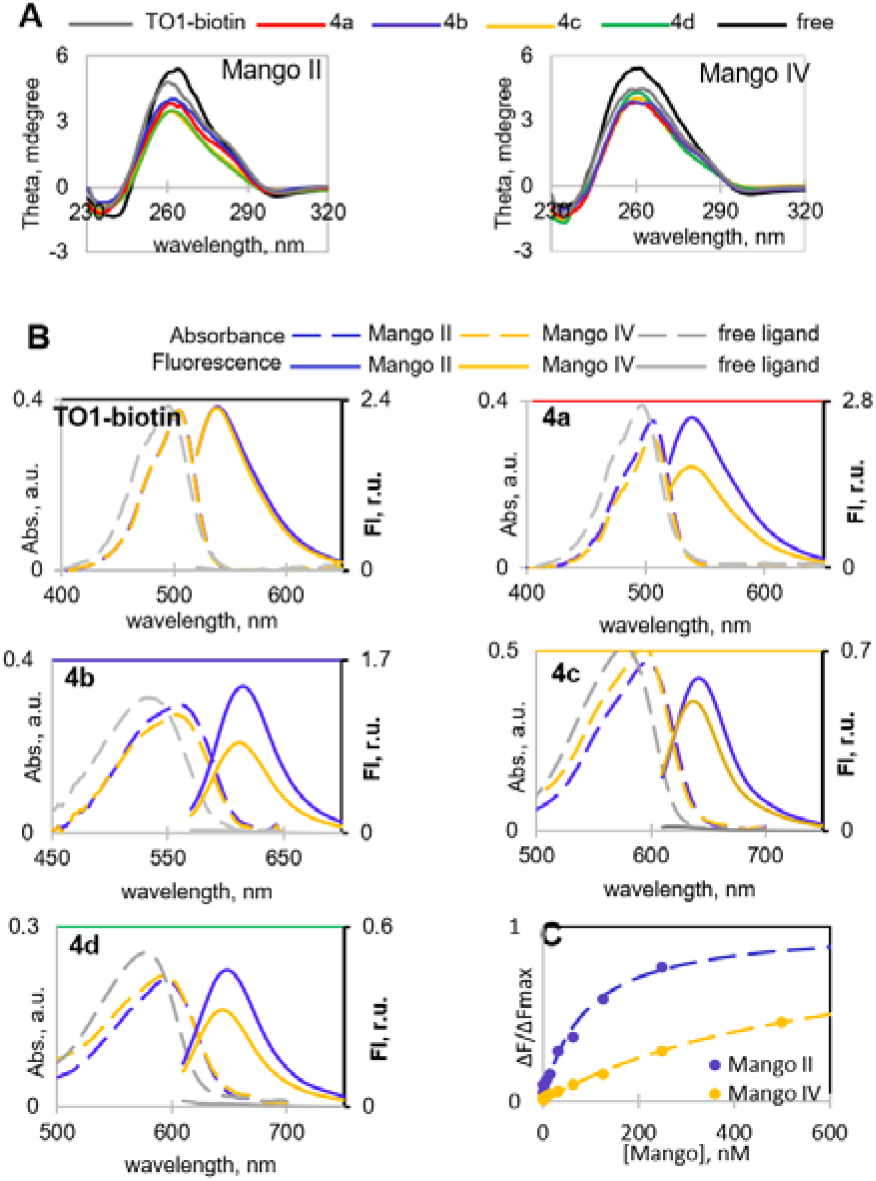
TO1-biotin and new dyes 4a-d in complexes with Mango II and IV aptamers: spectral properties and binding affinity. (A) Verification of dye impact on Mango secondary structure by CD spectroscopy. (B) Absorption and fluorescence emission spectra of the free dyes and their complexes. (C) **4b**-Mango binding assay. Conditions in A and B: 5 μM Mango aptamer, 5 μM dye, 10 mM Tris-HCl, pH 7.5, and 140 mM KCl. Conditions in C: 10 nM dye.

Next, we compared the dyes based on their key light-up properties, including increase of fluorescence and brightness in the bound state. Excitation at a complex-specific absorbance maxima (Table 1, Fig. S3) gave negligible fluorescence emission of the dyes in a free state (Fo), while equimolar mixtures with the aptamers showed up to nearly three orders of magnitude increase in fluorescence emission (F/Fo, Table 2). While TO1-biotin proved to be an equally efficient light-up probe for Mango II and Mango IV with a fluorescence quantum yield (Φ) equal to 0.43, its triazole-linked analog **4a** showed preference for Mango II (Φ = 0.42) compared to Mango IV (Φ = 0.32) and a slightly reduced brightness (Table 1). Interestingly, **4a** exhibited higher contrast (F/Fo) than TO1-biotin, presumably because the 1,2,3-triazolyl linker reduced the rotational flexibility of heteroaromatic systems compared to the amide bond. Dye **4b** was also a more robust probe for Mango II than Mango IV. It showed a red-shifted spectra with an increased Stokes shift (Δλ = 55 nm) compared to TO1-biotin and **4a** (Δλ = 35 nm). Dyes **4c** and **4d** were inferior to other variants and showed substantially reduced fluorescence quantum yields and brightness (Table 1).

**Table 1.** Spectral properties of the dyes in complexes with the aptamers

| Dye | free dye<br>abs. max<br>(nm) | free dye<sup>a</sup><br>$\epsilon$ (M<sup>-1</sup>cm<sup>-1</sup>) | Mango-dye<br>$\lambda_{ex}$<br>(nm) | Mango-dye<br>$\lambda_{em}$<br>(nm) | $\Phi$ Mango-dye<br>(r.u.) | | Brightness_Mango-dye<br>(M<sup>-1</sup>cm<sup>-1</sup>) | |
| --- | --- | --- | --- | --- | --- | --- | --- | --- |
|  |  |  |  |  | Mango II | Mango IV | Mango II | Mango IV |
| TO1-biotin | 495 | 77500 | 505 | 540 | 0.43 | 0.43 | 32200 | 32200 |
| **4a** | 495 | 77500 | 505 | 540 | 0.42 | 0.32 | 29400 | 19900 |
| **4b** | 535 | 62400 | 560 | 615 | 0.44 | 0.29 | 26200 | 15900 |
| **4c** | 580 | 101100 | 595 | 640 | 0.07 | 0.05 | 7100 | 5400 |
| **4d** | 580 | 51300 | 595 | 650 | 0.14 | 0.10 | 6400 | 4700 |

<sup>a</sup> Molar extinction at absorbance maximum of the free dye
Brightness = abs at complex max\* $\Phi$

**Table 2.** Specificity of the light-up effects

| dye | F/Fo<sup>a</sup> |  |  |  |  |  |  |  |  |  |
| --- | --- | --- | --- | --- | --- | --- | --- | --- | --- | --- |
|  | Mango II | Mango IV | control G4 RNA<sup>b</sup> | dsRNA<sup>b</sup> | ssRNA<sup>b</sup> | dsDNA<sup>b</sup> | ssDNA<sup>a</sup> | h/pG4-DNA<sup>a</sup> | aG4-DNA<sup>a</sup> | i-motif<sup>a</sup> |
| TO1-biotin | 380 | 380 | 240 (1.6)<sup>c</sup> | 22 (17) | 14 (27) | 4 (95) | 2 (190) | 9 (42) | 8 (48) | 7 (54) |
| **4a** | 824 | 550 | 572 (1.4) | 74 (11) | 38 (22) | 36 (23) | 6 (137) | 54 (15) | 49 (17) | 51 (16) |
| **4b** | 93 | 56 | 19 (4.9) | 7 (13) | 6 (16) | 2 (47) | 2 (47) | 4 (23) | 3 (31) | 2 (47) |
| **4c** | 55 | 47 | 30 (1.8) | 4 (1.8) | 3 (18) | 5 (11) | 3 (18) | 6 (9) | 5 (11) | 4 (14) |
| **4d** | 59 | 41 | 17 (3.5) | 2 (30) | 2 (30) | 5 (12) | 4 (15) | 6 (10) | 5 (12) | 4 (15) |

<sup>a</sup> Fluorescence emission in complex (F) normalized by fluorescence of the free dye (Fo) at a Mango-specific max.
<sup>b</sup> ssDNA is a random-sequence RNA (25 µg/mL); for other ON sequences, see Table S1.
<sup>c</sup> Selectivity toward Mango II compared with control RNA or DNA secondary structure as a ratio of F/Fo<sup>Mango II</sup> to F/Fo<sup>control</sup> is shown in parentheses.

To evaluate binding selectivity of the reported and new dyes towards their RNA aptamers, we compared light-up effects of dyes in the presence of Mango II, Mango IV, and other DNA/RNA structures using a set of control oligonucleotides (ONs) (Table S1). The set included the control G4 RNA *utr-z* from the human KRAS 5’-UTR sequence^38^, random-sequence ssRNA from salmon, the RNA hairpin rds26 and the DNA hairpin ds26^39, 40^, the telomeric DNA fragment 22AG^41, 42, 43^, which adopts a parallel-stranded or a hybrid-type G4 structure, along with its mutant 22CTA, which adopts an antiparallel-stranded G4 structure^44^, and a DNA fragment from C9ORF72 repeat expansions, which adopts an i-motif structure at near-physiological conditions^45^. None of these control ONs was comparable to Mango in terms of the light-up effect (Table 2). Dyes **4a-d** showed somewhat reduced selectivity towards Mango over DNA structures (~10X Mango:ODN light-up ratio) compared to TO1-biotin (up to 2 order of magnitude Mango:ODN light-up ratio). Among the new dyes, top selectivity was observed for **4b** (Table 2). In terms of selectivity towards Mango over RNA (especially G4 RNA), **4b** outperformed both **4a** and TO1-biotin (Table 2 and Fig. S4). For this dye, binding affinity toward Mango aptamers was evaluated in a fluorescence light-up assay, which gave Kd values of 80±60 nM and 0.6±0.4 μM for Mango II and Mango IV, respectively (Fig. 2C).

Considering improved selectivity of **4b** for MangoII/IV over randomly selected G4 RNA and its profound Stokes shift, we questioned whether it could be applied in combination with another dye/aptamer pair for multiplex imaging or FRET-based monitoring of RNA-RNA juxtaposition^21, 37^. Recently, fluorescent RNA aptamers Spinach and Mango I with their cognate fluorophores were used to develop an RNA apta-FRET system that responded to RNA conformational changes in *E. coli*^21^. We evaluated the possibility of using **4b**-mango II pair as an acceptor paired with Broccoli/DFHBI donor^11^ for constructing FRET systems. Broccoli aptamer was synthesized as previously described^46^ (for sequence, see Table S2). Its CD spectrum (Fig. S5a) was consistent with a presumed G4 structure. Upon titration with Broccoli aptamer, DFHBI showed a bathochromic absorbance shift (Fig. S5b) and became fluorescent (Fig. S5c) when excited at 470 nm (absorbance maximum of the DFHBI-Broccoli complex) with emission maximum at 510 nm. GFP-derived fluorophores are reportedly insensitive to Mango aptamers^21, 37^. Thus, selectivity of **4b** to Mango over Broccoli was the defining question with respect to DFHBI-Broccoli and **4b**-Mango II orthogonality. Broccoli-induced absorbance changes (Fig. S5d) and fluorescence enhancement (Fig. S5e) of **4b** were negligible compared to those induced by Mango II, indicating no interference between the aptamer-dye pairs. At the same time, the pairs showed substantial donor-acceptor spectra overlap (Fig. 3), suggesting efficient FRET upon donor-acceptor juxtaposition. To summarize this part, **4b**-Mango II and DFHBI-Broccoli pairs appear to be a perfect match and have prospects in multiplex or FRET-based RNA tracking.

**Fig 3.**
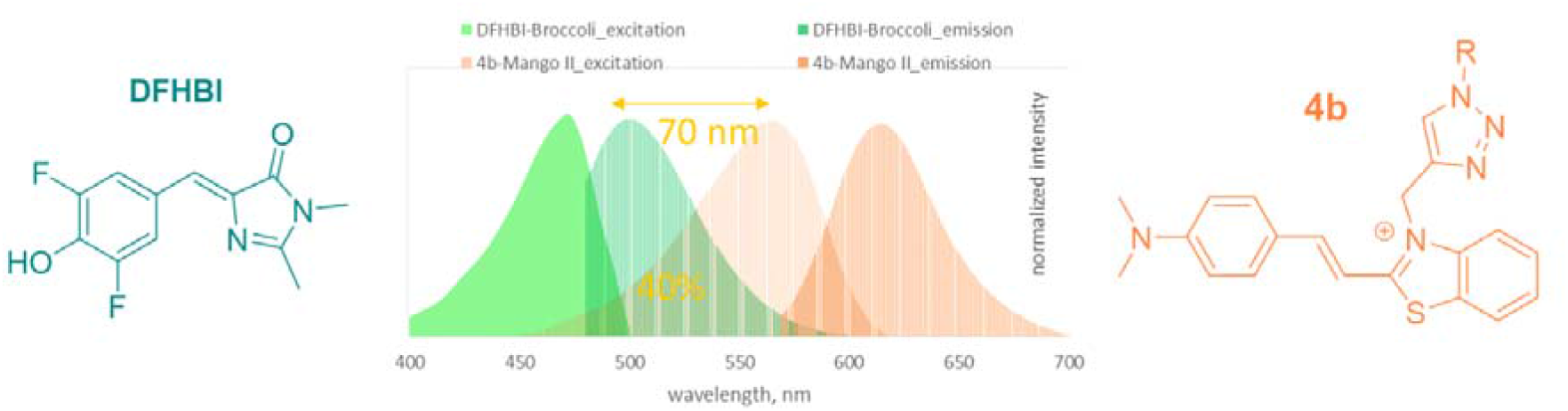
Excitation and emission spectra of DFHBI and 4b in complex with Broccoli and Mango II aptamers, respectively.

### Molecular modeling

To elucidate the molecular basis for the high light-up effects of **4a** and **4b** (Table 1), we modeled their complexes with Mango II based on the reported structure of TO1-biotin-Mango II^35^ (Fig. 4a). Dye **4c**, which showed poor light-up effect, was also analyzed as a semi-negative control. All dye models were truncated slightly by removing the biotin-tetraethylene glycol tag but keeping the amide bond in TO1-biotin and 1,2,3-triazolyl linker in **4a-c.** Amide/1,2,3-triazole linkers adjacent to the fluorogenic heteroaromatic system may impact relative positioning of the N-methylquinolinyl (MQ) moiety (TO1-biotin and **4a**) or its substitute (**4b** and **4c**) and the benzothiazolium (BT) group in the dye-Mango II complexes and thus might affect fluorescence enhancement. The biotin-containing tag itself is unlikely to make a major contribution, so it was excluded from the analysis. The dyes were localized on the surface of the external Mango G-quartet by docking, and the stability of the resulting complexes was tested by molecular dynamics (MD) simulations.

**Fig. 4.**
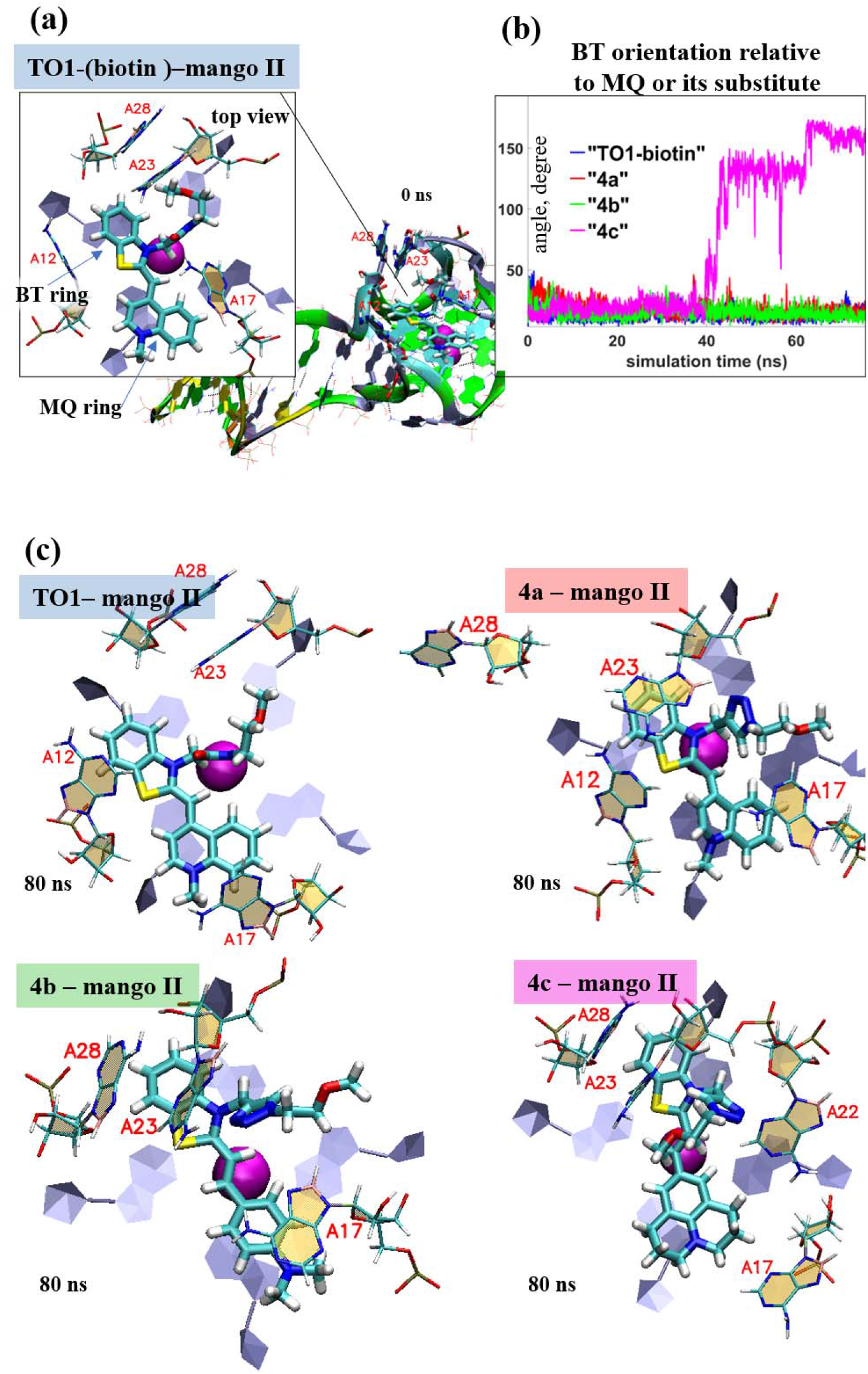
Dynamics of Mango II complexes with TO1-biotin and its analogs: planarity vs rotational mobility of dye residues. (a) Initial model of TO1-biotin-Mango II complex (0 ns MD simulation snapshot) illustrating starting positions of benzothiazolium (BT) and methylquinolinyl (MQ) moieties, their stacking with external Mango II G4 tetrad and orientation relative to key neighboring Mango II residues (A12, A17, A23, and A28). Biotin residue is not shown and was excluded from the analysis. (b) Evolution of the angle between the planes of BT ring and MQ ring (TO1-biotin and **4a**) or its substitute (**4b-c**). (c) End-point conformations of the complexes (80 ns MD simulation snapshots), top views. Color labeling of aptamer residues in TO1-biotin-Mango II full structure (a): G, green; A, grey; T, orange; C, cyan. Aptamer residues in complex fragment, top view: C, beige; G, grey. Ligand atoms: C, cyan; N, blue; O, red; S, yellow.

Key result of the 80 ns simulation that illustrates mobility of the fluorogenic heteroaromatic system within the complexes is shown in Fig. 4b. While BT and MQ rings in TO1-biotin and **4a**, as well as BT ring and *N,N*-dimethylaminophenyl (DAP) moiety in **4b** remained roughly planar throughout the simulation, in **4c** the 2,3,6,7-tetrahydro-1H,5H-pyrido[3,2,1-ij]quinolin-9-yl (THPQ) residue underwent full (180 degree) rotation. End-point structures of the complexes (top views of the dyes and the neighboring aptamer residues) are shown in Fig. 4c (for full structures, see Fig. S6). The observed rotation of THPQ, which caused the energy loss and thus reduced the fluorescent quantum yield of **4c**, likely results from inefficient stacking interactions between THPQ and A17. In contrast, MQ of TO1-biotin and **4a**, as well as DAP of **4b**, formed stable stacking contacts with A17, as was evident from the plots of COM distances and the angles between the normals and planes of these residues (Fig. S7).

The 1,2,3-triazole ring did not interfere with the dye geometry, but formed additional stacking contacts with the neighboring aptamer residue A23, which led to minor disruption of the local aptamer structure (A23-A26 stacking) in **4a**-Mango II complex. This might explain the slightly inferior performance of **4a** compared to TO1-biotin. In **4c**-Mango II complex, the 1,2,3-triazole ring formed additional contacts with A22, and in general **4c** induced a particularly pronounced alteration of local aptamer environment, as was evident from increased RMSD (Fig. S8). The **4b**-Mango II complex showed minimal fluctuations at the aptamer-dye interface (Fig. S7), which explains its top performance in fluorescent enhancement assays (Table 1). Despite the lower aptamer binding affinity of **4b** compared to TO1-biotin, it showed the lowest free energy of complex formation (mainly due to van der Waals interactions) among all new dyes (Fig. S9).

### Application of the new 4b-Mango II system for live-cell imaging and tracking of the mycobacterial sncRNA

The sncRNAs of intracellular pathogenic bacteria are powerful regulators of bacterial adaptation to the host’s immune defense^6, 47^. In this work, we tested **4b** for intracellular imaging of a *M. tuberculosis* small RNA MTS1338, detecting this sncRNA in bacteria *in vitro*, and tracking in infected macrophages. For this purpose, we used the modular RNA (ds_Mango II_MTS1338) consisting of MTS1338 fused with ds_Mango II (the optimized fluorescence-activating tag) that was prepared by truncating one of the stems in F30 folding scaffold (Fig S10). The sequences of the modular RNA and the genetic constructs used in the work are given in Table S2. The ability of the modular RNA bearing two functional domains, MTS1338 and the fluorogen-activating tag, to form a correct secondary structure, according to the RNAfold server (http://rna.-tbi.univie.ac.at/cgi-bin/RNAWebSuite/RNAfold.cgi), was shown (the free energy of the thermodynamic ensemble is −87.79 kcal/mol, Fig S10).

To exclude cytotoxicity-related artifacts, the effects of **4b** and other dyes on the metabolic activity of host-cell macrophages were verified. Cytotoxicity of the dyes was evaluated using murine macrophages (RAW 264.7 cell culture, ATCC® TIB-71™). No changes in cell morphology (data not shown) or metabolic activity were observed after 24 h incubation with the dyes (Fig. S11). At day 7 even at 50 μM concentration, none of the compounds caused a decrease in the level of cell metabolism by more than 50%, suggesting CC_50_ > 50 μM for all tested compounds. However, **4c** and **4d** at this concentration provoked morphological changes in cells (data not shown). We also performed cytotoxicity assays for bacterial cells. For bacteria, no differences were found in growth rate in the presence/absence of **4b** in the medium (Fig. S12).

Modular RNA and control ds_Mango II were obtained using *in vitro* transcription, and selective light-up effects of **4b** and the control dye TO1-biotin upon their interactions with these RNA were confirmed in a cuvette. Consistently with the data in Fig. 2, both dyes showed pronounced fluorescence in the presence of the Mango II-tagged RNA but not the tag-free (negative control) RNA (Fig. S13).

Next, we generated *Mycobacterium Smegmatis* (MSmeg) clones expressing the modular RNA, ds_Mango II, and the empty vector-transformed clone (as a negative control). The non-pathogenic *Mycobacterium smegmatis* mc(2)155 was used in the study, because its genome and metabolism are very similar to *Mycobacterium tuberculosis*, MSmeg also has a significantly higher growth rate and much easier undergoes genetic modifications, therefore is widely used as a surrogate organism^48^. MSmeg does not contain the MTS1338 gene.

Visualization of the modular RNA and ds_Mango II in live bacterial cell is shown in Fig. 5. The images demonstrate high fluorescence of modular RNA and ds_Mango II complexes with both TO1-biotin and **4b**. Since the imaging of live bacteria was performed in solution, the fluorescent signals from the construct/dye complexes (in green/red channels) do not always coincide with those generated by Hoechst 33258 (blue channel) due to the bacteria’s movement. No bright green or red signals were detected in experiments with bacteria transformed by empty vector pAMYC and stained with TO1-biotin or **4b**.

**Fig 5.**
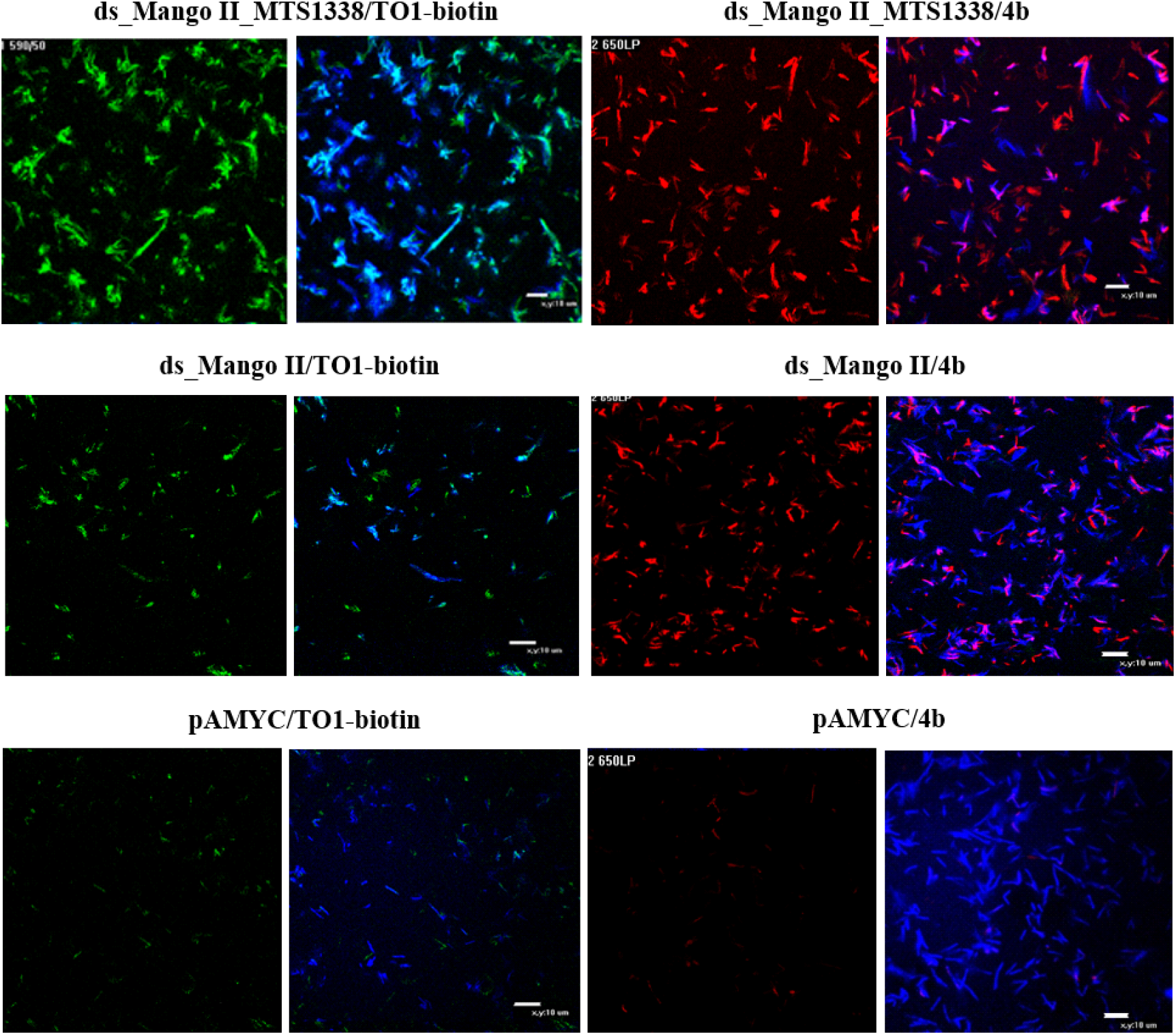
Visualization of the genetically encoded modular RNA ds_Mango II_MTS1338 and ds_Mango II in *M. smegmatis* in solution using TO1-biotin (in a green channel) and **4b** (in a red channel). Bacteria transformed by pAMYC without insertion were used as a negative control. Bacteria were stained with Hoechst 33258 (in a blue channel).

When macrophages infected with MSmeg recombinant strains expressing modular RNA or ds_Mango II were stained with TO1-biotin, the fluorescent bacteria were detected suggesting a correct folding of the label’s secondary structure (Fig. S14).

To further explore the dissemination and possible secretion of MTS1338 within infected live macrophages we monitored the changes in fluorescent signal of modular RNA and ds_Mango II complexes with **4b** for 1 and 1.5 hour (Fig. 6 and 7). Inspection of individual infected cells allowed observing a certain migration of the red fluorescent signal from the bacterial cell inside the infected macrophage, while the outlines of bacteria gradually disappeared. At the same time, no such signal migration was observed in macrophages infected with a control strain expressing ds_Mango II. Instead, we observed gradual attenuation of the signal and the disappearance of the bacterium. Similar to the results obtained by visualizing MTS1338 in infected macrophages with Broccoli aptamer^46^, this may indicate the MTS1338 secretion into the cytoplasm of macrophages and is consistent with the presumed function of MTS1338^49^. Although additional experiments are required to further characterize MTS1338 functions, the data obtained support the role in bacteria-host interactions. These results confirm successful application of the developed imaging system (short RNA imaging tag + red light-emitting fluorogenic dye) for visualization of sncRNAs within infected living cells.

**Fig 6.**
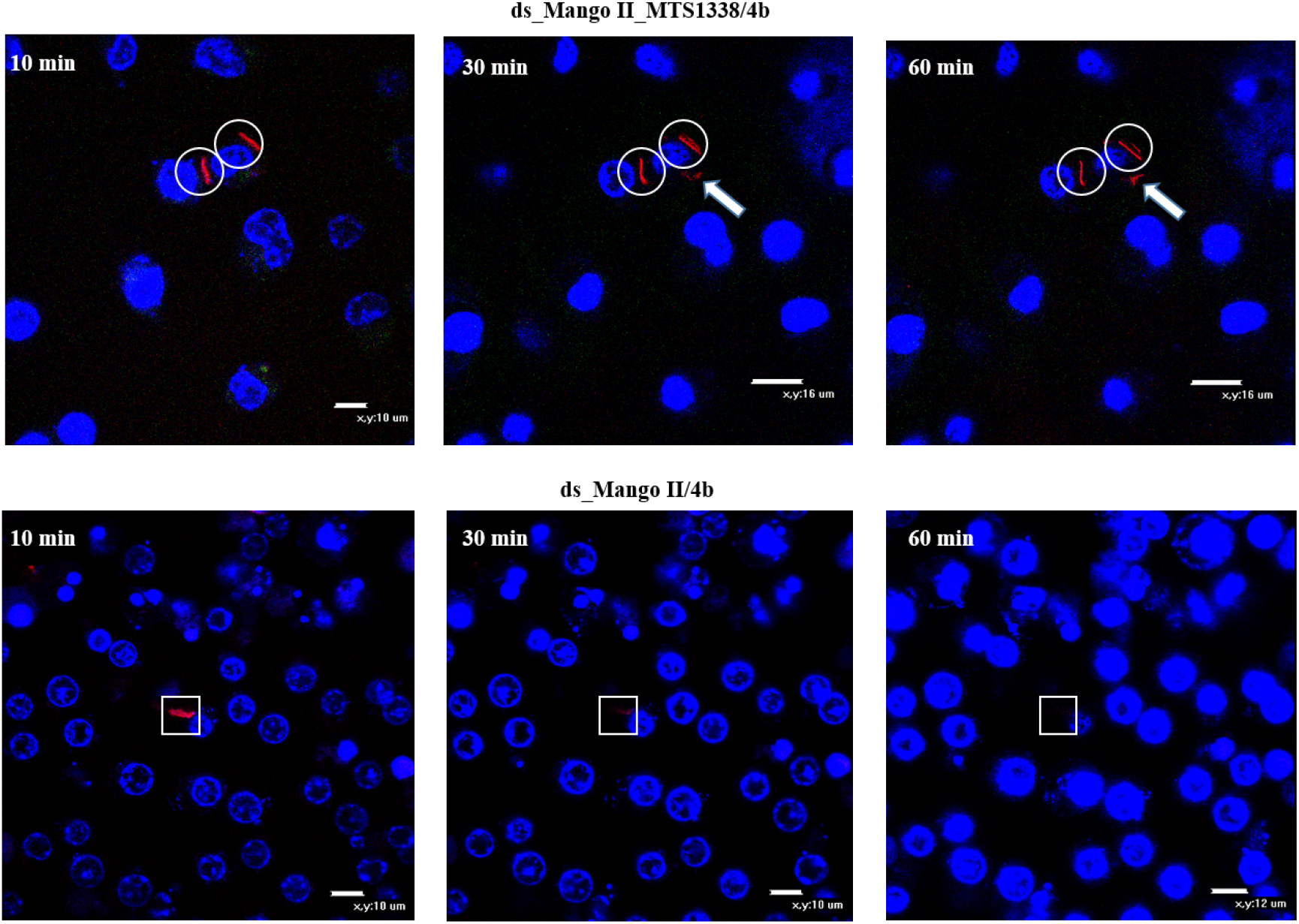
Tracking of the red fluorescent signal for 1 hour (10, 30, and 60 min), using the genetically encoded modular RNA or ds_Mango II and **4b** in infected RAW 264.7 macrophages (wide field images). Macrophage nuclei were stained with Hoechst 33258. The circles cover the signal maintenance; white arrows show fluorescent signal movement/occurrence while squares highlight the area of signal disappearance.

**Fig 7.**
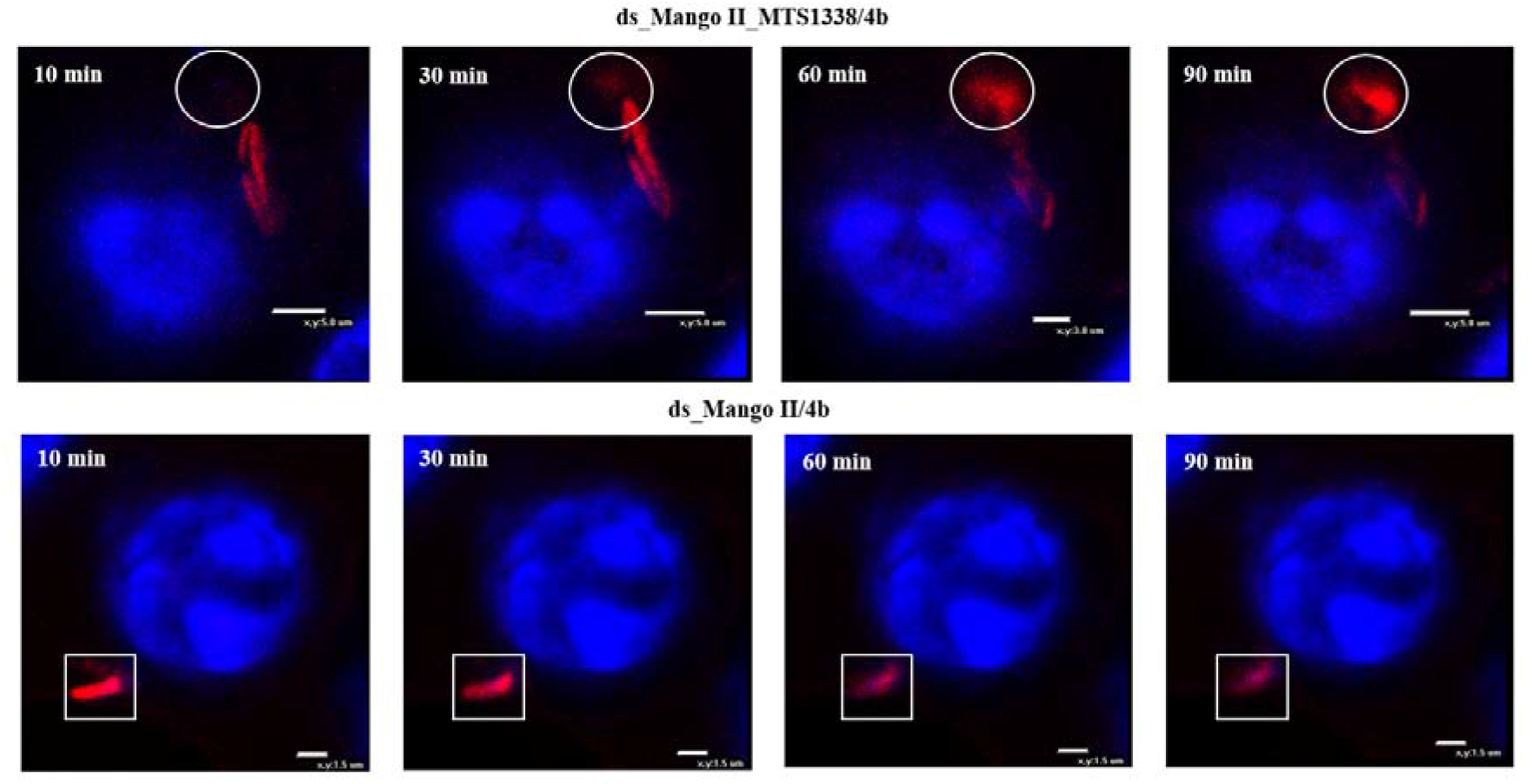
Tracking of the red fluorescent signal for 1.5 h (10, 30, 60, and 90 min), using the genetically encoded modular RNA or ds_Mango II and **4b** in infected RAW 264.7 macrophages (enlarged scale). Macrophage nuclei were stained with Hoechst 33258. The circles show fluorescent signal movement/occurrence, while squares highlight the area of signal disappearance.

## Discussion

RNA aptamers that recognize and activate small molecule fluorogenic dyes have shown immense potential in the field of RNA visualization, including the development of FRET for monitoring RNA-RNA interactions and their structural rearrangements within the living cell. Among available aptamer-dye pairs, only the Pepper-based imaging system appears to cover the entire range of visible light emission^16^. However, the discriminating ability of Pepper aptamers for dyes structurally dissimilar to their cognate partners has not yet been studied, leaving Pepper applicability for the design of FRET constructs under question. To ensure biocompatibility, an increase in Pepper thermal stability and a decrease in its magnesium-ion-dependence are needed^50^. As concerns usability for tracking small RNA, a minimal (43 nt) aptamer obtained by SELEX with subsequent optimization has not been exploited so far. For tagging natural RNAs of interest, such as 5S ribosomal RNA, 7SK small nuclear RNA and U6 splicing RNA, the initially selected relatively long (61 nt) aptamer D11 has been used^16^.

Mango-based imaging systems also look very promising with respect to both the fluorogenic dye (TO1-biotin) and the aptamers (Mango I-IV and others)^19, 35, 51^. TO1-biotin, stemming from the thiazole orange (TO) core, shows no significant cytotoxicity and is remarkable for high photostability, extinction coefficient, quantum yield and brightness^52, 53, 54, 55^. Mango aptamers are notable for their in-cell stability and high binding affinity toward TO1-biotin as well as a very short length (29-32 nt), albeit an additional F30 folding scaffold (58 nt) is required for livecell imaging^19, 56^.

Here, an efficient scheme for the synthesis of analogs of TO1-biotin, originally proposed as a fluorogenic dye for Mango aptamers^20^, has been developed. Very poor yields of the original transformations prompted us to use simple and effective CuAAC «click» reaction instead of the common amide bond-forming one (Scheme 1). As a result, **4a**, a TO1-biotin analog with the same thiazole orange (TO) fluorogenic fragment, was prepared in an overall yield of 25 % which is ~60 times higher than in a classical route to TO1-biotin^20^. The use of a different synthesis strategy led to the replacement of the amide bond by a 1,2,3-triazolyl linker, thus requiring the study of its effect on the dye spectral properties and the fluorescence-activating ability of Mango aptamers. To evaluate this, the most studied and stable in living cells aptamers Mango II and Mango IV were selected. The new dye **4a** exhibited spectral properties almost similar to the original dye upon activation by Mango II (Table 1). Inspired by these results and previous synthetic design of YO3-biotin, we also replaced the TO fluorogenic fragment with other residues that potentially could provide red-shifted fluorescence. Among them, **4b** with, *N,N*-dimethylaminophenyl moiety instead of 1-methylquinolinyl, showed slightly lower brightness than TO1-biotin and **4a**, but at the same time had a red-shifted emission maximum of 615 nm (Table 1). The proposed dye in complex with Mango II had absorption and emission wavelengths similar to complexes of YO3-biotin with MIII AU10 (Mango III mutant) and Peach aptamers^18^. Despite reduced brightness (26200 M^-1^cm^-1^ for **4b-**Mango II *versus* 77300 and 47600 M^−1^cm^−1^ for YO3-biotin-MIII AU10 and YO3-biotin-Peach, respectively), **4b** in complex with the stable Mango II provides the opportunity for live-cell imaging. In contrast to Mango III and its variants have only been long studied only in a cuvette, and Peach aptamers wait in the wings.

Another important issue is the contrast-defining selectivity of dyes for cognate aptamers over abundant intracellular DNA and RNA, which has not been systematically studied to date. Consistently with previous reports^20^, TO1-biotin showed selectivity for Mango aptamers over model RNA in our assays. The selectivity over RNA secondary structures was generally inferior to that over DNA structures (Table 2). Importantly, TO1-biotin and **4a** demonstrated only moderate preference for Mango II over genomic RNA G4 *utr-z* from the human KRAS 5’-UTR sequence^38^ (1.6 and 1.4-fold difference in fluorescence enhancement, respectively), while **4b** showed improved selectivity (4.9-fold difference in fluorescence enhancement, Table 2,).

Recently, a FRET system based on RNA aptamers Spinach and Mango I that is genetically encodable and respond to conformational changes have been developed^21^. Here, we reported a preliminary assessment of the applicability of **4b**-Mango II for developing a FRET system with DFHBI-Broccoli pair that has the brightness comparable to DFHBI-1T-Spinach, due to the high fluorescence-activating ability of the Broccoli aptamer^11^. In terms of the maximum possible FRET efficiency, **4b**-Mango II in combination with DFHBI-Broccoli (40% donor-acceptor overlap upon normalization of all emission/excitation spectra to a maximum intensity of 1) appeared superior to other known FRET pairs, namely DFHBI-Spinach with TO3-Mango I (23% spectral overlap) and DFHBI-1T-Spinach with YO3-biotin-Mango I (30% overlap) (Fig. 3)^21^. TO3-biotin-Mango I (acceptor) demonstrated a relatively large (120 nm) excitation redshift relative to the emission of DFHBI-Spinach (donor), hence a modest spectral overlap. Substitution of TO3-biotin for YO3-biotin resulted in a 40 nm excitation blueshift, which improved the spectral overlap by approximately 30%. Excitation of **4b**-Mango II was blue-shifted further by 20 nm, and its overlap with the donor was improved by additional 30% compared to YO3-biotin-Mango I. Due to the excitation maximum blue-shift of **4b** and its greater Stokes shift compared to YO3-biotin (Δλ = 55 nm and 40 nm, respectively), **4b**-Mango II in combination with DFHBI-Broccoli appear particularly suitable for excitation by 488/555 nm lasers and detection using common EGFP and Texas Red filters.

There are examples of how noncoding bacterial RNAs of intracellular pathogens can be secreted into the cytoplasm of infected cells and modulate the immune response during infection^57, 58^. Small *M. tuberculosis* RNA MTS1338 is presumed to be one of such secreted regulators and affects the protective cells of the immune system^59^. To verify this assumption, a system for visualizing small bacterial RNAs using fluorescence microscopy is in demand. Previously, we demonstrated the possibility of detecting sncRNA MTS1338 by fluorescence microscopy with the persistence of bacteria in macrophages using F30-2xdBroccoli aptamer^46^. However, the F30-2xdBroccoli sequence is twice as long as the RNA under study (MTS1338-117 bp vs F30-2xdBroccoli-234 bp) and may thus interfere with RNA function. To enable artifact-free tracking of the sncRNA in living cells, a shorter tag and a dye with enhanced contrast and selectivity are much needed. In this study, we truncated one of the stems of F30 folding scaffold in F30-Mango II tag (88 nt)^19^ resulting in the shortest terminal ds_Mango II label (52 nt) for live-cell RNA imaging reported to date (Fig. S10). The optimized tag and the new dye **4b** with improved characteristics for intracellular imaging/tracking of regulatory sncRNA was verified using MTS1338. We visualized the modular RNA and ds_Mango II in live bacterial cells using **4b** and TO1-biotin as a control of the correct aptamer folding (Fig. 5). The fluorescence intensity was comparable for both dyes and higher in bacterial cells transfected with the modular RNA-encoding plasmid, which is probably due to the label-stabilizing ability of MTS1338.

Correct folding of RNA secondary structure in macrophages infected with strains expressing modular RNA or ds_Mango II was confirmed by staining with TO1-biotin (Fig. S14). Finally, we tracked red fluorescence signal of **4b** within macrophages infected with bacteria bearing modular RNA or ds_Mango II for 1 and 1.5 hour (Fig. 6 and 7). In the case of modular RNA, signal movement was observed, supposedly indicating MTS1338 secretion into the cytoplasm of macrophages, while the presence of a tag lacking this RNA was only accompanied by the signal disappearance over time.

In conclusion, a new dye with improved selectivity and red fluorescence has been developed and shown to be promising in combination with stable in living cells Mango II aptamer for RNA imaging/tracking and FRET system development. The shortest fluorescence-activating tag for live-cell imaging to date has been proposed and its effectiveness has been tested for tracking the movement of a fluorescent signal from labeled bacterial RNA in a complex living system consisting of macrophages infected with *M. smegmatis*, a surrogate for *M. tuberculosis*. Thus, our approach can find applications in developing genetically encodable molecular tools for studying RNA-RNA interactions and conformational rearrangements, as well as sensors and advanced devices.

## Supporting information

Methods, Figures and Tables, NMR and HRMS spectra, HPLC profiles

## ACKNOWLEDGEMENTS

This work (except molecular modeling, binding assay, and cytotoxicity assays) was supported by Russian Foundation for Basic Research (grant N<u>o</u> 20-34-70143 to AVA), Russian Science Foundation (grant N<u>o</u> 22-15-00129 to AMV, molecular modeling; grant N<u>o</u> 22-14-00235 to TLA, cytotoxicity assays) and the Ministry of Science and Higher Education of the Russian Federation (grant 075-15-2019-1669 to AMV, binding assay).

## AUTHOR CONTRIBUTIONS

A.V.A. conceived of the study, and directed the project. O.S.B., V.B.T., T.S.Z., T.L.A., A.M.V. and A.V.A. designed the experiments, analyzed the data and wrote the manuscript. A.A.K., P.N.K., B.S.T., I.A.I., E.S.B. and A.V.A. synthesized and characterized compounds. T.S.Z. synthesized and purified oligonucleotides. J.I.S. and A.M.V. studied the spectral properties of dyes *in a cuvette*. V.B.T. performed molecular modeling. O.S.B., Y.V.S., L.V.A., Y.M.K. and L.I.K. performed cloning and cytotoxicity assays. O.S.B. performed the cellular imaging of the small non-coding RNA.

## ADDITIONAL INFORMATION

### Supplementary Information

accompanies this paper at https://doi.org/

### Reprints and permission

information is available online at http://npg.nature.com/reprintsandpermissions/

### Publisher’s note

Springer Nature remains neutral with regard to jurisdictional claims in published maps and institutional affiliations.

## REFERENCES

1. Romano G, Veneziano D, Acunzo M, Croce CM. Small non-coding RNA and cancer. Carcinogenesis 38, 485–491 (2017).

2. Zhang Z, Zhang J, Diao L, Han L. Small non-coding RNAs in human cancer: function, clinical utility, and characterization. Oncogene 40, 1570–1577 (2021).

3. Weinberg MS, Wood MJ. Short non-coding RNA biology and neurodegenerative disorders: novel disease targets and therapeutics. Hum Mol Genet 18, R27–39 (2009).

4. Wu YY, Kuo HC. Functional roles and networks of non-coding RNAs in the pathogenesis of neurodegenerative diseases. J Biomed Sci 27, 49 (2020).

5. Waters LS, Storz G. Regulatory RNAs in bacteria. Cell 136, 615–628 (2009).

6. Ahmed W, Zheng K, Liu ZF. Small Non-Coding RNAs: New Insights in Modulation of Host Immune Response by Intracellular Bacterial Pathogens. Front Immunol 7, 431 (2016).

7. Armitage BA. Imaging of RNA in live cells. Curr Opin Chem Biol 15, 806–812 (2011).

8. Braselmann E, Rathbun C, Richards EM, Palmer AE. Illuminating RNA Biology: Tools for Imaging RNA in Live Mammalian Cells. Cell Chem Biol 27, 891–903 (2020).

9. Trachman RJ, Ferre-D’Amare AR. Tracking RNA with light: selection, structure, and design of fluorescence turn-on RNA aptamers. Q Rev Biophys 52, e8 (2019).

10. Paige JS, Wu KY, Jaffrey SR. RNA mimics of green fluorescent protein. Science 333, 642–646 (2011).

11. Filonov GS, Moon JD, Svensen N, Jaffrey SR. Broccoli: rapid selection of an RNA mimic of green fluorescent protein by fluorescence-based selection and directed evolution. J Am Chem Soc 136, 16299–16308 (2014).

12. Song W, et al. Imaging RNA polymerase III transcription using a photostable RNA-fluorophore complex. Nat Chem Biol 13, 1187–1194 (2017).

13. Steinmetzger C, Palanisamy N, Gore KR, Hobartner C. A Multicolor Large Stokes Shift Fluorogen-Activating RNA Aptamer with Cationic Chromophores. Chemistry 25, 1931–1935 (2019).

14. Gotrik M, Sekhon G, Saurabh S, Nakamoto M, Eisenstein M, Soh HT. Direct Selection of Fluorescence-Enhancing RNA Aptamers. J Am Chem Soc 140, 3583–3591 (2018).

15. Braselmann E, et al. A multicolor riboswitch-based platform for imaging of RNA in live mammalian cells. Nat Chem Biol 14, 964–971 (2018).

16. Chen X, et al. Visualizing RNA dynamics in live cells with bright and stable fluorescent RNAs. Nat Biotechnol 37, 1287–1293 (2019).

17. Bouhedda F, et al. A dimerization-based fluorogenic dye-aptamer module for RNA imaging in live cells. Nat Chem Biol 16, 69–76 (2020).

18. Kong KYS, Jeng SCY, Rayyan B, Unrau PJ. RNA Peach and Mango: Orthogonal two-color fluorogenic aptamers distinguish nearly identical ligands. RNA, (2021).

19. Autour A, et al. Fluorogenic RNA Mango aptamers for imaging small non-coding RNAs in mammalian cells. Nat Commun 9, 656 (2018).

20. Dolgosheina EV, et al. RNA mango aptamer-fluorophore: a bright, high-affinity complex for RNA labeling and tracking. ACS Chem Biol 9, 2412–2420 (2014).

21. Jepsen MDE, et al. Development of a genetically encodable FRET system using fluorescent RNA aptamers. Nat Commun 9, 18 (2018).

22. Kolb HC, Finn MG, Sharpless KB. Click Chemistry: Diverse Chemical Function from a Few Good Reactions. Angew Chem Int Ed Engl 40, 2004–2021 (2001).

23. Rostovtsev VV, Green LG, Fokin VV, Sharpless KB. A stepwise huisgen cycloaddition process: copper(I)-catalyzed regioselective “ligation” of azides and terminal alkynes. Angew Chem Int Ed Engl 41, 2596–2599 (2002).

24. Tornoe CW, Christensen C, Meldal M. Peptidotriazoles on solid phase: [1,2,3]-triazoles by regiospecific copper(i)-catalyzed 1,3-dipolar cycloadditions of terminal alkynes to azides. J Org Chem 67, 3057–3064 (2002).

25. Pasini D. The click reaction as an efficient tool for the construction of macrocyclic structures. Molecules 18, 9512–9530 (2013).

26. Sahu A, Das D, Agrawal RK, Gajbhiye A. Bio-isosteric replacement of amide group with 1,2,3-triazole in phenacetin improves the toxicology and efficacy of phenacetin-triazole conjugates (PhTCs). Life Sci 228, 176–188 (2019).

27. Bi F, Ji S, Venter H, Liu J, Semple SJ, Ma S. Substitution of terminal amide with 1H-1,2,3-triazole: Identification of unexpected class of potent antibacterial agents. Bioorg Med Chem Lett 28, 884–891 (2018).

28. Mohammed I, et al. 1,2,3-Triazoles as Amide Bioisosteres: Discovery of a New Class of Potent HIV-1 Vif Antagonists. J Med Chem 59, 7677–7682 (2016).

29. Agouram N, El Hadrami EM, Bentama A. 1,2,3-Triazoles as Biomimetics in Peptide Science. Molecules 26, (2021).

30. Doak BC, Scanlon MJ, Simpson JS. Synthesis of unsymmetrical 1,1 ‘-disubstituted bis(1,2,3-triazole)s using monosilylbutadiynes. Org Lett 13, 537–539 (2011).

31. Sun XL, Stabler CL, Cazalis CS, Chaikof EL. Carbohydrate and protein immobilization onto solid surfaces by sequential Diels-Alder and azide-alkyne cycloadditions. Bioconjug Chem 17, 52–57 (2006).

32. Bureš F. Fundamental aspects of property tuning in push–pull molecules. RSC Adv 4, 58826–58851 (2014).

33. Dai P, et al. Influence of the Terminal Electron Donor in D-D-pi-A Organic Dye-Sensitized Solar Cells: Dithieno[3,2-b:2’,3’-d]pyrrole versus Bis(amine). ACS Appl Mater Interfaces 7, 22436–22447 (2015).

34. Kovalska VB, Kryvorotenko DV, Balanda AO, Losytskyy MYu, Tokar VP, Yarmoluk SM. Fluorescent homodimer styrylcyanines: synthesis and spectral-luminescent studies in nucleic acids and protein complexes. Dyes Pigm 67, 47–54 (2005).

35. Trachman RJ, 3rd, et al. Crystal Structures of the Mango-II RNA Aptamer Reveal Heterogeneous Fluorophore Binding and Guide Engineering of Variants with Improved Selectivity and Brightness. Biochemistry 57, 3544–3548 (2018).

36. Cawte AD, Unrau PJ, Rueda DS. Live cell imaging of single RNA molecules with fluorogenic Mango II arrays. Nat Commun 11, 1283 (2020).

37. Trachman RJ, 3rd, et al. Structure-Guided Engineering of the Homodimeric Mango-IV Fluorescence Turnon Aptamer Yields an RNA FRET Pair. Structure 28, 776–785 e773 (2020).

38. Miglietta G, et al. RNA G-Quadruplexes in Kirsten Ras (KRAS) Oncogene as Targets for Small Molecules Inhibiting Translation. J Med Chem 60, 9448–9461 (2017).

39. Malina J, Scott P, Brabec V. Stabilization of human telomeric RNA G-quadruplex by the water-compatible optically pure and biologically-active metallohelices. Sci Rep 10, 14543 (2020).

40. Yang P, De Cian A, Teulade-Fichou MP, Mergny JL, Monchaud D. Engineering bisquinolinium/thiazole orange conjugates for fluorescent sensing of G-quadruplex DNA. Angew Chem Int Ed Engl 48, 2188–2191 (2009).

41. Luu KN, Phan AT, Kuryavyi V, Lacroix L, Patel DJ. Structure of the human telomere in K+ solution: an intramolecular (3 + 1) G-quadruplex scaffold. J Am Chem Soc 128, 9963–9970 (2006).

42. Dai J, Carver M, Punchihewa C, Jones RA, Yang D. Structure of the Hybrid-2 type intramolecular human telomeric G-quadruplex in K+ solution: insights into structure polymorphism of the human telomeric sequence. Nucleic Acids Res 35, 4927–4940 (2007).

43. Marchand A, Gabelica V. Folding and misfolding pathways of G-quadruplex DNA. Nucleic Acids Res 44, 10999–11012 (2016).

44. Lim KW, et al. Sequence variant (CTAGGG)n in the human telomere favors a G-quadruplex structure containing a G.C.G.C tetrad. Nucleic Acids Res 37, 6239–6248 (2009).

45. Kovanda A, Zalar M, Sket P, Plavec J, Rogelj B. Anti-sense DNA d(GGCCCC)n expansions in C9ORF72 form i-motifs and protonated hairpins. Sci Rep 5, 17944 (2015).

46. Bychenko OS, Skvortsova YV, Grigorov AS, Azhikina TL. Use of Genetically Encoded Fluorescent Aptamers for Visualization of Mycobacterium tuberculosis Small RNA MTS1338 in Infected Macrophages. Dokl Biochem Biophys 493, 185–189 (2020).

47. Pita T, Feliciano JR, Leitao JH. Extracellular RNAs in Bacterial Infections: From Emerging Key Players on Host-Pathogen Interactions to Exploitable Biomarkers and Therapeutic Targets. Int J Mol Sci 21, (2020).

48. Chaturvedi V, Dwivedi N, Tripathi RP, Sinha S. Evaluation of Mycobacterium smegmatis as a possible surrogate screen for selecting molecules active against multi-drug resistant Mycobacterium tuberculosis. J Gen Appl Microbiol 53, 333–337 (2007).

49. Bychenko O, et al. Mycobacterium tuberculosis Small RNA MTS1338 Confers Pathogenic Properties to Non-Pathogenic Mycobacterium smegmatis. Microorganisms 9, (2021).

50. Strack R. A peck of Peppers. Nat Methods 16, 1075 (2019).

51. Trachman RJ, 3rd, et al. Structure and functional reselection of the Mango-III fluorogenic RNA aptamer. Nat Chem Biol 15, 472–479 (2019).

52. Weston SA, Parish CR. New fluorescent dyes for lymphocyte migration studies. Analysis by flow cytometry and fluorescence microscopy. J Immunol Methods 133, 87–97 (1990).

53. Lee LG, Chen CH, Chiu LA. Thiazole orange: a new dye for reticulocyte analysis. Cytometry 7, 508–517 (1986).

54. Ishiguro T, Saitoh J, Yawata H, Otsuka M, Inoue T, Sugiura Y. Fluorescence detection of specific sequence of nucleic acids by oxazole yellow-linked oligonucleotides. Homogeneous quantitative monitoring of in vitro transcription. Nucleic Acids Res 24, 4992–4997 (1996).

55. Rye HS, et al. Stable fluorescent complexes of double-stranded DNA with bis-intercalating asymmetric cyanine dyes: properties and applications. Nucleic Acids Res 20, 2803–2812 (1992).

56. Filonov GS, Kam CW, Song W, Jaffrey SR. In-gel imaging of RNA processing using broccoli reveals optimal aptamer expression strategies. Chem Biol 22, 649–660 (2015).

57. Gu H, et al. Salmonella produce microRNA-like RNA fragment Sal-1 in the infected cells to facilitate intracellular survival. Sci Rep 7, 2392 (2017).

58. Pagliuso A, et al. An RNA-Binding Protein Secreted by a Bacterial Pathogen Modulates RIG-I Signaling. Cell Host Microbe 26, 823–835 e811 (2019).

59. Salina EG, et al. MTS1338, A Small Mycobacterium tuberculosis RNA, Regulates Transcriptional Shifts Consistent With Bacterial Adaptation for Entering Into Dormancy and Survival Within Host Macrophages. Front Cell Infect Microbiol 9, 405 (2019).

