## Supplementary material for "Red light-emitting short Mango-based system enables tracking a mycobacterial small noncoding RNA in infected macrophages": Methods, Figures and Tables, NMR and HRMS spectra, HPLC profiles

### Chemistry

#### General

All reagents and solvents were commercially available unless otherwise mentioned. Thin layer chromatography (TLC) was performed on plates (Merck) precoated with silica gel (60 μm, F254) and visualized using UV light (254 and 365 nm). Column chromatography (CC) was performed on silica gel (0.040–0.063 mm, Merck, Germany). $^{1}H$ and $^{13}C$ NMR spectra were recorded on the Bruker Avance III 600 spectrometer at 600 and 151 MHz, respectively. Chemical shifts are reported in δ (ppm) units using residual 1H signals from deuterated solvents as references. The multiplicity are reported using the following abbreviations: s (singlet), d (doublet), t (triplet), m (multiplet) and br (broad). The coupling constants (J) are given in Hz. ESI HR mass spectra were acquired on a Thermo Scientific LTQ Orbitrap hybrid instrument (Thermo Electron Corp., Bremen, Germany) in continuous flow direct sample infusion (positive ion mode). 2-Methyl-3-(prop-2-yn-1-yl)benzo[d]thiazol-3-ium bromide **1** was prepared according to the reported method[1].

#### Preparative HPLC purification of TO1-biotin and its analogs 4a-d

Purification of the compounds was performed on a Gilson HPLC system (331/332 pump with GX-271 liquid handler) using an Luna C18(2) (100×21.20 mm, 5 μm) column. MeCN (with 0.1% TFA) and aq. TFA (0.1%) were used as eluents. UV detection was achieved at 210 nm and 280 nm. TO1-biotin and **4a-d** were purified in a linear gradient from 20 to 70% of MeCN in 45 min at a flow rate of 25 mL/min. Fractions containing the target compound were collected, organic solvent was removed *in vacuo* and water was lyophilized.

#### LC-HRMS analysis

All experiments were carried out on a Dionex 3000 UltiMate HPLC system coupled with QExactive Orbitrap mass spectrometer (Bremen, Germany) using Macherey-Nagel Pyramid C18

Nucleodur (2.0×100 mm, 1.8 μm) HPLC column. Mobile phase A: 0.1% formic acid in 5% aqueous solution of acetonitrile; mobile phase B: 0.1% formic acid in acetonitrile. The following gradient at a flow rate of 0.50 mL/min was used: 0-1.0 min 5% B, 1.0-10.0 min 5-95.0% B, 10.0-12.0 95% B, 12.0-13.0 min 95.0-5% B, 13.0-15.0 min 5% B. The injection volume was 1 μL. The resolving power was 35 000 (for m/z = 200). The Sheath, Aux and Spare gases were set to 45, 35 and 5, respectively. The Spray Voltage was 4.1 kV, the temperature of the desolvating capillary was 350 °C. S-Leans RF level was 50 and the source temperature was set to 200 °C. All mass spectra were obtained in Full MS followed by data dependent analysis (DDa) in a positive ionization mode. The retention time are shown in the relevant parts of the chemical procedures related to product characterization.

**Preparation of intermediates**

*(E)-2-((1-methylquinolin-4(1H)-ylidene)methyl)-3-(prop-2-yn-1-yl)benzo[d]thiazol-3-ium bromide* **2a**

To a suspension of 2-methyl-3-(prop-2-yn-1-yl)benzo[d]thiazol-3-ium bromide **1** (270 mg, 1 mmol) in $CH_2Cl_2$ (20 mL) 1-methylquinolinium bromide (270 mg, 1.2 mmol) and TEA (0.7 mL, 5 mmol) were added and the resulting mixture was stirred at rt for 48 hours. The mixture was concentrated *in vacuo*. The solid obtained was triturated with $Et_2O$ (3 x 15 mL) and then purified by column chromatography on silica gel (1→10% $CH_3OH$ in $CH_2Cl_2$) yielding **2a** (176 mg, 0.43 mmol, 43 %) as a red solid. $^1H$ NMR (600 MHz, DMSO-d6): δ 8.81 (d, J = 8.5 Hz, 1H), 8.78 (d, J = 7.1 Hz, 1H), 8.13 (d, J = 8.7 Hz, 1H), 8.06 (d, J = 7.1 Hz, 1H), 8.04 (d, J = 8.2 Hz, 1H), 7.85 (t, J = 7.7 Hz, 1H), 7.79 (d, J = 8.3 Hz, 1H), 7.61 (t, J = 7.8 Hz, 1H), 7.46 (d, J = 7.1 Hz, 1H), 7.41 (t, J = 7.6 Hz, 1H), 7.05 (s, 1H), 5.59 (d, J = 1.9 Hz, 2H), 4.24 (s, 3H), 3.53 (t, J = 1.9 Hz, 1H). $^{13}C$ NMR (151 MHz, DMSO-d6): δ 158.18, 149.00, 145.54, 139.12, 137.96, 133.32, 128.08, 127.18, 125.29, 124.44, 124.19, 123.36, 122.93, 118.40, 112.44, 108.65, 88.02, 76.82, 76.31, 42.55, 35.32. HRMS (ESI) m/z: calcd for $C_{21}H_{17}N_2S^+$ $[M-Br]^+$: 329.1107; found 329.1118; $t_R$ = 4.01 min.

*(E)-2-(2-(4-(Dimethylamino)phenyl)ethenyl)-3-(2-propynyl)benzo[d]thiazolium bromide* **2b**

This derivative (yield 79%) was prepared according to the reported method[2]. $^1H$ NMR (600 MHz, DMSO-$d_6$): δ 8.32 (d, J = 8.0 Hz, 1H), 8.14 (d, J =15.0 Hz, 1H), 8.13 (d, J =8.3 Hz, 1H), 7.95 (d, J =8.7 Hz, 2H), 7.80 (t, J =7.7 Hz, 1H), 7.74 (d, J =15.0 Hz, 1H), 7.68 (t, J =7.6 Hz, 1H), 6.86 (d, J = 8.7 Hz, 2H), 5.82 (s, 2H), 3.72 (t, J = 2.0 Hz, 1H), 3.13 (s, 6H). $^{13}C$ NMR (151 MHz, DMSO-d6): δ 171.34, 153.83, 151.56, 140.31, 133.30 (2C), 128.92, 127.40, 126.57, 123.99, 121.43, 115.50, 112.01, 105.18, 78.01, 75.96, 37.48. HRMS (ESI) m/z: calcd for $C_{20}H_{19}N_2S^+$ $[M-Br]^+$: 319.1263; found 319.1255; $t_R$ = 4.11 min.

*(E)-3-(prop-2-yn-1-yl)-2-(2-(2,3,6,7-tetrahydro-1H,5H-pyrido[3,2,1-ij]quinolin-9-yl)vinyl)benzo[d]thiazol-3-ium bromide* **2c**

This derivative (yield 74%) was prepared according to the reported method[2]. $^1H$ NMR (600 MHz, DMSO-d6): δ 8.22 (d, J = 8.0 Hz, 1H), 8.02 (d, J = 8.5 Hz, 1H), 7.93 (d, J = 14.8 Hz, 1H), 7.75 (t, J = 7.7 Hz, 1H), 7.63 (t, J = 7.8 Hz, 1H), 7.52 (s, 2H), 7.48 (d, J = 7.8 Hz, 1H), 5.66 (d, J = 2.2 Hz, 2H), 3.59 (t, J = 2.2 Hz, 1H), 3.44-3.39 (m, 4H), 2.78-2.74 (m, 4H), 1.96-1.90 (m, 4H). HRMS (ESI) m/z: calcd for $C_{24}H_{23}N_2S^+$ $[M-Br]^+$: 371.1576; found 371.1593; $t_R$ = 4.81 min.

*(E)-3-(prop-2-yn-1-yl)-2-(2-(1,2,2,4-tetramethyl-1,2-dihydroquinolin-6-yl)vinyl)benzo[d]thiazol-3-ium bromide* **2d**

A mixture of 2-methyl-3-(prop-2-yn-1-yl)benzo[d]thiazol-3-ium bromide **1** (270 mg, 1 mmol) and 1,2,2,4-tetramethyl-1,2-dihydroquinoline-6-carbaldehyde (430 mg, 2 mmol) in acetic anhydride (3 mL) was stirred at 90°C for 8 h. Then, to a mixture $H_2O$ (0.5 mL) was added and the stirring was continued for 30 min followed by concentration *in vacuo*. The residue was purified by column chromatography on silica gel (1→10% $CH_3OH$ in $CH_2Cl_2$) yielding **2d** (330 mg, 0.71 mmol, 71 %) as a purple solid. $^1$H NMR (600 MHz, DMSO-d6): δ 8.30 (d, J = 8.0 Hz, 1H), 8.13 (d, J = 15.0 Hz, 1H), 8.11 (d, J = 8.7 Hz, 1H), 7.86 (d, J = 8.9 Hz, 1H), 7.80 (t, J = 7.9 Hz, 1H), 7.68 (t, J = 7.7 Hz, 1H), 7.64 (d, J = 15.0 Hz, 1H), 7.61 (d, J = 1.6 Hz, 1H), 6.72 (d, J = 8.9 Hz, 1H), 5.79 (d, J = 2.2 Hz, 2H), 5.53 (s, 1H), 3.71 (t, J = 2.2 Hz, 1H), 2.99 (s, 3H), 2.04 (s, 3H), 1.40 (s, 6H). $^{13}$C NMR (151 MHz, DMSO-d6): δ 170.98, 151.54, 149.82, 140.34, 134.34, 130.15, 128.89, 127.32, 126.45, 125.72 (2C), 123.92, 121.51, 121.49, 115.41, 110.74, 104.62, 78.03, 75.99, 58.14, 37.40, 31.48, 28.55, 18.19. HRMS (ESI) m/z: calcd for $C_{25}H_{25}N_2S^+$ [M-Br]$^+$: 385.1733; found 385.1746; $t_R$ = 5.08 min.

*N-(2-(2-(2-(2-azidoethoxy)ethoxy)ethoxy)ethyl)-5-(2-oxohexahydro-1H-thieno[3,4-d]imidazol-4-yl)pentanamide* **3**

This derivative (83%) was prepared starting from 2-(2-(2-(2-azidoethoxy)ethoxy)ethoxy)ethan-1-amine according to the reported method[3]. $^1$H NMR (600 MHz, DMSO-d6): δ 7.80 (t, J = 5.5 Hz, 1H), 6.39 (s, 1H), 6.34 (s, 1H), 4.33-4.29 (m, 1H), 4.15-4.11 (m, 1H), 3.62-3.59 (m, 2H), 3.58-3.49 (m, 8H), 3.42-3.38 (m, 4H), 3.21-3.17 (m, 2H), 3.12-3.08 (m, 1H), 2.83 (dd, J = 12.4 Hz, J = 5.1 Hz, 1H), 2.59 (d, J = 12.4 Hz, 1H), 2.07 (t, J = 7.4 Hz, 2H), 1.65-1.58 (m, 1H), 1.55-1.44 (m, 3H), 1.36-1.24 (m, 2H). $^{13}$C NMR (151 MHz, DMSO-d6): δ 172.1, 162.6, 69.7, 69.7, 69.6, 69.4, 69.1, 69.0, 61.0, 59.1, 55.3, 49.9, 39.7 (overlaps with DMSO), 38.4, 35.0, 28.1, 27.9, 25.1.

TO1-biotin (yield 6.8%) was prepared according to the reported condensation method[4]. HRMS (ESI) m/z: calcd for $C_{38}H_{49}N_6O_6S_2^+$ [M-$CF_3COO^-$]$^+$: 749.3150; found 749.3143; $t_R$ = 7.42 min.

**General procedure for copper catalyzed azide−alkyne cycloaddition (preparation of final compounds)**

A solution of the corresponding alkyne-containing derivative **2** (0.06 mmol) and azido-containing biotinylated derivative **3** (40 mg, 0.09 mmol, 1.5 eq) in a mixture of $CH_3OH/CH_3CN$ (1:1, v/v, 5 mL) was degassed and then copper (I) iodide (1.1 mg, 0.006 mmol, 0.1 eq), TBTA (1.6 mg, 0.003 mmol, 0.05 eq) and DIPEA (55 μL, 0.3 mmol, 5.0 eq) were sequentially added under a nitrogen atmosphere. The reaction mixture was stirred at room temperature overnight and concentrated *in vacuo*. The residue was partitioned between a mixture of *n*-butanol/$CH_2Cl_2$ (1:10, v/v, 10 mL) and 0.1 M EDTA solution (10 mL) and the aqueous phase was additionally extracted with *n*-butanol/$CH_2Cl_2$ (1:10, v/v, 10 mL). The combined organic layers was concentrated *in vacuo* and the residue obtained was subjected to preparative HPLC.

*2-((-1-methylquinolin-4(1H)-ylidene)methyl)-3-((1-(13-oxo-17-((3aR,4R,6aS)-2-oxohexahydro-1H-thieno[3,4-d]imidazol-4-yl)-3,6,9-trioxa-12-azaheptadecyl)-1H-1,2,3-triazol-4-yl)methyl)benzo[d]thiazol-3-ium 2,2,2-trifluoroacetate* **4a**

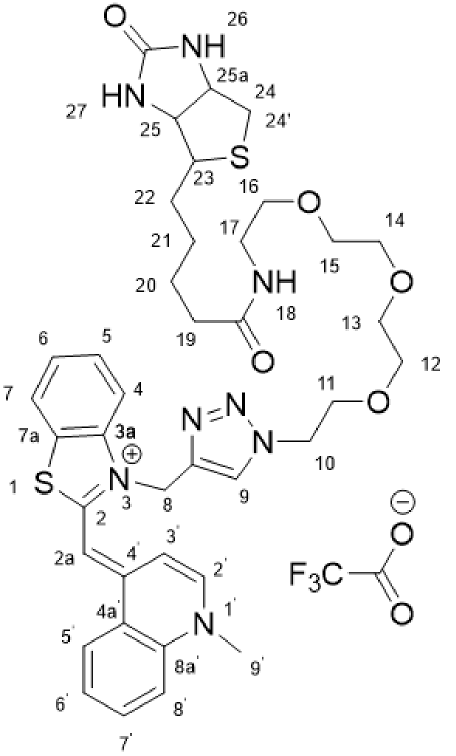

(Yield 58.1 %). $^{1}$H NMR (600 MHz,DMSO-$d_6$): δ 8.78 (1H, d, *J* = 8.6 Hz, **H5'**), 8.67 (1H, d, *J* = 7.3 Hz, **H2'**), 8.31 (1H, s, **H9**), 8.13 – 8.10 (1H, m, **H8'**), 8.07 – 8.03 (2H, m, **H7'**, **H7**), 7.96 (1H, d, *J* = 8.3 Hz, **H4**), 7.86 – 7.82 (1H, m, **H6'**), 7.80 – 7.75 (1H, m, **H18**), 7.65 – 7.61 (1H, m, **H5**), 7.45 – 7.40 (2H, m, **H3'**, **H6**), 7.35 (1H, s, **H2a**), 6.38 (1H, s, **H26**), 6.34 (1H, s, **H27**), 5.93 (2H, s, **H8**), 4.51 (2H, t, *J* = 5.1 Hz, **H10**), 4.32 – 4.27 (1H, m, **H25a**), 4.22 (3H, s, **H9'**), 4.13 – 4.09 (1H, m, **H25**), 3.77 (2H, t, *J* = 5.1 Hz, **H11**), 3.45 – 3.37 (10H, m, **H12-H16**), 3.19 – 3.13 (2H, m, **H17**), 3.10 – 3.05 (1H, m, **H23**), 2.81 (1H, dd, *J* = 12.4, 5.1 Hz, **H24**), 2.57 (1H, d, *J* = 12.4 Hz, **H24'**), 2.05 (2H, t, *J* = 7.5 Hz, **H19**), 1.54 – 1.39 (6H, m, **H20-H22**). HRMS (ESI) m/z: calcd for $C_{39}H_{49}N_8O_5S_2^+$ $[M-CF_3COO^-]^+$: 773.3262; found 773.3260; $t_R$ = 7.91 min.

*2-(4-(dimethylamino)styryl)-3-((1-(13-oxo-17-((3aR,4R,6aS)-2-oxohexahydro-1H-thieno[3,4-d]imidazol-4-yl)-3,6,9-trioxa-12-azaheptadecyl)-1H-1,2,3-triazol-4-yl)methyl)benzo[d]thiazol-3-ium 2,2,2-trifluoroacetate* **4b**

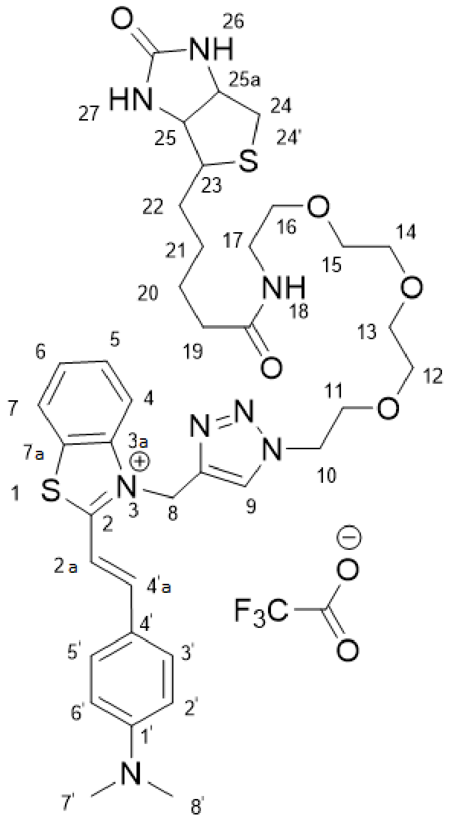

(Yield 37.4 %). $^{1}$H NMR (600 MHz, DMSO-$d_6$): δ 8,34 (1H, s, **H9**), 8.29 (1H, d, $J$ = 8.0 Hz, **H7**), 8.24 (1H, d, $J$ = 8.9 Hz, **H4**), 8.16 (1H, d, $J$ = 14.6 Hz, **H2a**), 7.93 (2H, d, $J$ = 9.0 Hz, **H2**', **H6**'), 7.89 (1H, d, $J$ = 15.2 Hz, **H4'a**), 7.80 – 7.75 (2H, m, **H5**, **H18**), 7.69 – 7.65 (1H, m, **H6**), 6.89 (2H, d, $J$ = 9.0 Hz, **H3**', **H5**'), 6.38 (1H, s, **H25**), 6.34 (1H, s, **H26**), 6.17 (2H, s, **H8**), 4.52 (2H, t, $J$ = 5.1 Hz, **H10**), 4.32 – 4.28 (1H, m, **H25a**), 4.14 – 4,10 (1H, m, **H25**), 3.77 (2H, t, $J$ = 5.1 Hz, **H11**), 3.42 – 3.37 (10H, m, **H12-H16**), 3.19 – 3.15 (2H, m, **H17**), 3.14 (6H, s, **H7**', **H8**'), 3.10-3.06 (1H, m, **H23**), 2.81 (1H, dd, $J$ = 12.4, 5.1 Hz, **H24**), 2.59 – 2.56 (1H, m, **H24**'), 2.05 (2H, t, $J$ = 7.5 Hz, **H19**), 1.64 – 1.40 (6H, m, **H20-H22**). HRMS (ESI) m/z: calcd for $C_{38}H_{51}N_8O_5S_2^+$ [M-$CF_3COO^-$]$^+$: 763.3418; found 763.3403; $t_R$ = 8.27 min.

*3-((1-(13-oxo-17-((3aR,4R,6aS)-2-oxohexahydro-1H-thieno[3,4-d]imidazol-4-yl)-3,6,9-trioxa-12-azaheptadecyl)-1H-1,2,3-triazol-4-yl)methyl)-2-(2-(2,3,6,7-tetrahydro-1H,5H-pyrido[3,2,1-ij]quinolin-9-yl)vinyl)benzo[d]thiazol-3-ium 2,2,2-trifluoroacetate* **4c**

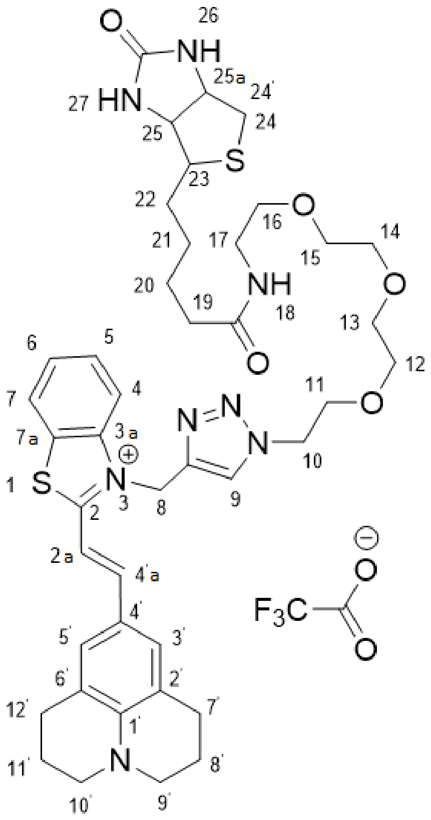

(Yield 40.4 %) $^{1}$H NMR (600 MHz, DMSO-$d_6$): δ 8.32 (1H, s, **H9**), 8.23 (1H, d, *J* = 8.0 Hz, **H7**), 8.12 (1H, d, *J* = 8.4 Hz, **H4**), 7.95 (1H, d, *J* = 14.8 Hz, **H2a**), 7.81 – 7.76 (1H, m, **H18**) 7.73 – 7.68 (2H, m, **H5, H4'a**), 7.63 – 7.59 (1H, m, **H6**), 7.52 (2H, s, **H3'**, **H5'**), 6.38 (1H, s, **H26**), 6.34 (1H, s, **H27**), 6.08 (2H, s, **H8**), 4.52 (2H, t, *J* = 5.1 Hz, **H10**), 4.32 – 4.28 (1H, m, **H25a**), 4.14 – 4.10 (1H, m, **H25**), 3.78 (2H, t, *J* = 5.1 Hz, **H11**), 3.46 – 3.35 (14H, m, **H12-H16, H10'**, **H9'**), 3.21 – 3.13 (2H, m, **H17**), 3.12 – 3.05 (1H, m, **H23**), 2.81 (1H, dd, *J* = 12.4, 5.1 Hz, **H24**), 2.74 (4H, t, *J* = 6.3 Hz, **H7'**, **H12'**), 2.58 (1H, d, *J* = 12.4 Hz, **H24'**), 2.05 (2H, t, *J* = 7.4 Hz, **H19**), 1.94 – 1.88 (4H, m, **H11'**, **H8'**), 1.65 – 1.40 (6H, m, **H20-H22**). HRMS (ESI) m/z: calcd for $C_{42}H_{55}N_8O_5S_2^+$ [M-$CF_3COO^-$]$^+$: 815.3731; found 815.3768; $t_R$ = 4.46 min.

*3-((1-(13-oxo-17-((3aR,4R,6aS)-2-oxohexahydro-1H-thieno[3,4-d]imidazol-4-yl)-3,6,9-trioxa-12-azaheptadecyl)-1H-1,2,3-triazol-4-yl)methyl)-2-(2-(1,2,2,4-tetramethyl-1,2-dihydroquinolin-6-yl)vinyl)benzo[d]thiazol-3-ium 2,2,2-trifluoroacetate* **4d**

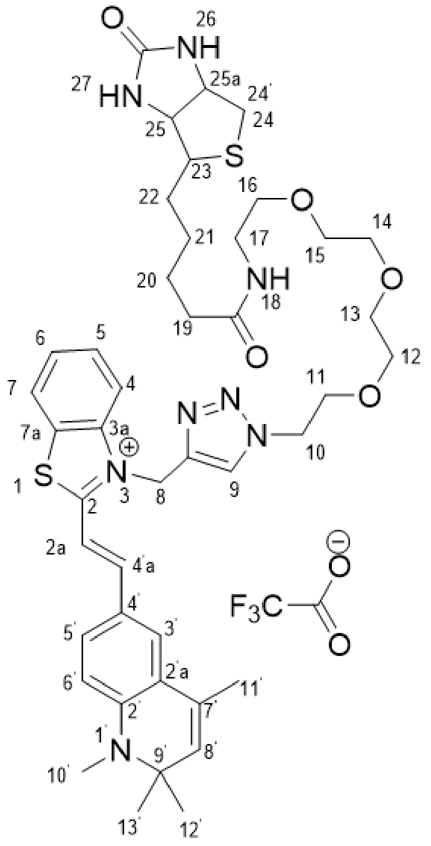

(Yield 41.9 %) $^{1}$H NMR (600 MHz, DMSO-$d_6$): δ 8.32 (1H, s, **H9**), 8.28 (1H, d, *J* = 8.8 Hz, **H7**), 8.21 (1H, d, *J* = 8.7 Hz, **H4**), 8.11 (1H, d, *J* = 15.0 Hz, **H2a**), 7.86 – 7.74 (4H, m, **H4'a, H6', H18, H5**), 7.68 – 7.64 (1H, m, **H6**), 7.61 – 7.60 (2H, m, **H3'**, **H5'**), 6.71 (1H, d, *J* = 9.0 Hz, **H8'**), 6.38 (1H, s, **H26**), 6.35 (1H, s, **H27**), 6.17 (2H, s, **H8**), 4.52 (2H, t, *J* = 5.1 Hz, **H10**), 4.32 – 4.28 (1H, m, **H25a**), 4.13 – 4.09 (1H, m, **H25**), 3.77 (2H, t, *J* = 5.0 Hz, **H11**), 3.46 – 3.35 (10H, m, **H12-H16**), 3.19 – 3.15 (2H, m, **H17**), 3.12 – 3.06 (1H, m, **H23**), 2.98 (3H, s, **H10'**), 2.81 (1H, dd, *J* = 12.4, 5.2 Hz, **H24**), 2.57 – 2.54 (1H, m, **H24**'), 2.08 – 2.04 (2H, m, **H19**) 2.03 (3H, s, **H11**'), 1.51 – 1.41 (6H, m, **H20- H22**), 1.40 (6H, s, **H13', H12'**). HRMS (ESI) m/z: calcd for $C_{43}H_{57}N_8O_5S_2^+$ [M-$CF_3COO^-$]$^+$: 829.3888; found 829.3924; $t_R$ = 4.67 min.

#### Methods

**Oligonucleotide synthesis.** Oligonucleotides (ONs) (purity >95%, HPLC) were obtained from Litekh (Russia).

**Absorption spectroscopy, circular dichroism spectroscopy, fluorimetry, and binding assays.** Secondary structures of Mango-RNA were confirmed by circular dichroism (CD) spectroscopy using 5 μM solutions of free RNA or their 1:1 complexes with the dyes in the working buffer (10 mM Tris-HCl, pH 7.5, and 140 mM KCl). All samples were heated to 90°C for 5 min and snap-cooled on ice (rapid annealing) or cooled gradually to room temperature (slow annealing) prior to all measurements. The annealing scheme had no noticeable effect on the CD spectra. The spectra were registered with a Chirascan spectrophotometer (Applied Photophysics, UK) at room temperature in quartz cuvettes of 1 cm path.

Spectral properties of the new dyes and their 1:1 complexes with Mango RNA, as well as the 1:4 DFHBI complex with Broccoli RNA, were characterized using 5 μM dye solutions in the same working buffer. Complexes of 4b with genetically engineered RNA constructs were characterized using 1 μM solutions in a 20 mM sodium phosphate buffer, pH 7.2, supplemented with 0.05% Tween-20 and 140 mM KCl. Absorption, fluorescence excitation and emission spectra were registered with a Chirascan spectrophotometer equipped with an CS/SEM fluorescence accessory at room temperature. Fluorescence quantum yields $\Phi^F$ were calculated using equation (1)

$$\Phi^F = \Phi_r^F \times \frac{1 - 10^{A_r}}{1 - 10^{A}} \times \frac{F}{F_r} \times \frac{n^2}{n_r^2} \qquad (1)$$

where $\Phi^F_r$ = 0.95 is fluorescence quantum yield of the reference dye rhodamine 6G (R6G) in ethanol, $A_r$ and $F_r$ are absorbance and fluorescence of R6G in ethanol, A and F are absorbance and fluorescence of the dye-RNA complex in the working buffer, and $n_r$ and n are refractive indices of ethanol and the buffer, respectively.

For rough evaluation of Mango-4b binding constants, 10 nM solution of 4b in the working buffer supplemented with 0,05% Tween-20 was titrated with increasing concentrations of Mango RNA, and integral fluorescence was registered in microcapillaries using Monolith NT.115 device (NanoTemper, Germany) equipped with a RED/GREEN detector in RED mode at 25 ºC. Normalized fluorescence was analyzed using MO.Affinity Analysis software (NanoTemper, Germany), and Kd values were obtained by fitting the experimental data to equation (2)

$$\frac{\Delta F}{\Delta F_{max}} = \frac{[dye]+[RNA]+K_d-\sqrt{([dye]+[RBD]+K_d)^2-4\cdot[dye]\cdot[RNA])}}{2\cdot[dye]} \qquad (2)$$

where [dye] and [RNA] are total concentrations of 4b and Mango-RNA, respectively.

**Cell viability assay on murine macrophages (cell culture RAW 264.7, ATCC® TIB-71™).** Two-fold dilutions of studied compounds TO1-biotin and **4a-d** and DMSO as a negative control were prepared in cultural medium (FSBSI "Chumakov FSC R&D IBP RAS", Russia). Cell suspensions were added to the wells with compound dilutions or DMSO control (approx. $2 \times 10^4$ cells per well). The final concentration series of eight dilutions started from 50 μM. The cells were incubated at 37 °C in a $CO_2$-incubator for 24 hours and 7 days. After incubation, the cells were analyzed under a microscope and for cell viability. For this, cultural medium was substituted with resazurin solution (25 mg/mL). Cells were incubated at 37 °C in a $CO_2$-incubator for 4 h. Then 20 mL of 10% SDS was added to stop the reaction. Fluorescence was measured with Promega GloMax-Multi Detection System at $\lambda_{ex}$ 525 nm and $\lambda_{em}$ 580-640 nm. As additional controls, the same series of cells treated with compounds or DMSO dilutions but without resazurin solution to subtract the background fluorescence were used; and a medium with resazurin solution to set up a minimal value of non-reduced resazurin. All experimental procedures were performed in two replicates. Statistical analysis was performed and fluorescence curves were plotted using MS Excel 2013. The 50% cytotoxic concentration ($CC_{50}$) was calculated (compound concentration required to induce cytopathic effect in 50% of the cells in monolayer).

**Molecular modeling.** All 3D models were built using molecular graphics software package Sybyl-X software (Certara, USA). Partial charges on the ligands atoms were calculated using DFT/M06-2X[5]/6-311+g(d,p) and conductor-like polarizable continuum model (CPCM)[6], 6-31g* basis sets and Merz-Singh-Kollman scheme[7] with RESP (Restrained ElectroStatic Potential) method[8]. All quantum mechanics simulations were carried out using Gaussian 09 program[9]. To obtain the starting conformations of RNA-dye complexes, docking procedure was performed using ICM-Pro 3.9.2[10]. MD simulations were performed using Amber 20 software[11]. The influence of the solvent was simulated using OPC3[12] model of water molecules. The simulation performed by with periodical boundary conditions and a rectangular box. The buffer between the RNA-dye complex and the periodic box wall was at least 15 Å. For neutralizing of the negative charge of RNA backbone, $K^+$ ions were used. The parameters needed for interatomic energy calculation were taken from the force fields RNA.YIL[13,14] for RNA and from the general amber force field (gaff2) for the dyes. The MD simulations in production phase were carried out using constant temperature (T = 300 K) and constant pressure (p = 1 atm) over 80 ns. To control the temperature, Langevin thermostat was used with the collision frequency of 1 $ps^{-1}$. Energies were estimated by using the MM-GBSA approach. The polar contribution $E_{GB}$ was computed using the Generalized Born (GB) method and the algorithm developed by Onufriev et al. for calculating the effective Born radii[15]. The non-polar contribution to the solvation energy (Esurf), which includes solute-solvent van der Waals interactions and the free energy of cavity formation in solvent, was estimated from a solvent-accessible surface area (SASA).

**Cytotoxicity assays for *M. smegmatis.*** The cytotoxic effects of **4b** on bacteria were assessed by MSmeg cellular growth in the presence of the dye at various concentrations, since the measurements of bacterial growth rate reflect the concentration of actively growing cells[16]. Pre-cultured MSmeg strains bearing empty pAMYC vector[17] were diluted in ratio 1:100 with fresh LB medium containing chloramphenicol (34 μg/mL), Tween 80 (0.05%) and **4b** at a concentration of 400 nM, 1μM or without dye (positive control), and grown up at 37 °C while shaken at 200 rpm. Then, the optical density of the culture ($OD_{600}$) until the stationary growth phase was measured (Fig. S12).

***In vitro* transcription.** The modular RNA ds_Mango II_MTS1338, ds_Mango II tag and Broccoli aptamer were synthesized using T7 RiboMAX™ Express Large Scale RNA Production System according to the manufacturer’s recommendations (Promega, Madison, WI, USA). The DNA templates for RNA synthesis were generated by PCR using the appropriate expression plasmids (see below and [18]) and oligonucleotide primers T7MngII_F/1338_R, T7MngII_F/MngII_R, T7Broc_F/Broc_R (Table S2).

**Bacterial Strains and Growth Conditions.** *Mycobacterium smegmatis* mc(2)155 (MSmeg) obtained from the bacterial collection of the Bach Institute of Biochemistry (Research Center of Biotechnology of the Russian Academy of Sciences, Moscow, Russia) was pre-cultured for 24 h at 37 $^{o}$C on an orbital shaker (200 rpm) in 40 mL of Luria Bertani (LB) medium supplemented with 0.05% Tween-80 (to prevent cell clumping) and used as inoculums for further experiments. MSmeg recombinant strains were grown in LB supplemented with 34 μg/mL chloramphenicol.

**Construction of MSmeg Recombinant Strains.** The modular RNA and aptamer expression plasmids were constructed by inserting the ds_Mango II_MTS1338 and ds_Mango II encoding DNA fragments synthesized commercially (Evrogen , Russia) into HindIII site of pAMYC ($Cm^r$), between mycobacterial rrnB promoter of MSmeg and rrnB-T1 terminator[19]. The sequences of the genetic constructs are given in Table S2.

Plasmids were amplified in *Escherichia coli* DH5_α grown in Luria Bertani (LB) broth and LB agar supplemented with chloramphenicol (34 μg/mL). MSmeg cells were transformed by electroporation with the constructed plasmids and empty pAMYC vector (negative control), and recombinant strains were selected on LB agar containing with 34 μg/mL chloramphenicol.

**Confocal Microscopy of ds_Mango II_MTS1338, ds_Mango II in bacteria and in infected macrophages.** Recombinant MSmeg strains were grown up to the logarithmic growth phase ($OD_{600}$ 0.8). Cells were pelleted by centrifugation at 3000 x g for 15 min, washed twice with PBS and either incubated for 30 minutes in visualization buffer (10 mM Tris pH 7.5, 140 mM KCl, 300 nM Hoechst 33258, 400 nM dye) or resuspended in RPMI-1640 medium (Gibco Europe, Paisley, UK) for infection in macrophages. *M. smegmatis* transformed by pAMYC without insertion was used as a negative control strain.

For the infection RAW 264.7 cells cultured in RPMI-1640 supplemented with 10% fetal calf serum (FCS) (Gibco) were seeded in the same medium on cover glasses (18 × 18 mm Menzel Gläsercoverslips, Thermo Fisher Scientific, Schwerte, Germany) placed in 6-well culture plates (Costar, Cambridge, MA, USA). After 24 h, cells ($5\times10^4$ cells/glass) were infected with ds_Mango II_MTS1338, ds_Mango II or pAMYC strains at MOI 10:1 for 1 h. After 1 h of infection, the medium was removed, and infected macrophages washed three times with PBS and incubated for 30 minutes in visualization buffer (see above). Then the samples were analyzed using an Eclipse TE2000 confocal microscope (Nikon, Tokyo, Japan). ds_Mango II_MTS1338 and ds_Mango II in Mycobacteria were detected in a green channel (ex 488/em 590 nm) for TO1-biotin and in a red channel (ex 543/em 650 nm) for **4b**, whereas Hoechst 33258-stained cell nuclei were visualized in a blue channel (ex 408/em 515 nm).

#### Figures and Tables

```
==================================================================
Acq. Operator   : SYSTEM                          Seq. Line :   2
Sample Operator : SYSTEM
Acq. Instrument : LCMS                             Location :   P1-F-06
Injection Date  : 6/7/2022 8:20:36 PM                   Inj :   1
                                                Inj Volume : 2.000 µl
Method          : C:\Users\Public\Documents\ChemStation\1\Data\Seq1 2022-06-07 20-00-05
                  \Purity_test_SEQ.M (Sequence Method)
Last changed    : 6/7/2022 8:00:02 PM by SYSTEM
```

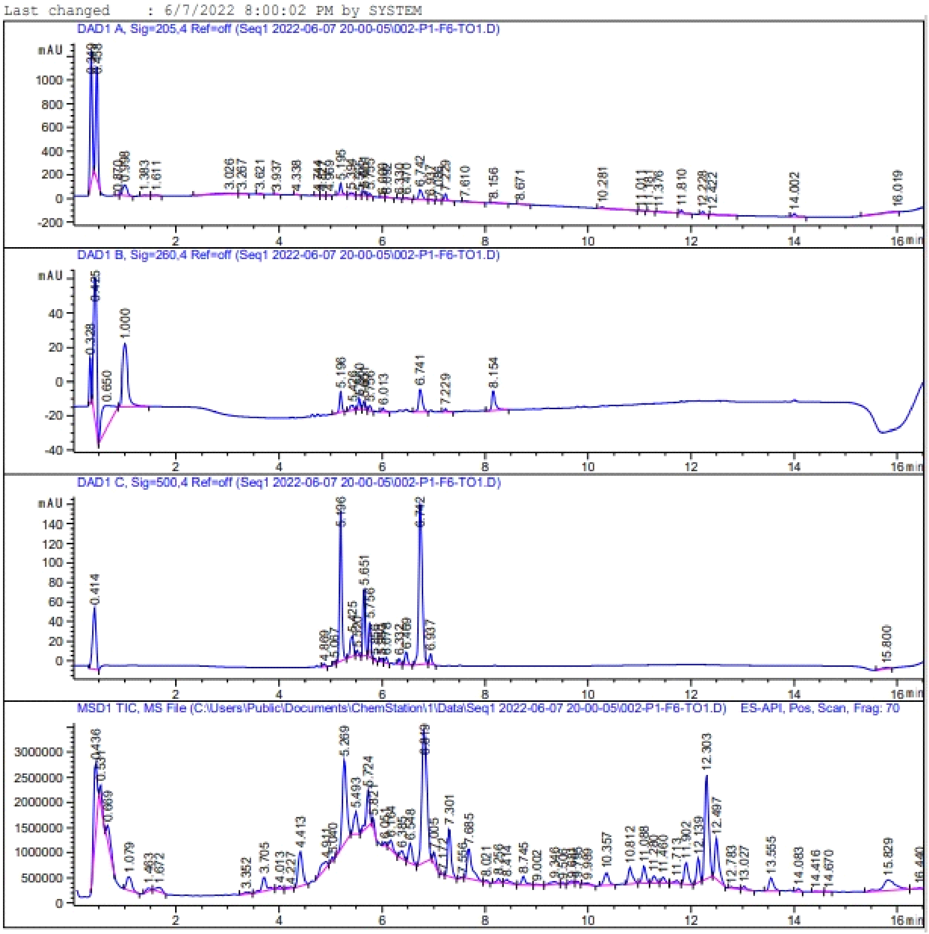

Signal 2: DAD1 B, Sig=260,4 Ref=off

| Peak # | RetTime [min] | Type | Width [min] | Area [mAU*s] | Height [mAU] | Area % |
|---|---|---|---|---|---|---|
| 1 | 0.328 | BB | 0.0362 | 54.04987 | 26.05453 | 4.8408 |
| 2 | 0.425 | BB | 0.0760 | 371.08383 | 83.26328 | 33.2350 |
| 3 | 0.650 | BB | 0.2368 | 205.10118 | 13.38522 | 18.3692 |
| 4 | 1.000 | BB | 0.1094 | 257.25781 | 36.88705 | 23.0405 |
| 5 | 5.196 | BB | 0.0481 | 37.90986 | 12.25208 | 3.3953 |
| 6 | 5.426 | BB | 0.0735 | 15.01566 | 2.84694 | 1.3448 |
| 7 | 5.560 | BB | 0.0467 | 18.16730 | 6.47644 | 1.6271 |
| 8 | 5.651 | BB | 0.0434 | 13.59577 | 5.04635 | 1.2177 |
| 9 | 5.756 | BB | 0.0473 | 9.47278 | 3.12857 | 0.8484 |
| 10 | 6.013 | BB | 0.0576 | 9.18503 | 2.34827 | 0.8226 |
| 11 | 6.741 | BB | 0.0774 | 64.77995 | 13.16849 | 5.8018 |
| 12 | 7.229 | BB | 0.0507 | 5.68122 | 1.71434 | 0.5088 |
| 13 | 8.154 | BB | 0.0703 | 55.24644 | 11.44420 | 4.9480 |
| Totals : | | | | 1116.54673 | 218.01576 | |

Signal 3: DAD1 C, Sig=500,4 Ref=off

| Peak # | RetTime [min] | Type | Width [min] | Area [mAU*s] | Height [mAU] | Area % |
|---|---|---|---|---|---|---|
| 1 | 0.414 | BB | 0.0738 | 296.36264 | 62.01221 | 13.8570 |
| 2 | 4.869 | BB | 0.0496 | 9.19292 | 3.01077 | 0.4298 |
| 3 | 5.067 | BB | 0.0405 | 7.65717 | 3.13967 | 0.3580 |
| 4 | 5.196 | BB | 0.0482 | 479.45288 | 154.35622 | 22.4177 |
| 5 | 5.425 | BB | 0.0795 | 98.80371 | 20.77542 | 4.6197 |
| 6 | 5.520 | BB | 0.0416 | 14.73562 | 5.43499 | 0.6890 |
| 7 | 5.651 | BB | 0.0453 | 195.92386 | 68.72435 | 9.1608 |
| 8 | 5.756 | BB | 0.0447 | 98.25571 | 35.02248 | 4.5941 |
| 9 | 5.856 | BB | 0.0396 | 5.11800 | 2.16438 | 0.2393 |
| 10 | 5.980 | BB | 0.0411 | 9.35109 | 3.75288 | 0.4372 |
| 11 | 6.078 | BB | 0.0505 | 17.15796 | 5.48513 | 0.8023 |
| 12 | 6.332 | BB | 0.0451 | 16.56164 | 5.84273 | 0.7744 |
| 13 | 6.469 | BB | 0.0572 | 48.85206 | 13.18771 | 2.2842 |
| 14 | 6.742 | BB | 0.0747 | 796.35474 | 163.79013 | 37.2350 |
| 15 | 6.937 | BB | 0.0482 | 32.29179 | 10.39579 | 1.5099 |
| 16 | 15.800 | BB | 0.1589 | 12.65432 | 1.12765 | 0.5917 |
| Totals : | | | | 2138.72611 | 558.22250 | |

**Fig. S1. HPLC profile and purity assessment (Area %) of TO1-biotin**

```
=====================================================================
Acq. Operator   : SYSTEM                        Seq. Line :   3
Sample Operator : SYSTEM
Acq. Instrument : LCMS                           Location :   P1-F-07
Injection Date  : 6/7/2022 8:38:20 PM                 Inj :   1
                                               Inj Volume : 2.000 µl
Method          : C:\Users\Public\Documents\ChemStation\1\Data\Seq1 2022-06-07 20-00-05
                  \Purity_test_SEQ.M (Sequence Method)
Last changed    : 6/7/2022 8:00:02 PM by SYSTEM
```

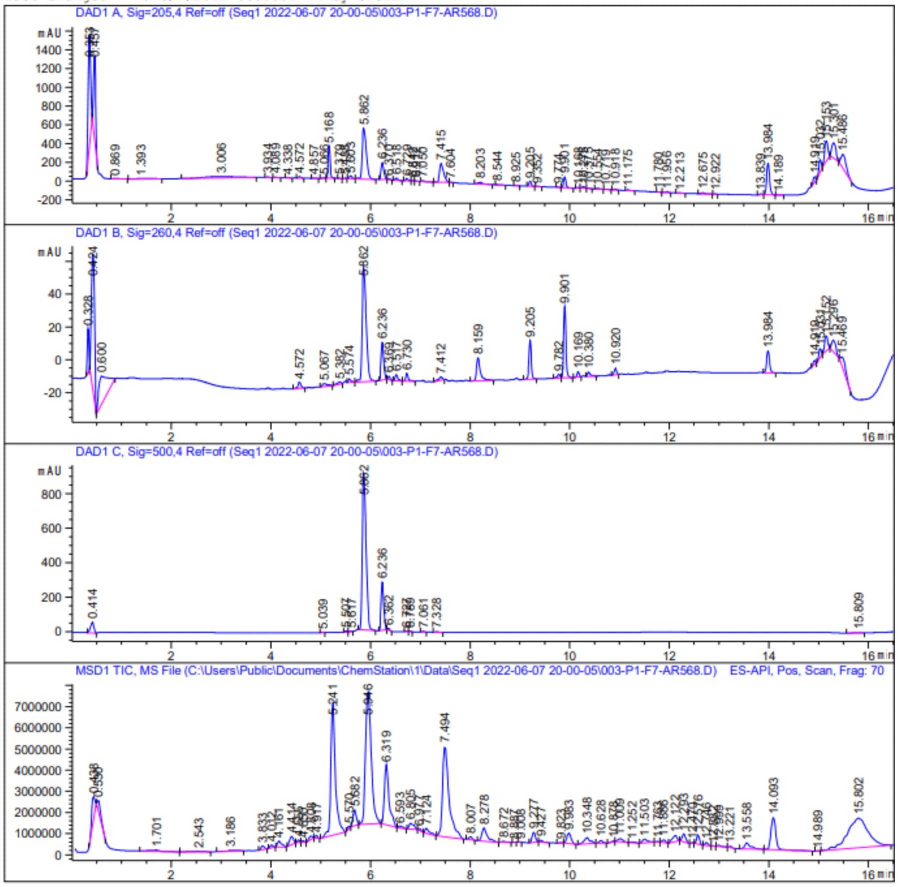

Signal 2: DAD1 B, Sig=260,4 Ref=off

| Peak # | RetTime [min] | Type | Width [min] | Area [mAU*s] | Height [mAU] | Area % |
|---|---|---|---|---|---|---|
| 1 | 0.328 | BB | 0.0365 | 55.86491 | 26.62923 | 3.2155 |
| 2 | 0.424 | BB | 0.0749 | 364.32654 | 83.50948 | 20.9699 |
| 3 | 0.600 | BB | 0.1743 | 204.65083 | 16.77651 | 11.7793 |
| 4 | 4.572 | BB | 0.0634 | 16.22027 | 3.83103 | 0.9336 |
| 5 | 5.067 | BB | 0.1136 | 13.47335 | 1.83614 | 0.7755 |
| 6 | 5.382 | BB | 0.0796 | 7.87303 | 1.44395 | 0.4532 |
| 7 | 5.574 | BB | 0.0556 | 8.25050 | 2.20549 | 0.4749 |
| 8 | 5.862 | BB | 0.0910 | 388.79724 | 69.74603 | 22.3784 |
| 9 | 6.236 | BB | 0.0592 | 84.93285 | 22.93749 | 4.8886 |
| 10 | 6.369 | BB | 0.0477 | 6.47509 | 2.11326 | 0.3727 |
| 11 | 6.517 | BB | 0.0555 | 13.86676 | 3.71574 | 0.7981 |
| 12 | 6.730 | BB | 0.0519 | 17.03082 | 4.98400 | 0.9803 |
| 13 | 7.412 | BB | 0.0822 | 10.53726 | 1.97737 | 0.6065 |
| 14 | 8.159 | BB | 0.0779 | 74.17325 | 13.98298 | 4.2693 |
| 15 | 9.205 | BB | 0.0504 | 77.76118 | 23.59655 | 4.4758 |
| 16 | 9.782 | BB | 0.0589 | 7.95707 | 2.06902 | 0.4580 |
| 17 | 9.901 | BB | 0.0521 | 140.78503 | 43.06445 | 8.1033 |
| 18 | 10.169 | BB | 0.0495 | 10.03316 | 3.29793 | 0.5775 |
| 19 | 10.380 | BB | 0.0654 | 12.05541 | 2.63145 | 0.6939 |
| 20 | 10.920 | BB | 0.0531 | 12.77775 | 3.81622 | 0.7355 |
| 21 | 13.984 | BB | 0.0637 | 54.41923 | 13.32218 | 3.1323 |
| 22 | 14.919 | BB | 0.0583 | 8.53185 | 2.47197 | 0.4911 |
| 23 | 15.031 | BB | 0.0576 | 18.28042 | 5.38979 | 1.0522 |
| 24 | 15.152 | BB | 0.0747 | 41.53691 | 9.19940 | 2.3908 |
| 25 | 15.296 | BB | 0.0996 | 42.22917 | 7.07576 | 2.4306 |
| 26 | 15.469 | BB | 0.1321 | 44.53720 | 5.29875 | 2.5635 |
| Totals : | | | | 1737.37708 | 376.92217 | |

Signal 3: DAD1 C, Sig=500,4 Ref=off

| Peak # | RetTime [min] | Type | Width [min] | Area [mAU*s] | Height [mAU] | Area % |
|---|---|---|---|---|---|---|
| 1 | 0.414 | BB | 0.0738 | 296.74530 | 62.02163 | 4.7487 |
| 2 | 5.039 | BB | 0.0426 | 8.17295 | 3.11775 | 0.1308 |
| 3 | 5.507 | BB | 0.0415 | 16.84101 | 6.66646 | 0.2695 |
| 4 | 5.617 | BB | 0.0407 | 6.44444 | 2.45051 | 0.1031 |
| 5 | 5.862 | BB | 0.0833 | 4781.86279 | 910.46252 | 76.5225 |
| 6 | 6.236 | BB | 0.0600 | 1054.82727 | 279.57550 | 16.8800 |
| 7 | 6.362 | BB | 0.0428 | 28.02592 | 10.60487 | 0.4485 |
| 8 | 6.727 | BB | 0.0390 | 7.42418 | 3.20829 | 0.1188 |
| 9 | 6.789 | BB | 0.0372 | 7.34015 | 3.14812 | 0.1175 |
| 10 | 7.061 | BB | 0.0488 | 14.34298 | 4.81032 | 0.2295 |
| 11 | 7.328 | BB | 0.0589 | 13.21496 | 3.43748 | 0.2115 |
| 12 | 15.809 | BB | 0.1820 | 13.72213 | 1.12708 | 0.2196 |
| Totals : | | | | 6248.96408 | 1290.63055 | |

**Fig. S2. HPLC profile and purity assessment (Area %) of 4a**

**Table S1. Sequences of chemically synthesized oligonucleotides used for evaluating the spectral properties, affinity and selectivity of fluorogenic dyes**.

| code | Sequence, 5'→3' |
|---|---|
| Mango-II | r(GCACGUACGAAGGAGAGGAGAGGAAGAGGAGAGUACGUGC) |
| Mango-IV | r(GCACGUACCGAGGGAGUGGUGAGGAUGAGGCGAGUACGUGC) |
| G4 RNA (utr-z) | r(GGCGGCGGCAGUGGCGGCGG) |
| dsRNA (hairpin ds26) | r(CAAUCGGAUCGAAUUCGAUCCGAUUG) |
| dsDNA (hairpin ds26) | d(CAATCGGATCGAATTCGATCCGATTG) |
| ssDNA ($dT_{18}$) | d(TTTTTTTTTTTTTTTTTT) |
| h/pG4-DNA (hybrid/parallel G4 22AG) | d(AGGGTTAGGGTTAGGGTTAGGG) |
| aG4 (antiparallel G4 22CTA) | d(AGGGCTAGGGCTAGGGCTAGGG) |
| i-motif (hexanucleotide repeat) | d(CCCCGGCCCCGGCCCCGGCCCC) |

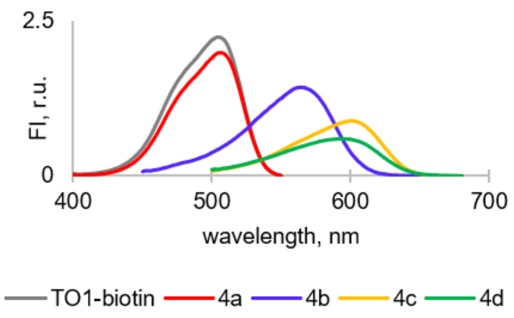

**Fig. S3. Fluorescence excitation spectra of TO1-biotin and new dyes 4a-d in complexes with Mango II.** Conditions: 5 µM dye, 5 µM Mango II, 10 mM Tris-HCl, pH 7.5, and 140 мM KCl. Emission was registered at 540 nm (TO1-biotin and **4a**), 615 nm (**4b**), 640 nm (**4c**), or 650 nm (**4d**).

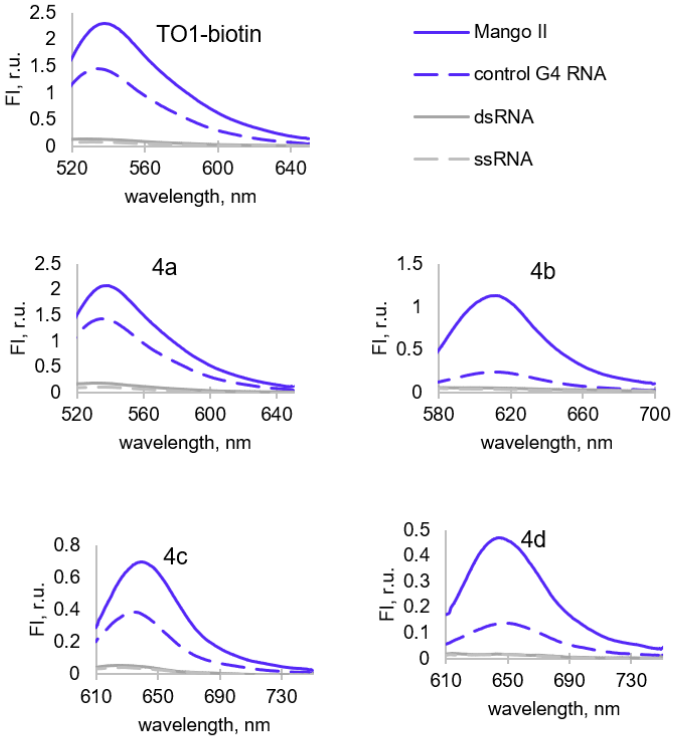

**Fig. S4. Fluorescence emission spectra of TO1-biotin and new ligands 4a-d in complexes with Mango II and control RNA.** Conditions: 5 µM dye, 5 µM Mango II/control G4 RNA, or dsRNA and 25 µg/mL ssRNA, 10 mM Tris-HCl, pH 7.5, and 140 мM KCl. Excitation at 505 nm (TO1-biotin and **4a**), 560 nm (**4b**), and 595 nm (**4c** and **4d**).

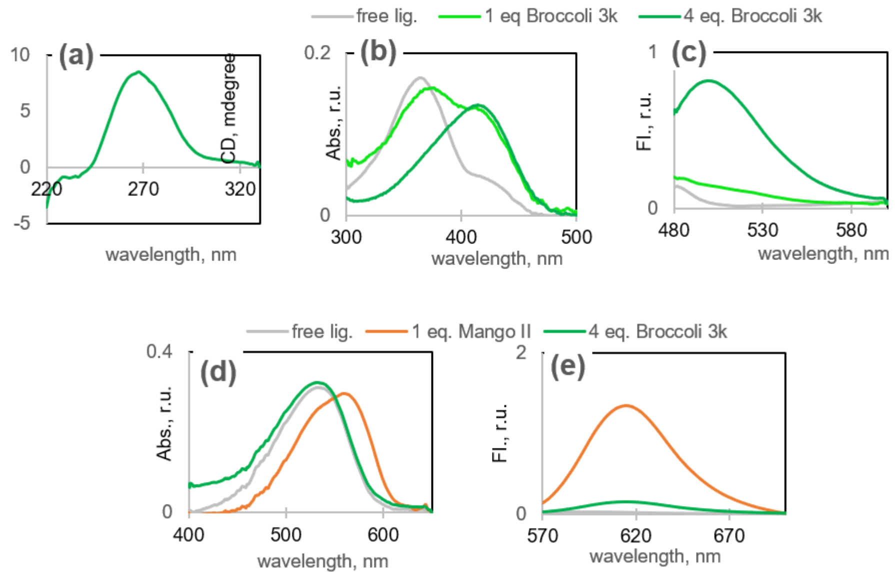

**Fig. S5. Orthogonal RNA-dye pairs Broccoli-DFHBI and Mango II-4b.** (a) Verification of Broccoli RNA secondary structure using circular dichroism spectroscopy (CD). Conditions: 5 µM Broccoli, 10 mM Tris-HCl, pH 7.5, and 140 mM KCl. (b) Absorbance of **DFHBI** in free state and in complex with Broccoli. (c) Fluorescence of DFHBI in free state and in complex with Broccoli (excitation: 470 nm). (d) Absorbance of **4b** with or without RNA aptamers. (e) Fluorescence of **4b** with or without RNA aptamers (excitation: 470 nm). Conditions: 5 µM dye, 10 mM Tris-HCl, pH 7.5, and 140 mM KCl.

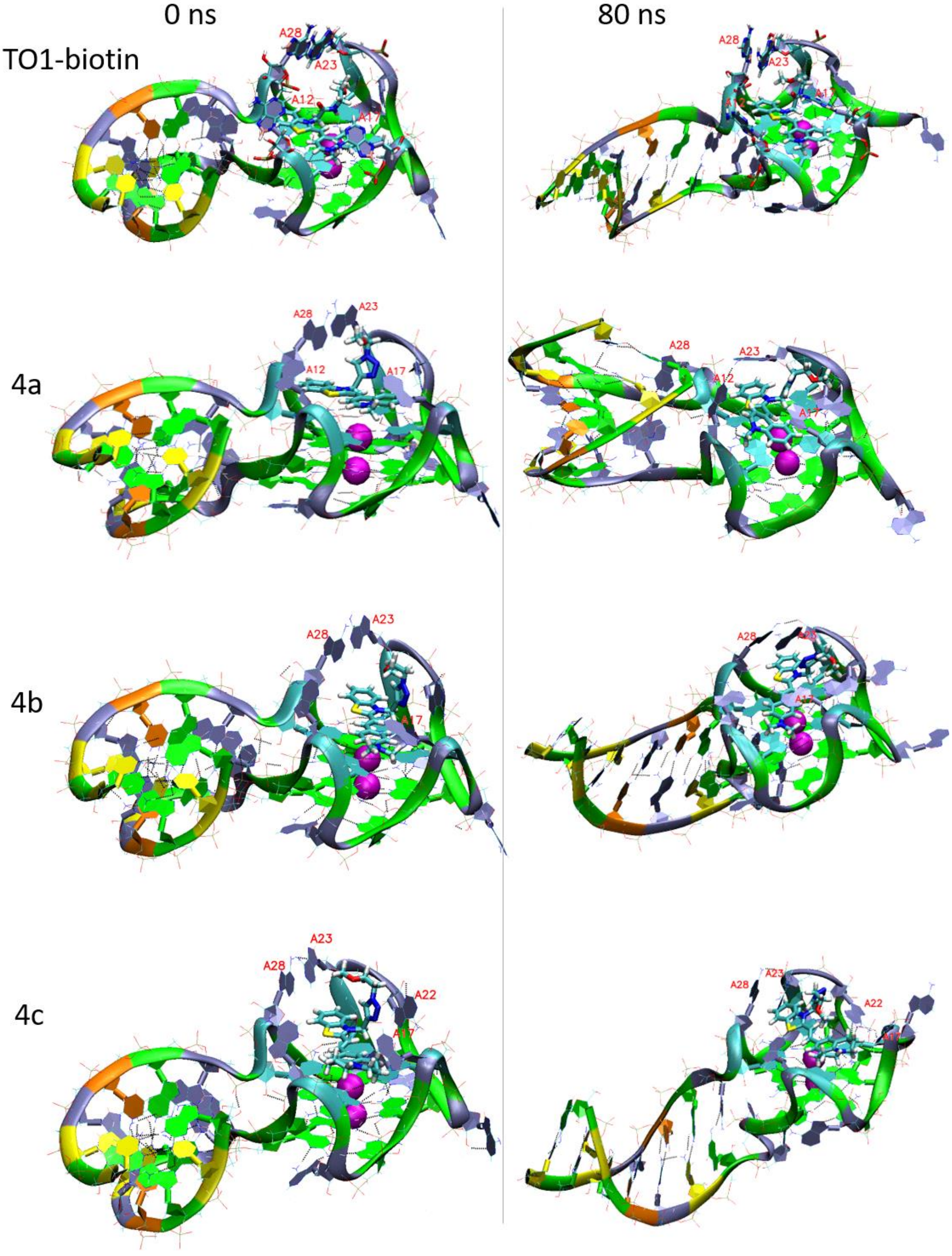

**Fig. S6. Molecular dynamics of dye-Mango II complexes: full structures of the complexes at 0 and 80 ns simulation times**. Color labeling of aptamer residues: G, green; A, grey; T, orange; C, cyan. Ligand atoms: C, cyan; N, blue; O, red; S, yellow.

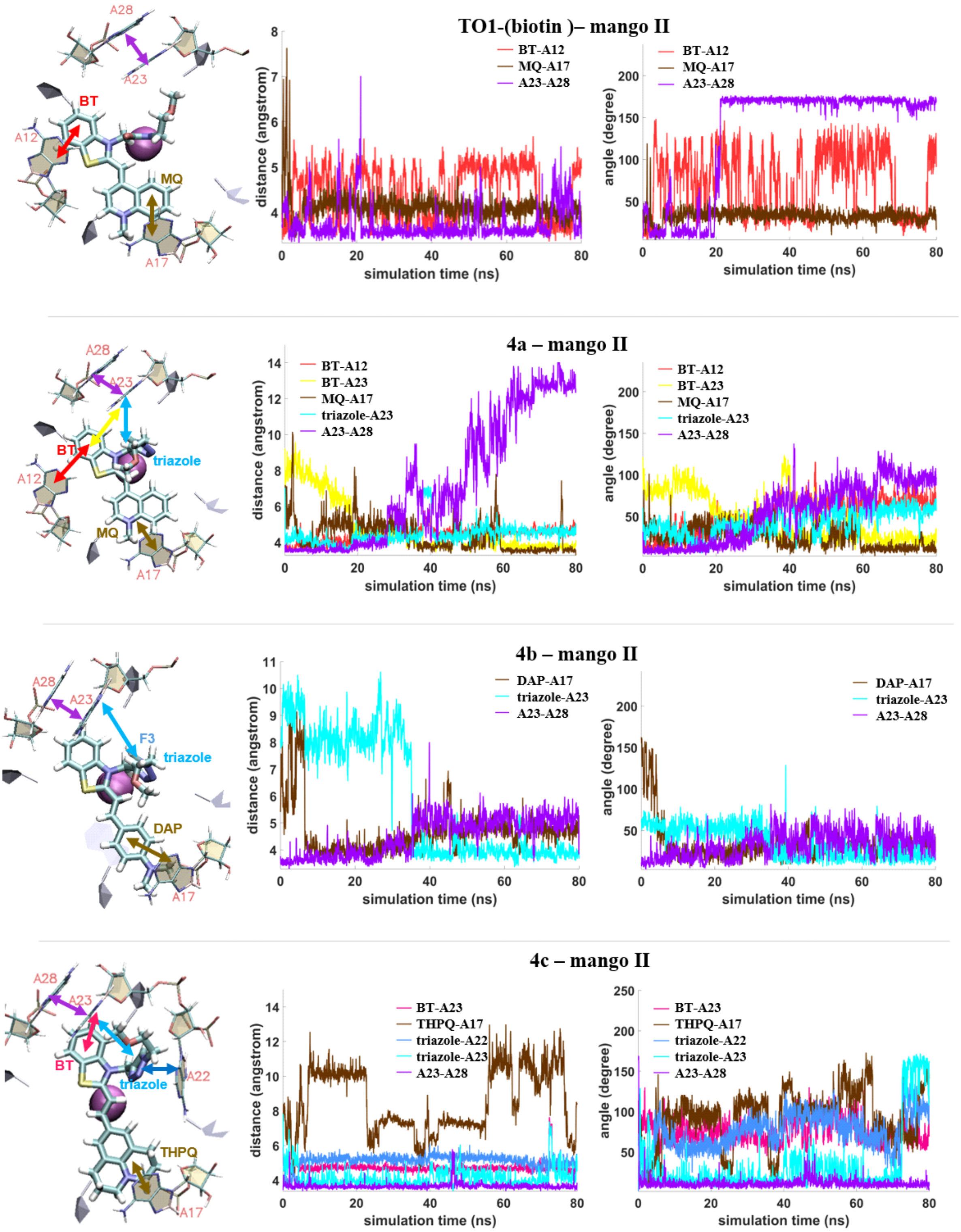

**Fig. S7. Molecular dynamics of dye-Mango II complexes: binding modes**. Left: dye positioning in complex with mango II. Arrows mark distances between COMs of key dye and aptamer residues. Middle: distances between the centers of mass (COM) of the specified residues. Right: angles between the normal to the planes of these residues. BT, benzothiazolium ring; MQ, N-methylquinolinyl group; DAP, *N*,*N*-dimethylaminophenyl group; THPQ, 2,3,6,7-tetrahydro-H,5H-pyrido[3,2,1-ij]quinolinyl group. In each pair of the residues, only the closest rings were considered, and for these rings COM distances and angles were analyzed.

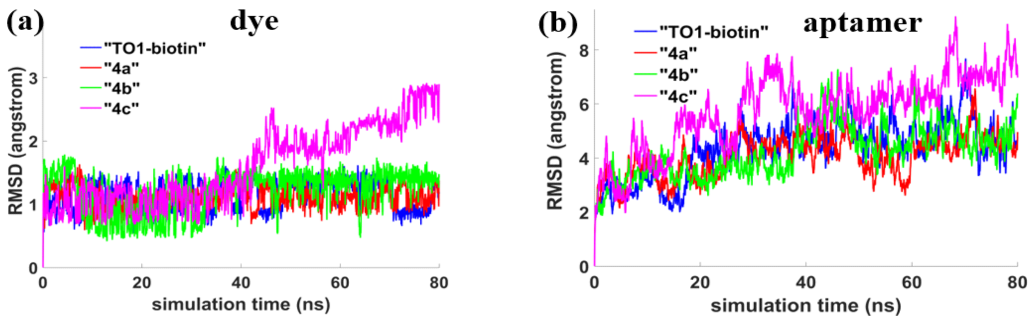

**Fig. S8. Molecular dynamics of dye-Mango II complexes: RMSD of the dyes (a) and the aptamer (b).**

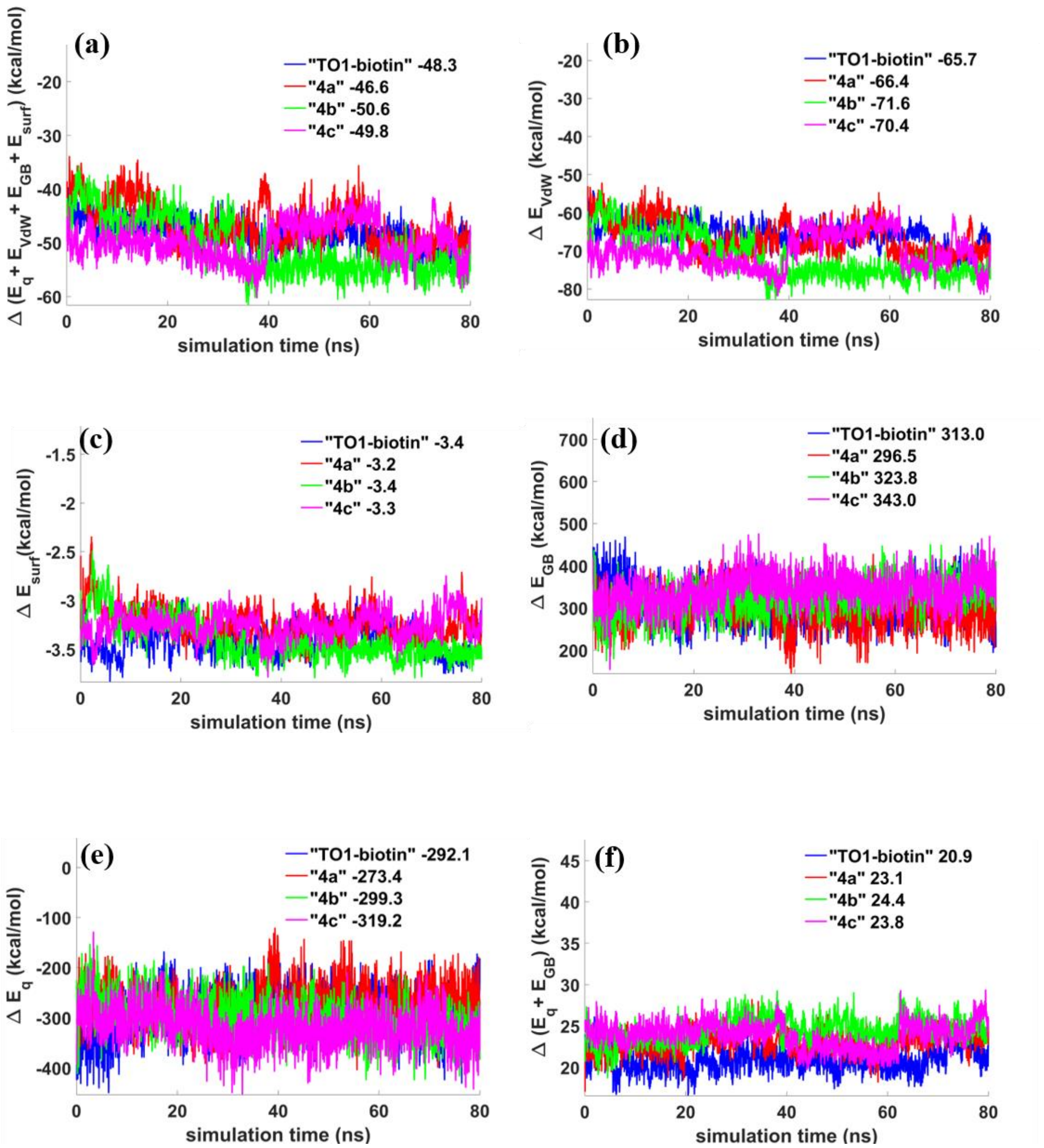

**Fig. S9. Molecular dynamics simulation: free energy plots**. (a) Total RNA-dye interaction energy. (b) Contribution of Van der Waals interactions. (c) Non-polar contribution to solvation energy. (d) Polar contribution to solvation energy. (e) Contribution of electrostatic interactions. (f) Electrostatic and polar solvation energy.

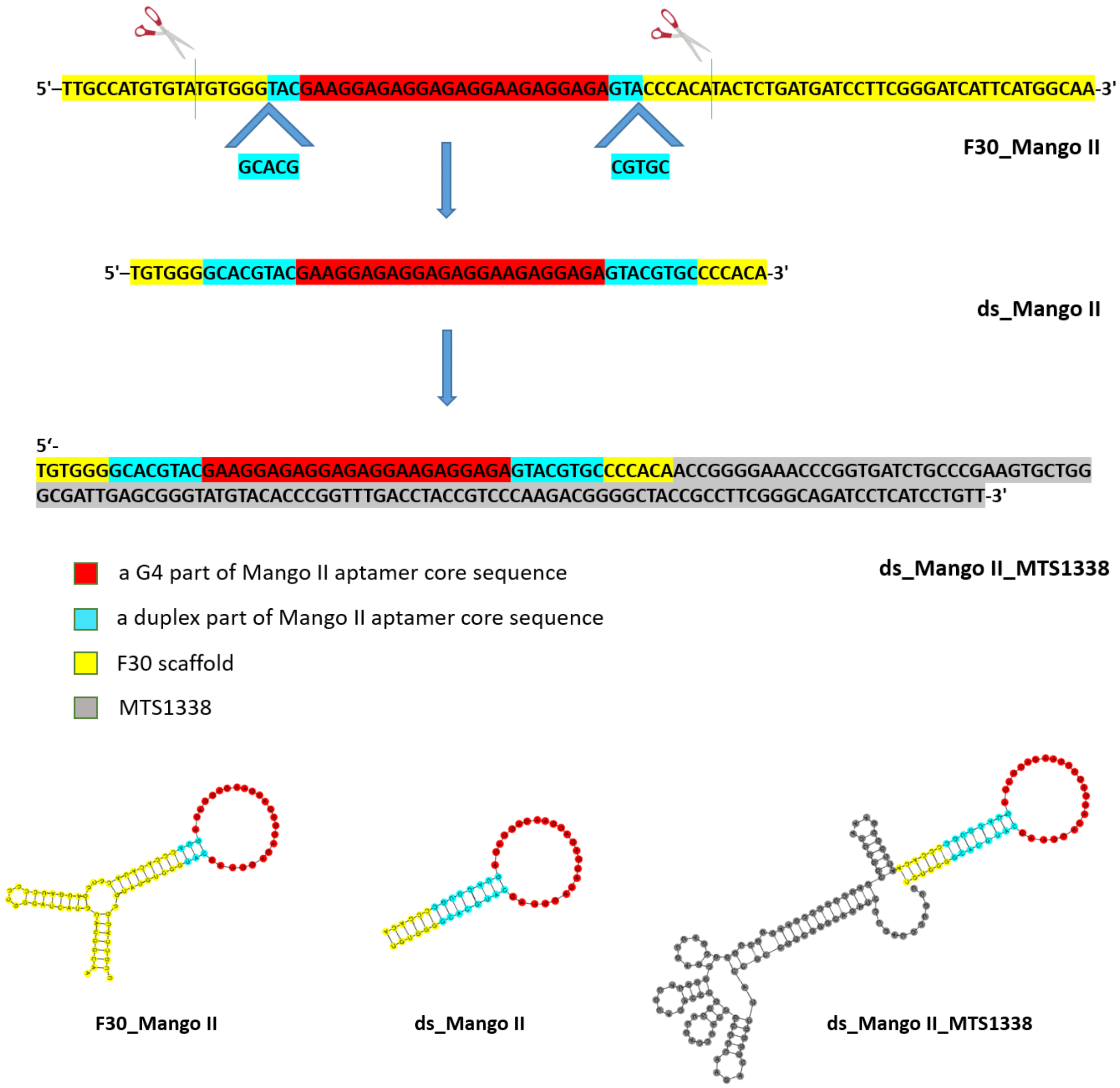

**Fig. S10. The sequences and the secondary structure of F30_Mango II, ds_Mango II aptamers, and modular RNA ds_Mango II_MTS1338 according to the RNAfold server** (http://rna.-tbi.univie.ac.at/cgi-bin/RNAWebSuite/RNAfold.cgi).

**Table S2. Nucleotide sequences of the genetic constructs and the oligonucleotides used for their analysis**

| code | sequence 5'→3' |
|---|---|
| rrnB promoter | CTAGAGTGACCGCGTCTGACCAGGGAAAATAGCCCTCTGACCTGGGGATTTGACTCCC AGTTTCCAAGGACGTAACTTA |
| MTS1338 | ACCGGGGAAACCCGGTGATCTGCCCGAAGTGCTGGGCGATTGAGCGGGTATGTACAC CCGGTTTGACCTACCGTCCCAAGACGGGGCTACCGCCTTCGGGCAGATCCTCATCCTGTT |
| term terminator | GCTTCCCCGCGAAAGCGGGGTTTTTTTTTTTAGCT |
| ds_Mango II | TGTGGGGCACGTACGAAGGAGAGGAGAGGAAGAGGAGAGTACGTGCCCCACA |
| Broccoli | GAGACGGTCGGGTCCAGATATTCGTATCTGTCGAGTAGAGTGTGGGCTC |
| rrnB_ds_Mango II_term | CTAGAGTGACCGCGTCTGACCAGGGAAAATAGCCCTCTGACCTGGGGATTTGACTCCCAGTTTCCAAGGACGTAACTTATGTGGGGCACGTACGAAGGAGAGGAGAGGAAGAGGAGAGTACGTGCCCCACACCCCGCGAAAGCGGGGTTTTTTTTTTTAGCT- |
| rrnB_ds_Mango II_MTS1338_term | CTAGAGTGACCGCGTCTGACCAGGGAAAATAGCCCTCTGACCTGGGGATTTGACTCCCAGTTTCCAAGGACGTAACTTATGTGGGGCACGTACGAAGGAGAGGAGAGGAAGAGGAGAGTACGTGCCCCACAACCGGGGAAACCCGGTGATCTGCCCGAAGTGCTGGGCGATTGAGCGGGTATGTACACCCGGTTTGACCTACCGTCCCAAGACGGGGCTACCGCCTTCGGGCAGATCCTCATCCTGTTCCCCGCGAAAGCGGGGTTTTTTTTTTTAGCT |
| rrnB_Broccoli _term | CTAGAGTGACCGCGTCTGACCAGGGAAAATAGCCCTCTGACCTGGGGATTTGACTCCCAGTTTCCAAGGACGTAACTTAGAGACGGTCGGGTCCAGATATTCGTATCTGTCGAGTAGAGTGTGGGCTCCCCCGCGAAAGCGGGGTTTTTTTTTTTAGCT |
| T7MngII_F | GTTTTTTTTAATACGACTCACTATAGGTGTGGGGCACGTAC |
| MngII_R | TGTGGGGCACGTACTCTCCT |
| 1338_R | AACAGGATGAGGATCTGCCCG |
| T7Broc_F | GTTTTTTTTAATACGACTCACTATAGGGAGACGGTCGGGTC |
| Broc_R | GAGCCCACACTCTACTCGACAGATACGAAT |
| q1338-F | GTGCTGGGCGATTGAGC |
| q1338-R | GCGGTAGCCCCGTCTT |
| 16S-F | TACGTAGGGTGCGAGCGTTG |
| 16S-R | CCCGCACGCTCACAGTTAAG |

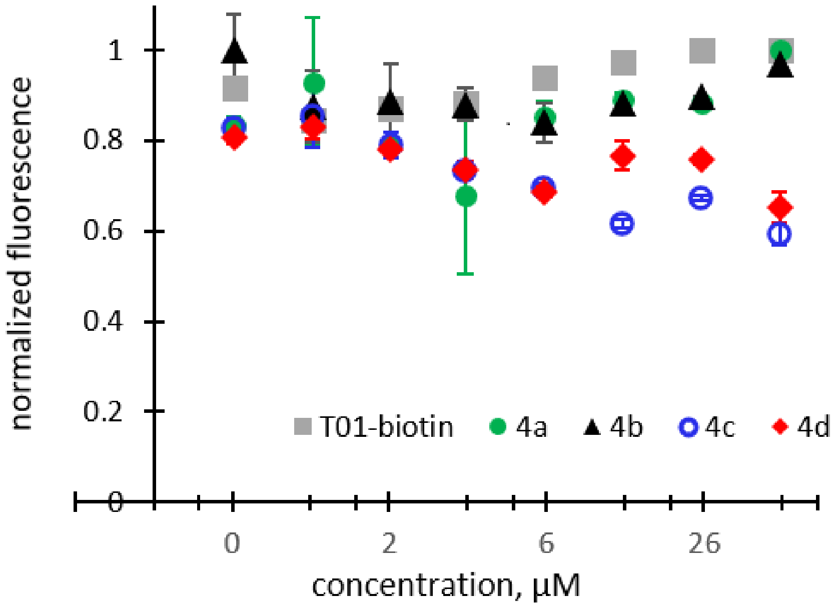

**Fig. S11. Cytotoxicity of TO1-biotin and 4a-d in RAW 264.7 cell lines after 7 d incubation.**

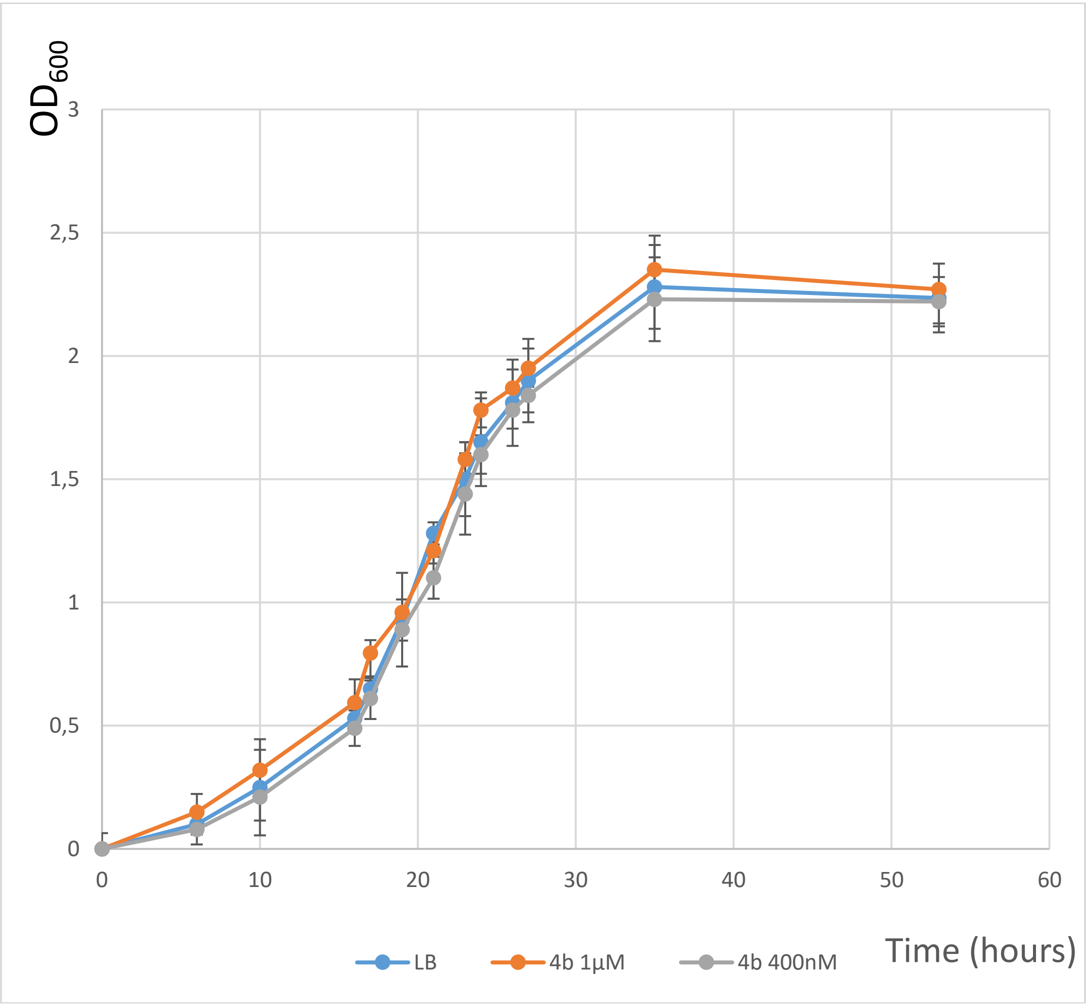

**Fig. S12. Evaluation of the dye cytotoxicity for bacterial cells using optical density ($OD_{600}$) measurements**. Growth curves of *M. smegmatis*_pAMYC in the presence of **4b** in concentrations 400 nM, 1 µM and without dye (positive control). The data are presented as the mean +-SD of three independent experiments.

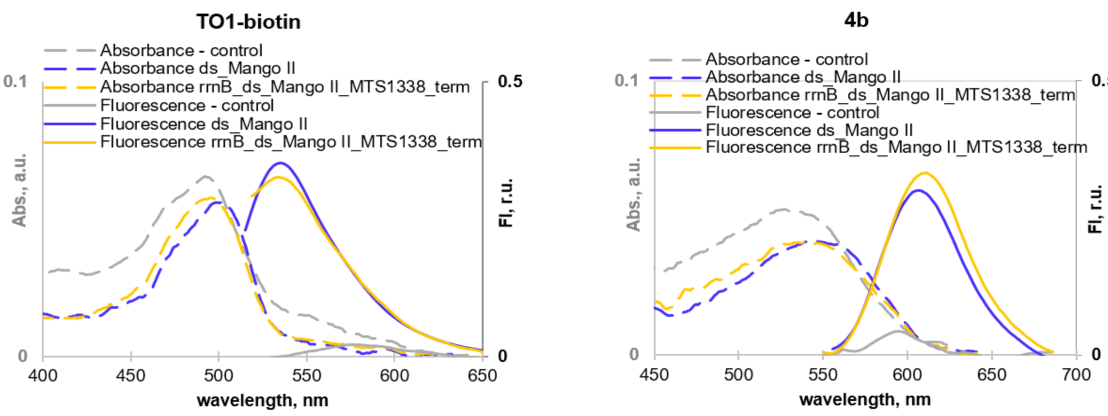

**Fig. S13. Spectra of TO1-biotin and 4b in complexes with the Mango II-labeled genetic construct and control RNA.** Absorption and fluorescence emission spectra of the dyes in complexes with rrnB_Mango II_MTS1338_term, ds_Mango II, and random-sequence Mango-free (-control) RNA. Conditions in A and B: 1 µM RNA, 1 µM dye, 20 mM sodium phosphate buffer, pH 7.2, 0.05% Tween-20, and 140 mM KCl.

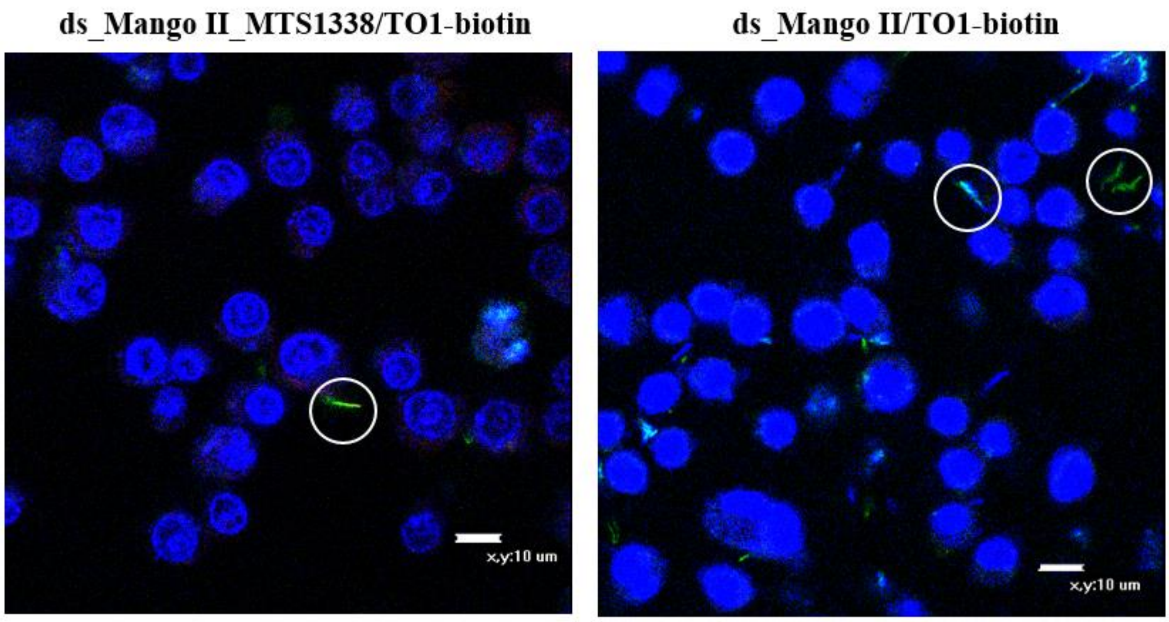

**Fig S14. Visualization of the genetically encoded modular RNA ds_Mango II_MTS1338 and ds_Mango II tag in M. smegmatis in infected RAW 264.7 macrophages using TO1-biotin (in a green channel)**. Macrophage nuclei were stained with Hoechst 33258 (in a blue channel). The circles cover fluorescent bacteria.

### NMR spectra

*2-((1-methylquinolin-4(1H)-ylidene)methyl)-3-(prop-2-yn-1-yl)benzo[d]thiazol-3-ium bromide* **2a**

$^{1}H$ spectrum

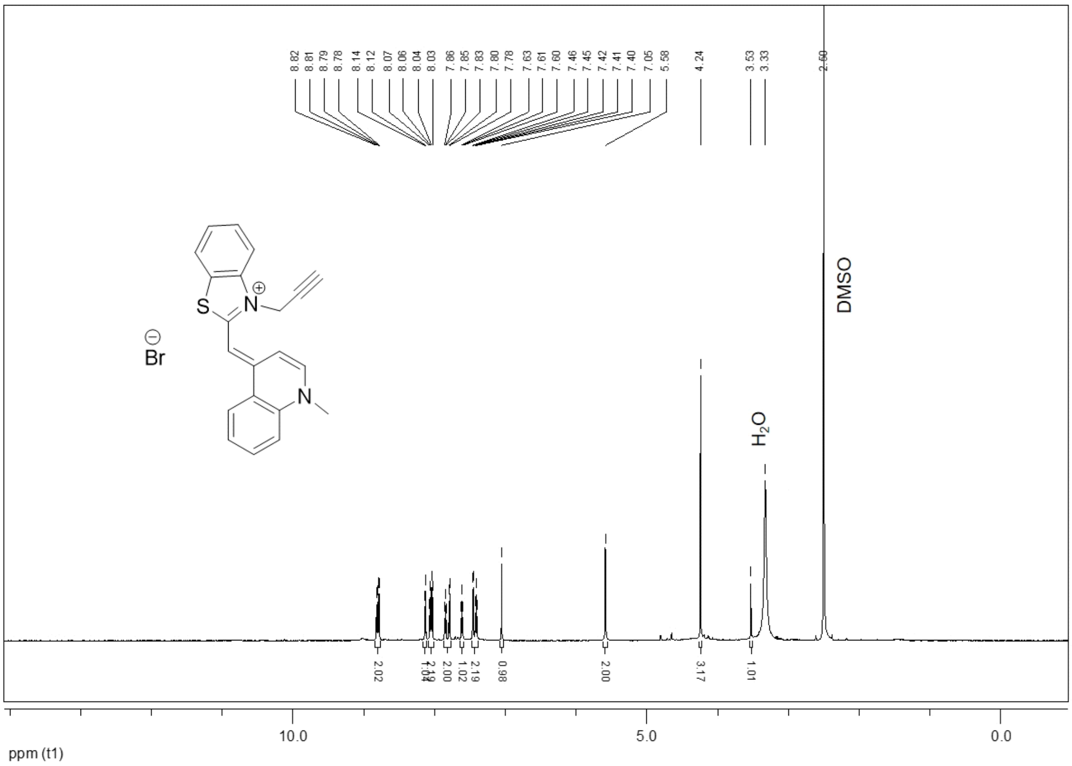

$^{13}C$ spectrum

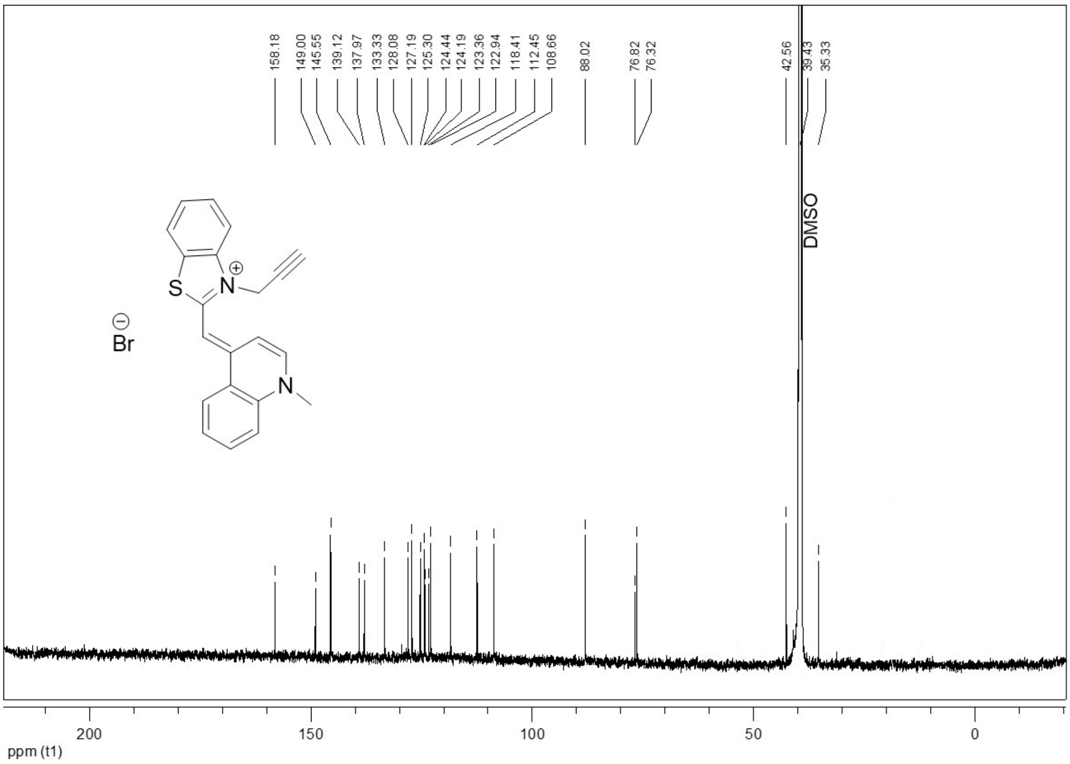

*2-(2-(4-(Dimethylamino)phenyl)ethenyl)-3-(2-propynyl)benzo[d]thiazolium bromide* **2b**

$^1$H spectrum

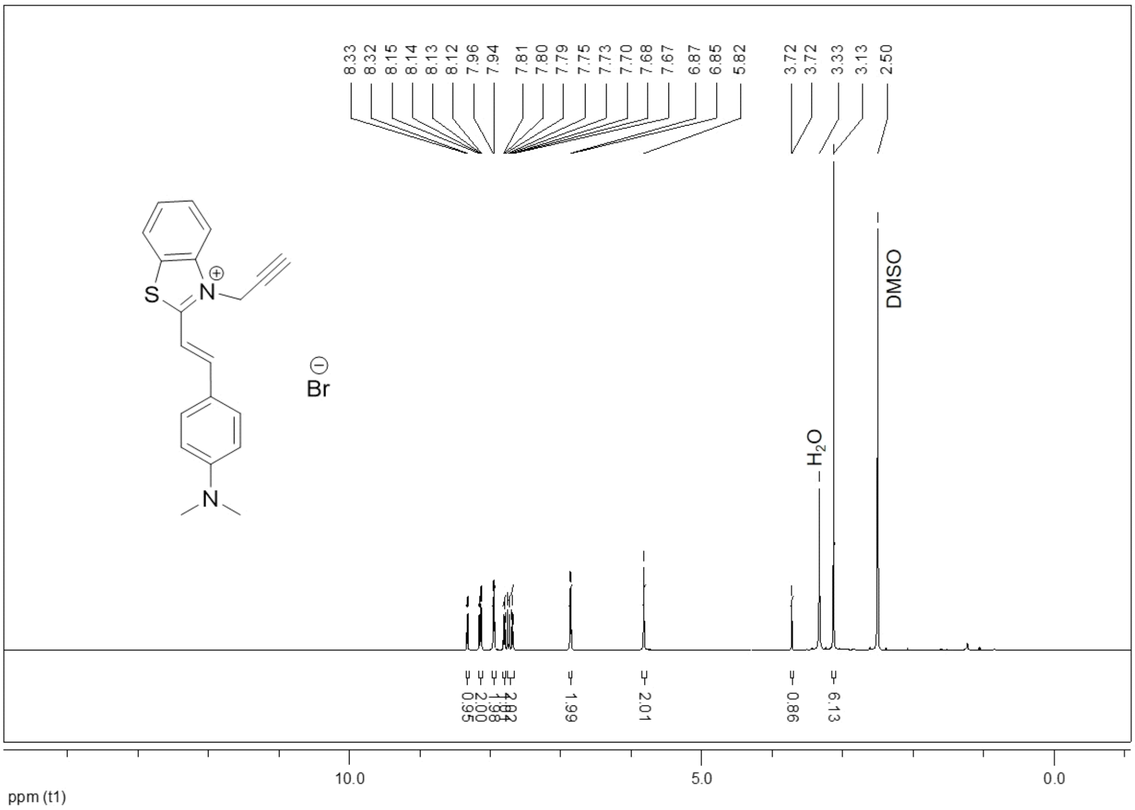

$^{13}$C spectrum

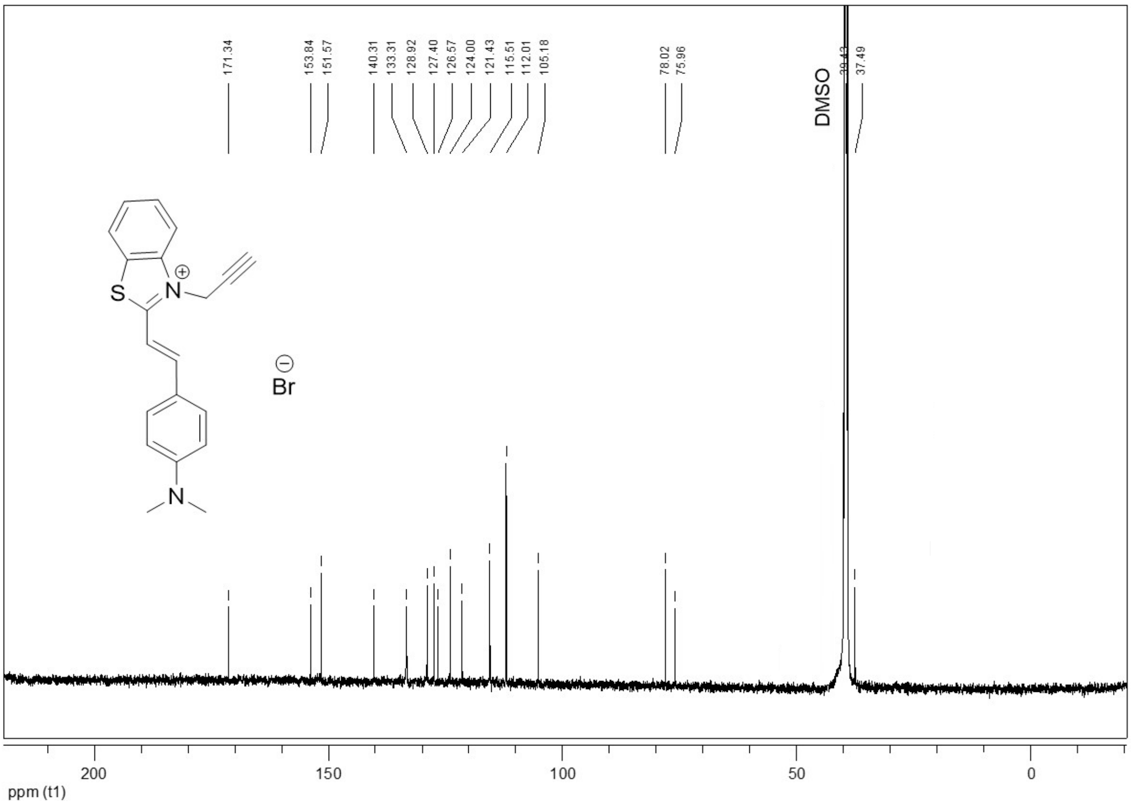

*3-(prop-2-yn-1-yl)-2-(2-(2,3,6,7-tetrahydro-1H,5H-pyrido[3,2,1-ij]quinolin-9-yl)vinyl)benzo[d]thiazol-3-ium bromide* **2c**

$^{1}$H spectrum

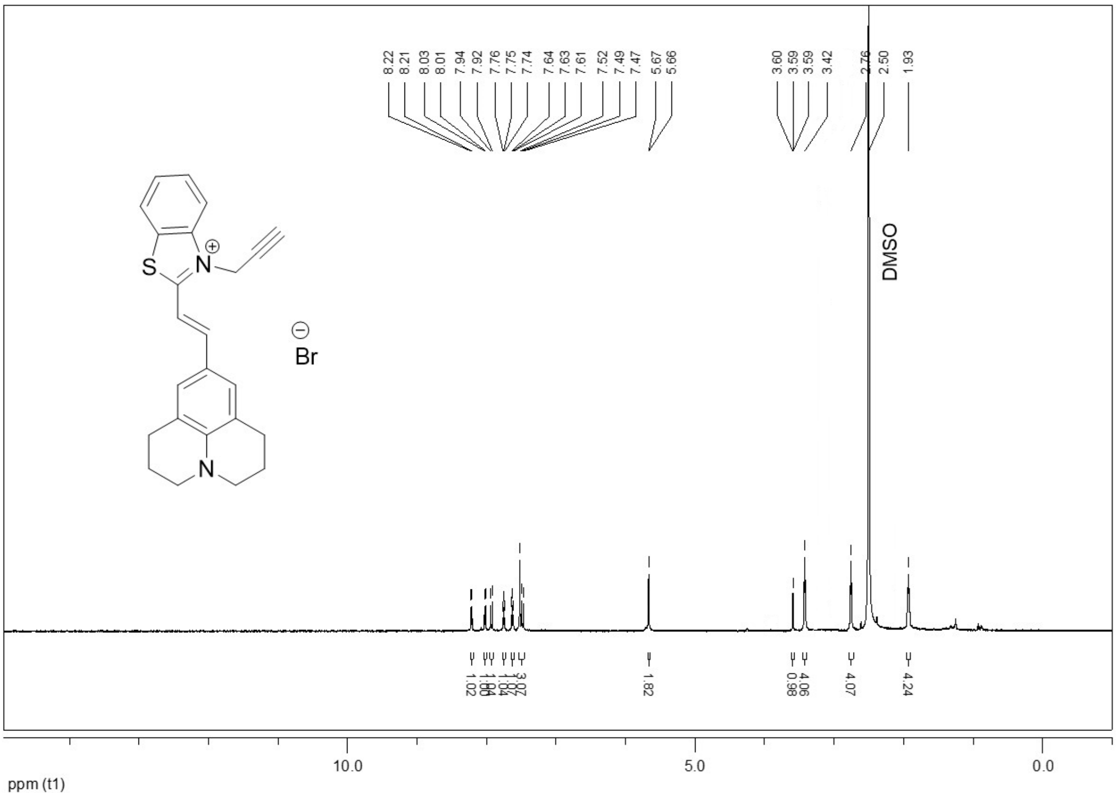

*3-(prop-2-yn-1-yl)-2-(2-(1,2,2,4-tetramethyl-1,2-dihydroquinolin-6-yl)vinyl)benzo[d]thiazol-3-ium bromide* **2d**

$^{1}$H spectrum

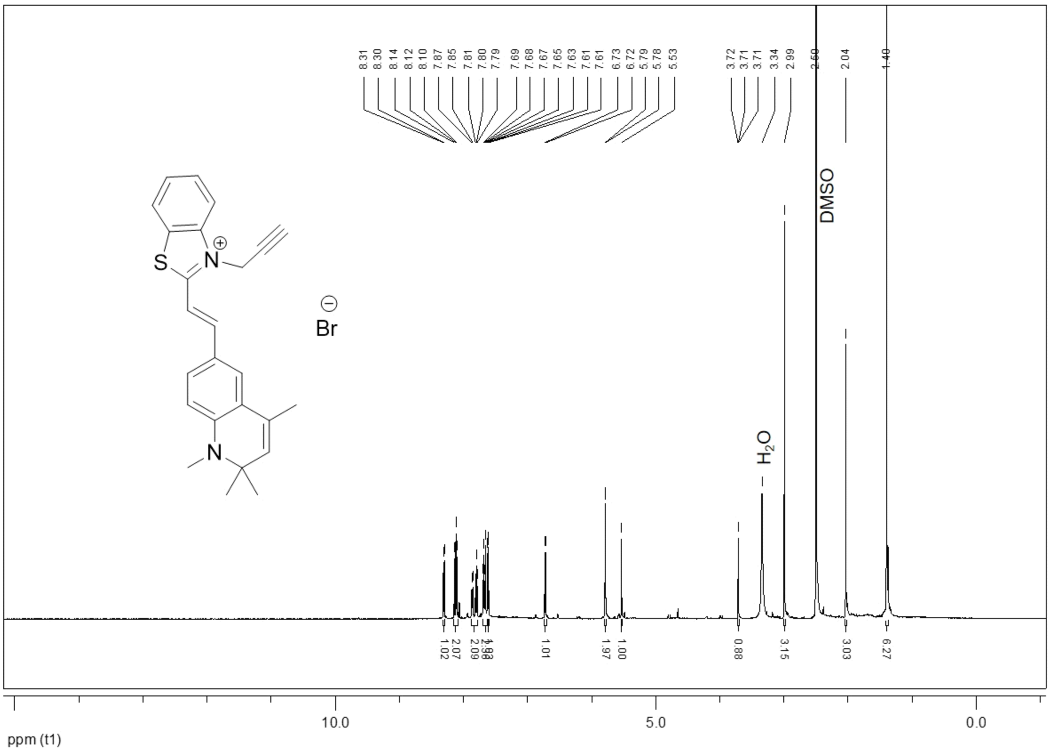

$^{13}$C spectrum

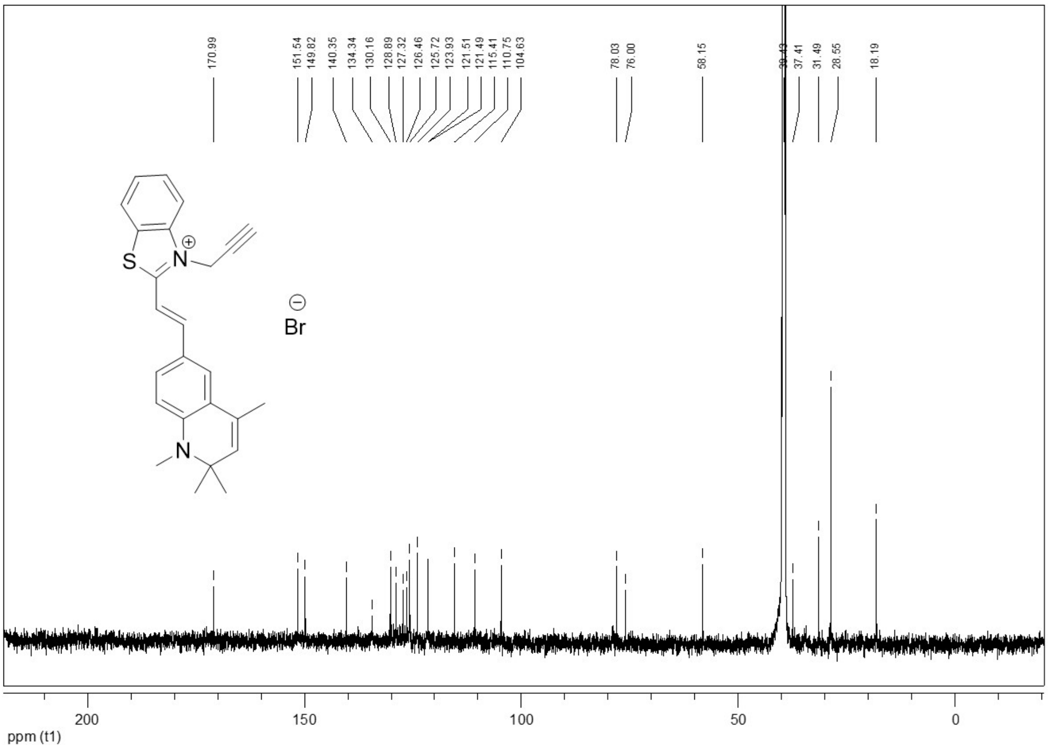

*N-(2-(2-(2-(2-azidoethoxy)ethoxy)ethoxy)ethyl)-5-(2-oxohexahydro-1H-thieno[3,4-d]imidazol-4-yl)pentanamide* **3**

$^{1}$H spectrum

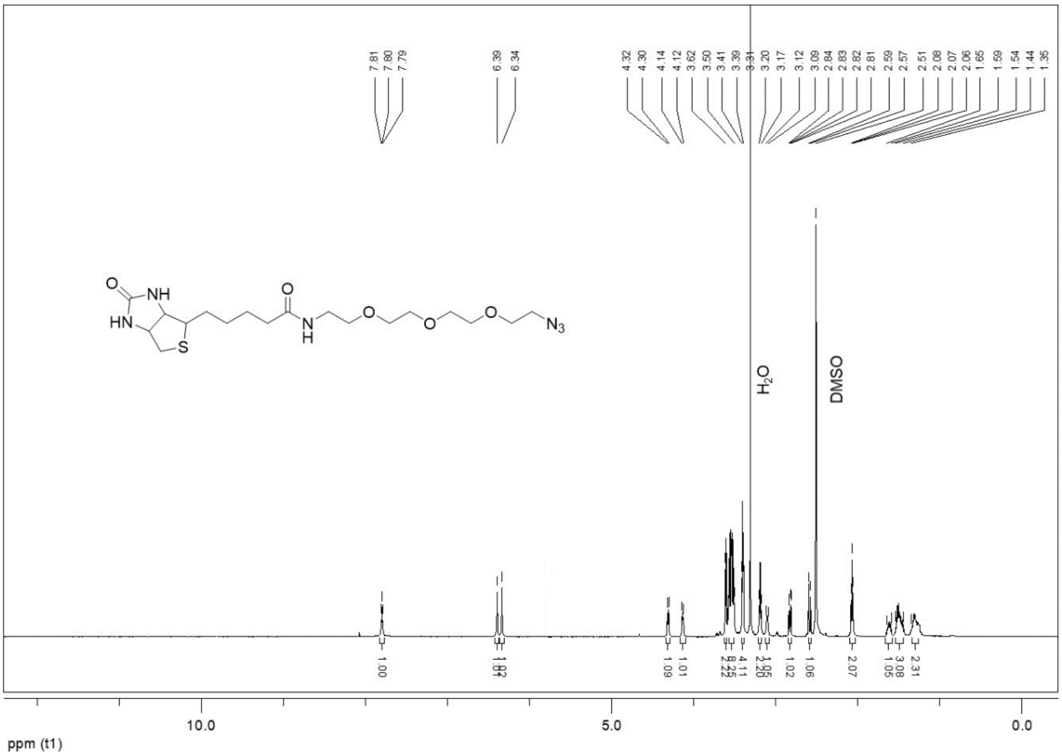

$^{13}$C spectrum

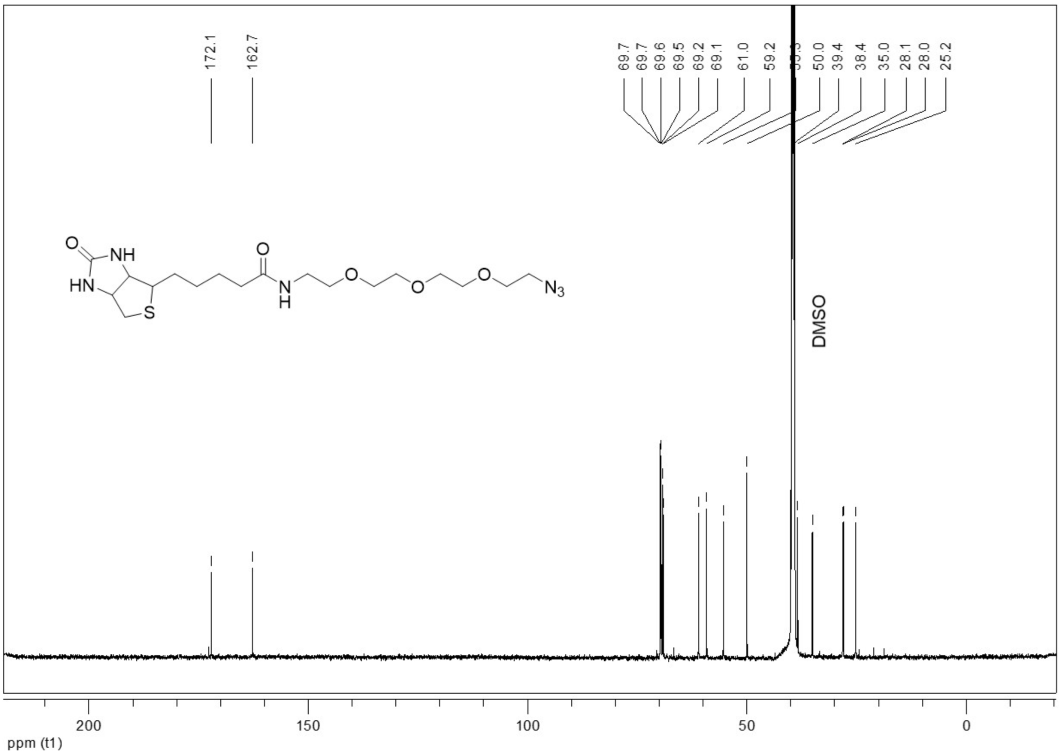

*2-((1-methylquinolin-4(1H)-ylidene)methyl)-3-((1-(13-oxo-17-(2-oxohexahydro-1H-thieno[3,4-d]imidazol-4-yl)-3,6,9-trioxa-12-azaheptadecyl)-1H-1,2,3-triazol-4-yl)methyl)benzo[d]thiazol-3-ium 2,2,2-trifluoroacetate* **4a**

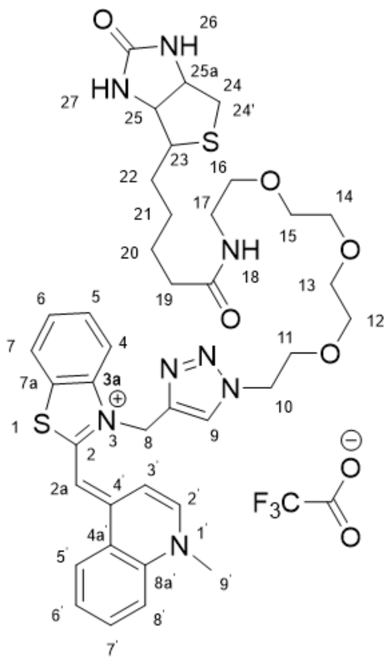

$^{1}$H spectrum

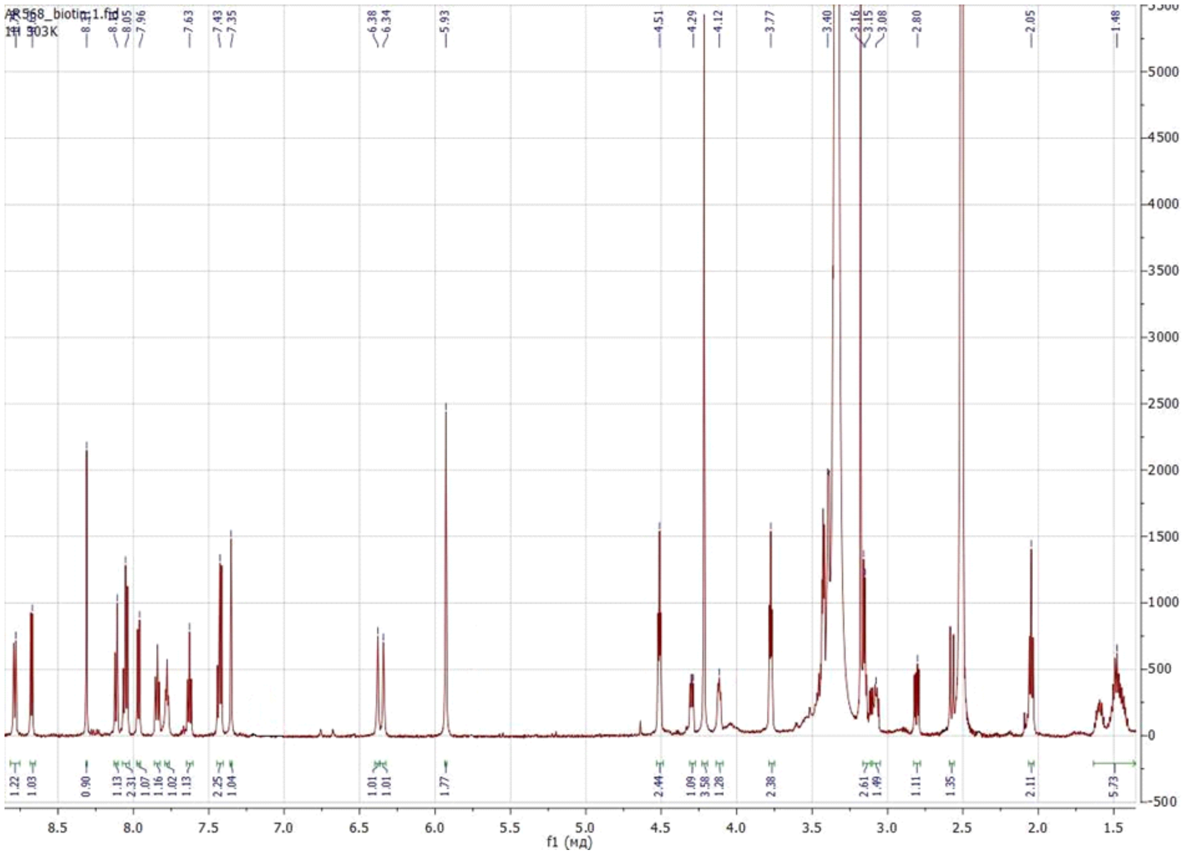

An expansion of the 2D $^{1}$H COSY NMR spectrum (aromatic region)

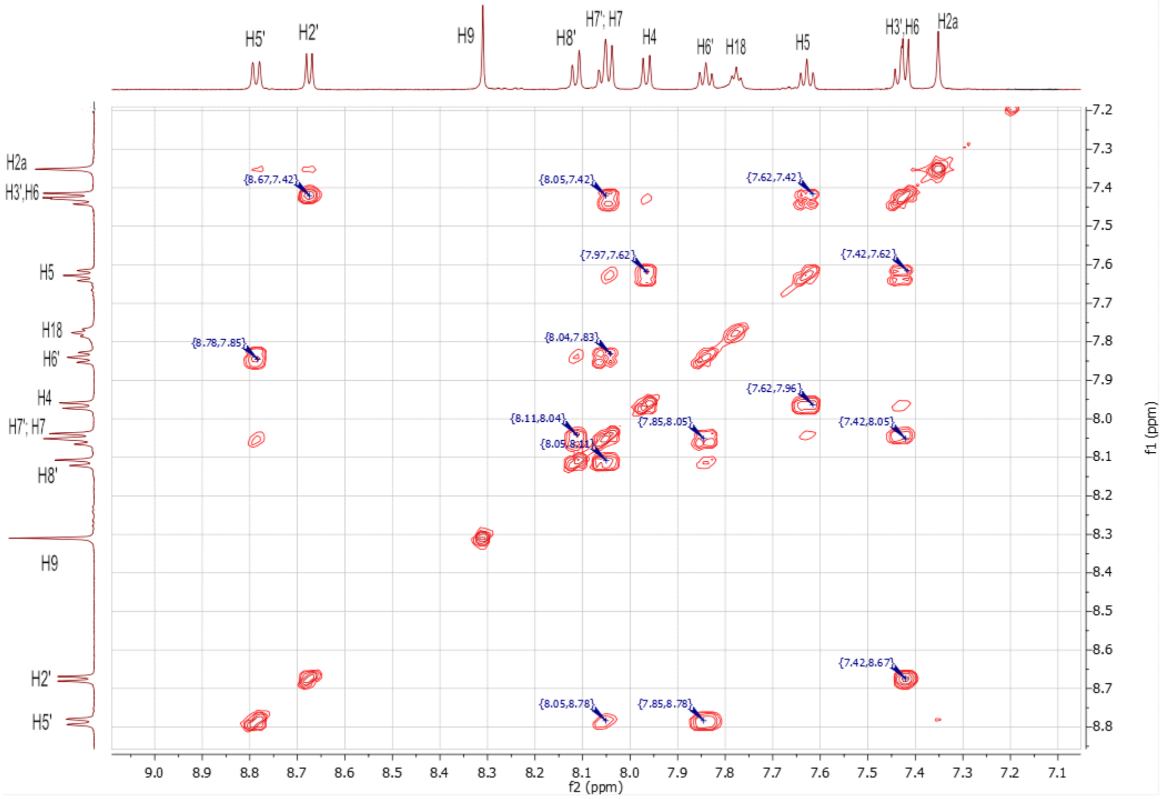

An expansion of the 2D $^{1}$H COSY NMR spectrum (aliphatic region)

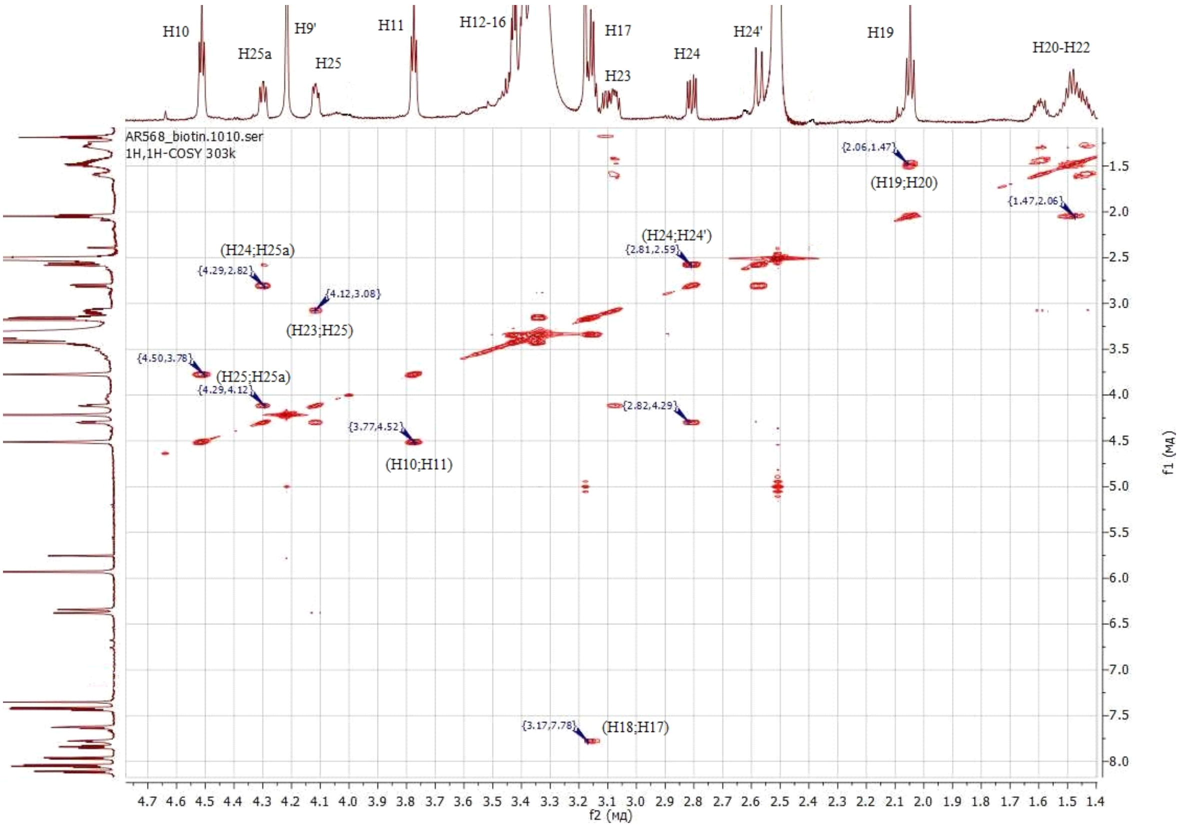

*2-(4-(dimethylamino)styryl)-3-((1-(13-oxo-17-(2-oxohexahydro-1H-thieno[3,4-d]imidazol-4-yl)-3,6,9-trioxa-12-azaheptadecyl)-1H-1,2,3-triazol-4-yl)methyl)benzo[d]thiazol-3-ium 2,2,2-trifluoroacetate* **4b**

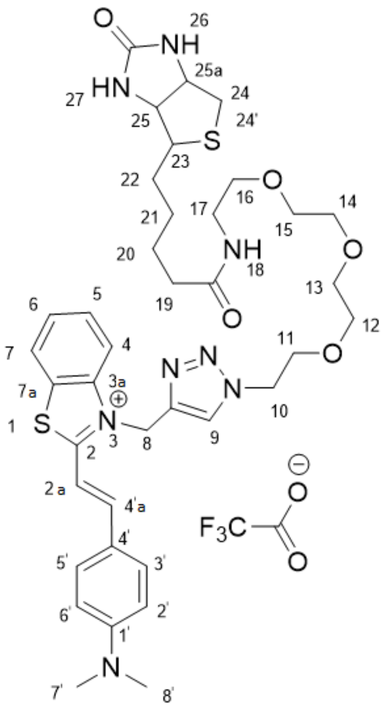

$^{1}$H spectrum

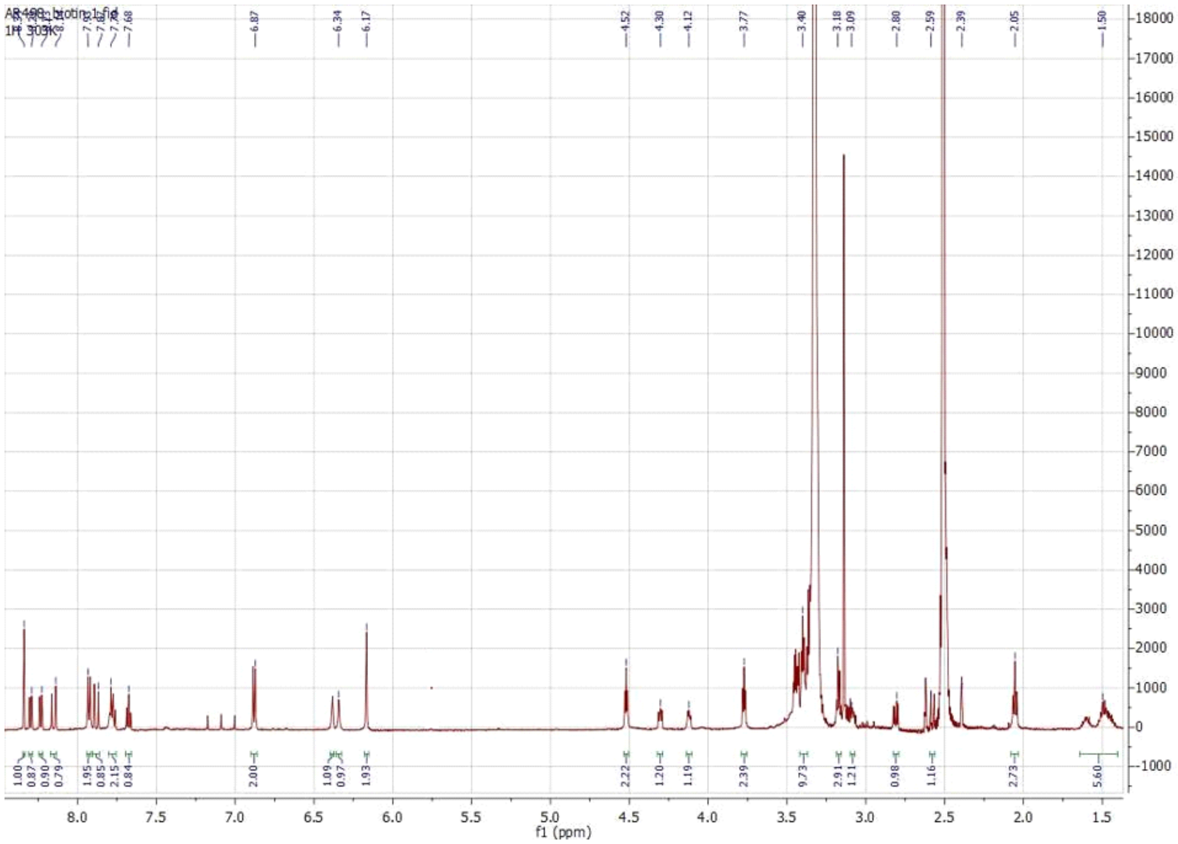

An expansion of the 2D $^{1}H$ COSY NMR spectrum (aromatic region)

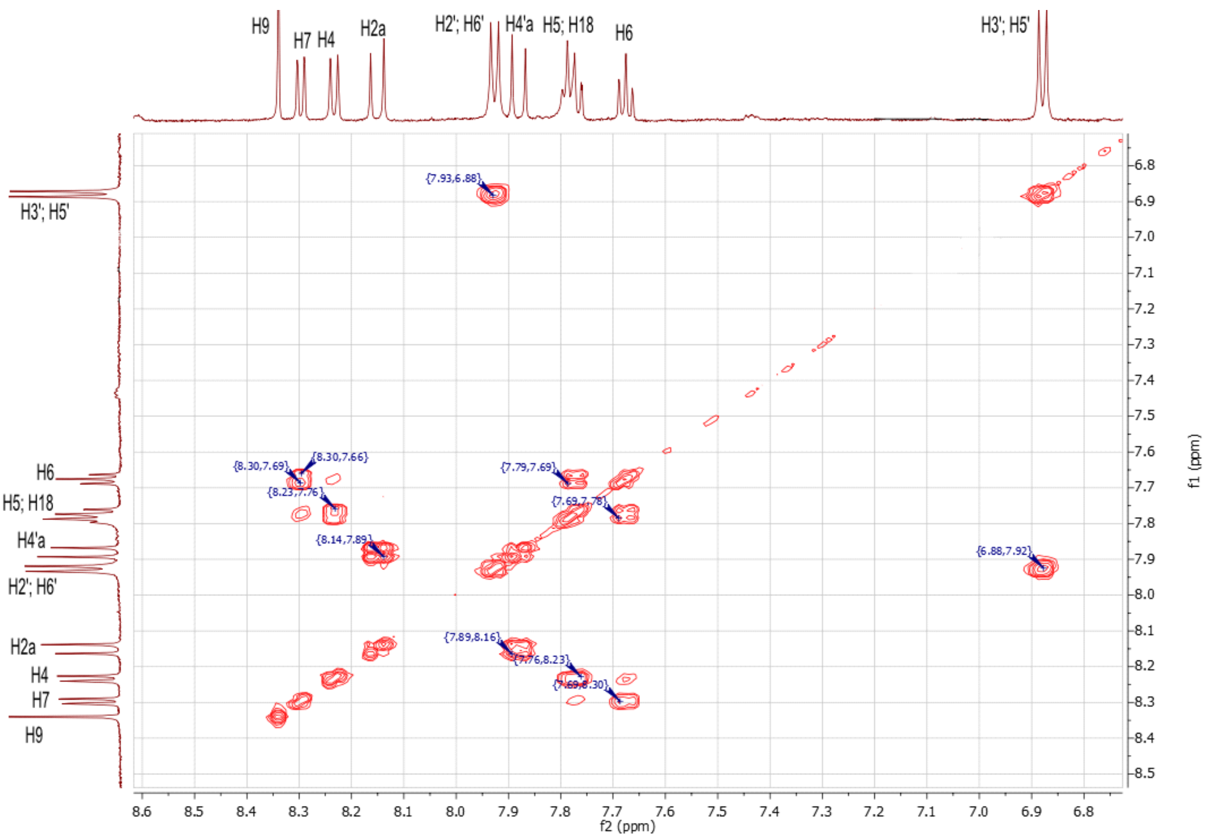

An expansion of the 2D $^{1}H$ COSY NMR spectrum (aliphatic region)

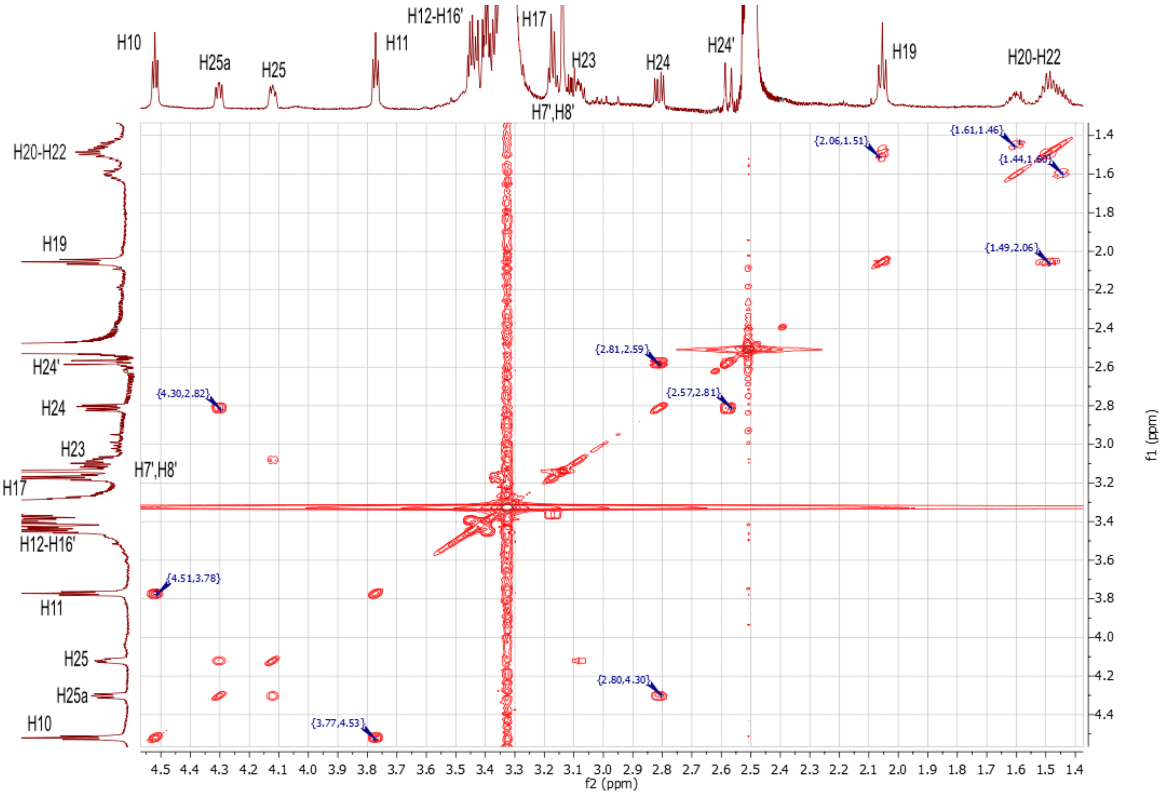

*3-((1-(13-oxo-17-(2-oxohexahydro-1H-thieno[3,4-d]imidazol-4-yl)-3,6,9-trioxa-12-azaheptadecyl)-1H-1,2,3-triazol-4-yl)methyl)-2-(2-(2,3,6,7-tetrahydro-1H,5H-pyrido[3,2,1-ij]quinolin-9-yl)vinyl)benzo[d]thiazol-3-ium 2,2,2-trifluoroacetate* **4c**

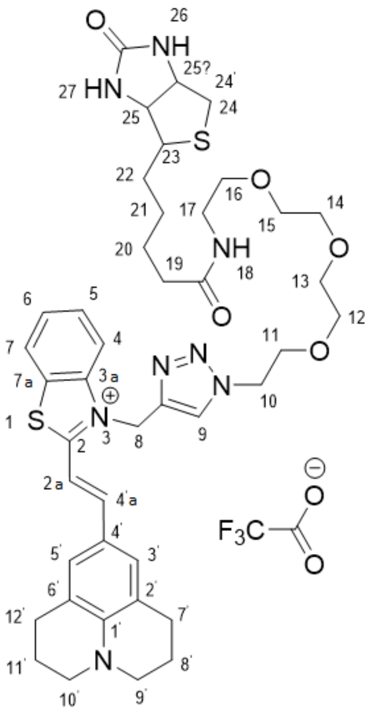

$^1$H spectrum

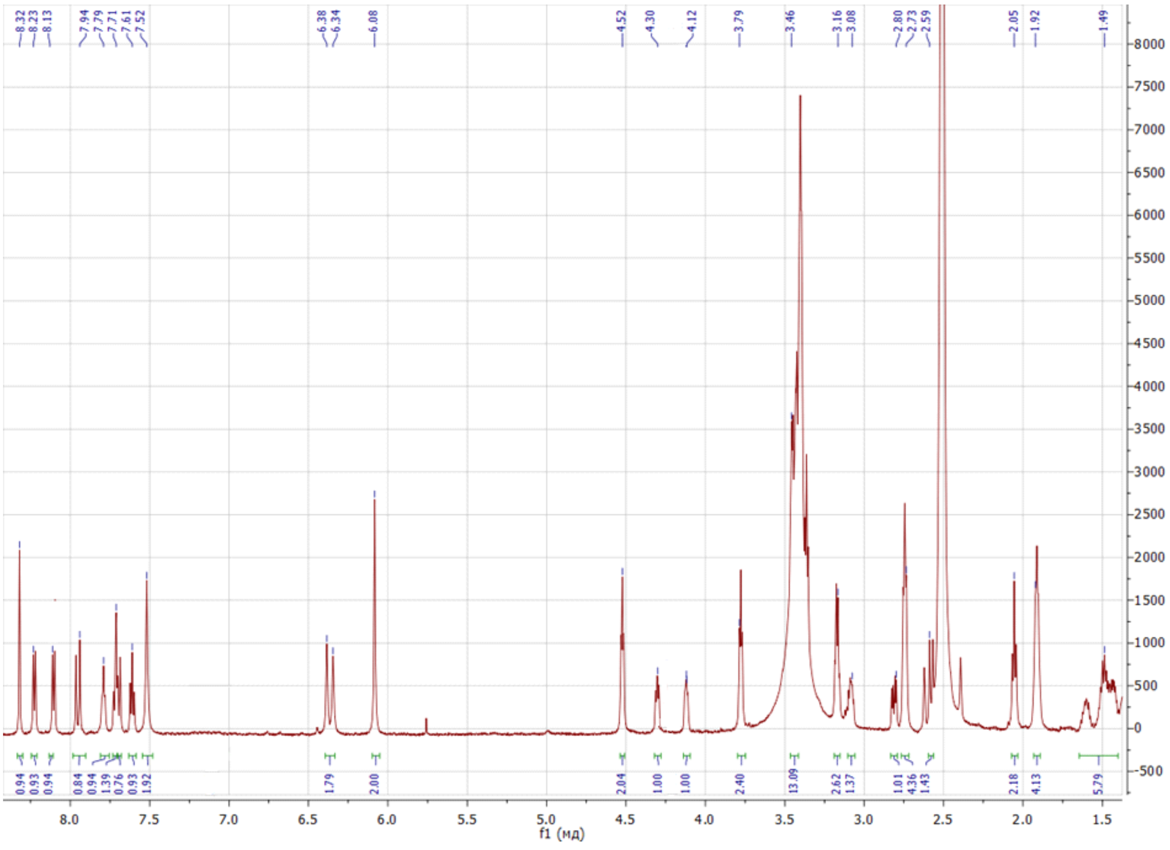

An expansion of the 2D $^{1}$H COSY NMR spectrum (aromatic region)

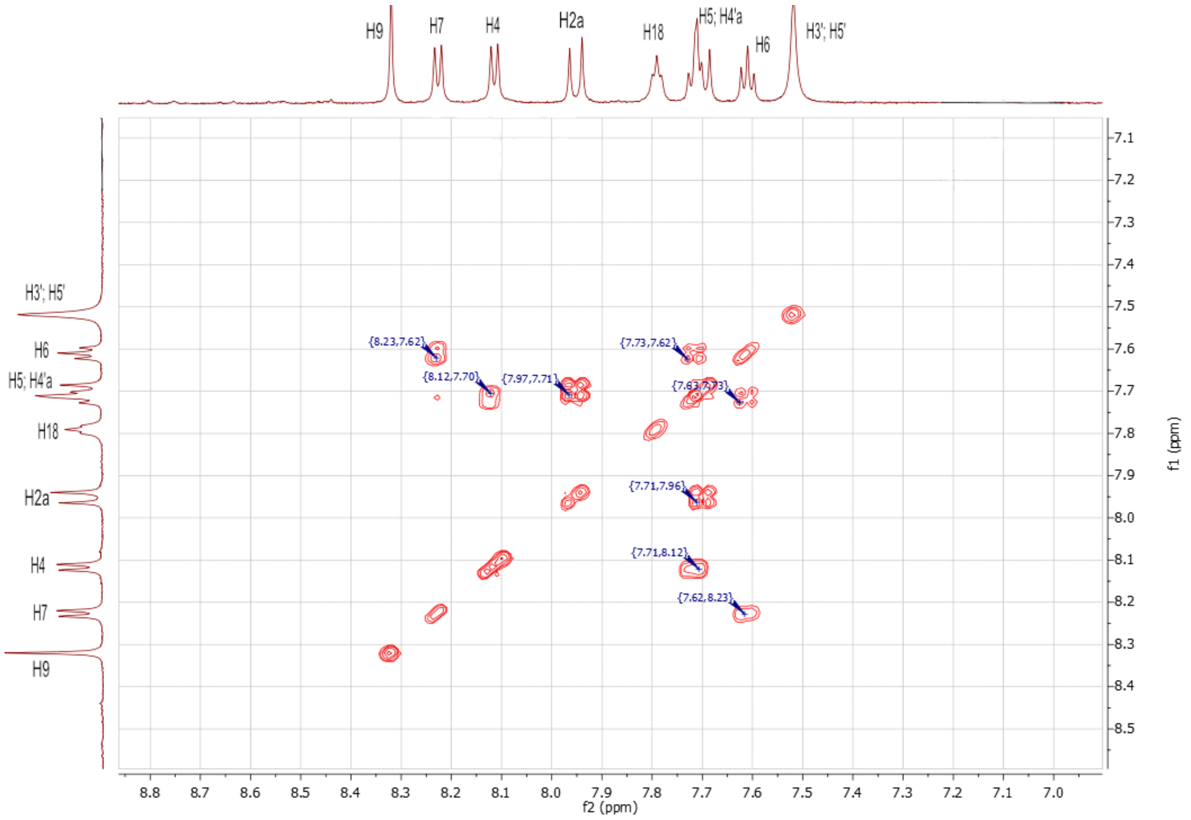

An expansion of the 2D $^{1}$H COSY NMR spectrum (aliphatic region)

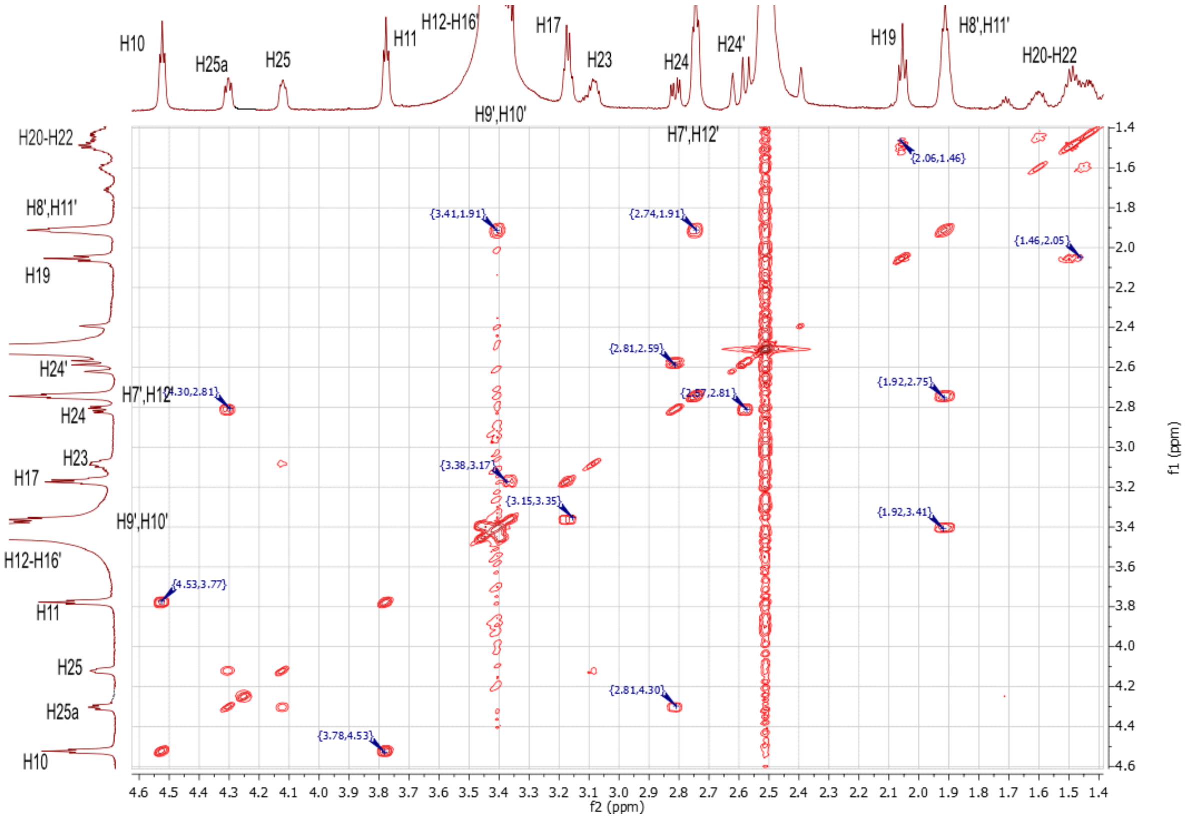

*3-((1-(13-oxo-17-(2-oxohexahydro-1H-thieno[3,4-d]imidazol-4-yl)-3,6,9-trioxa-12-azaheptadecyl)-1H-1,2,3-triazol-4-yl)methyl)-2-(2-(1,2,2,4-tetramethyl-1,2-dihydroquinolin-6-yl)vinyl)benzo[d]thiazol-3-ium 2,2,2-trifluoroacetate* **4d**

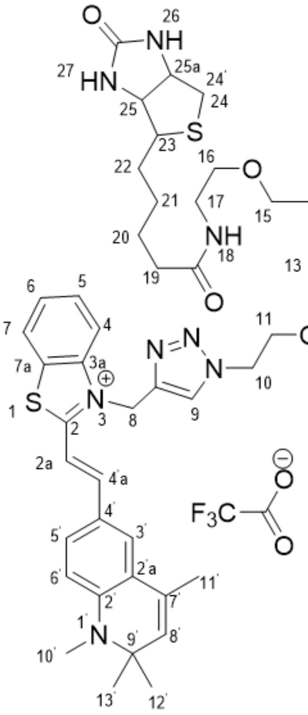

$^1$H spectrum

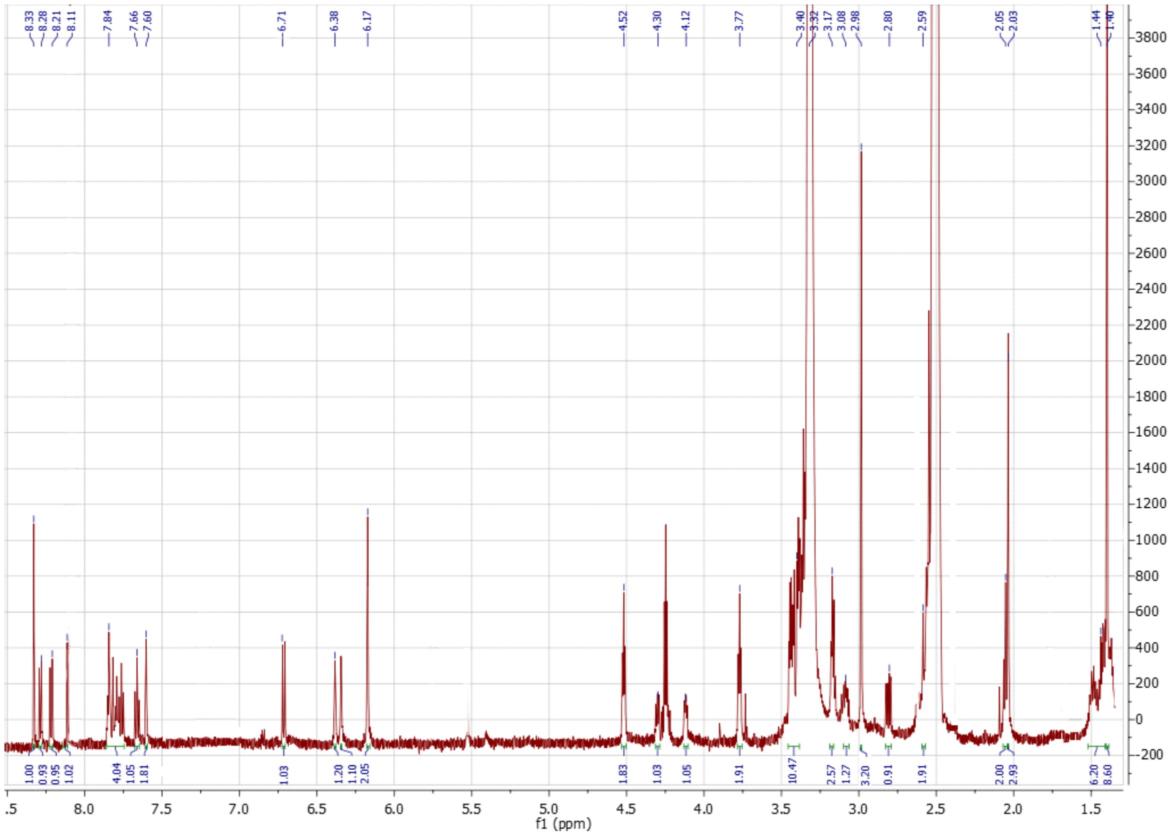

An expansion of the 2D $^{1}$H COSY NMR spectrum (aromatic region)

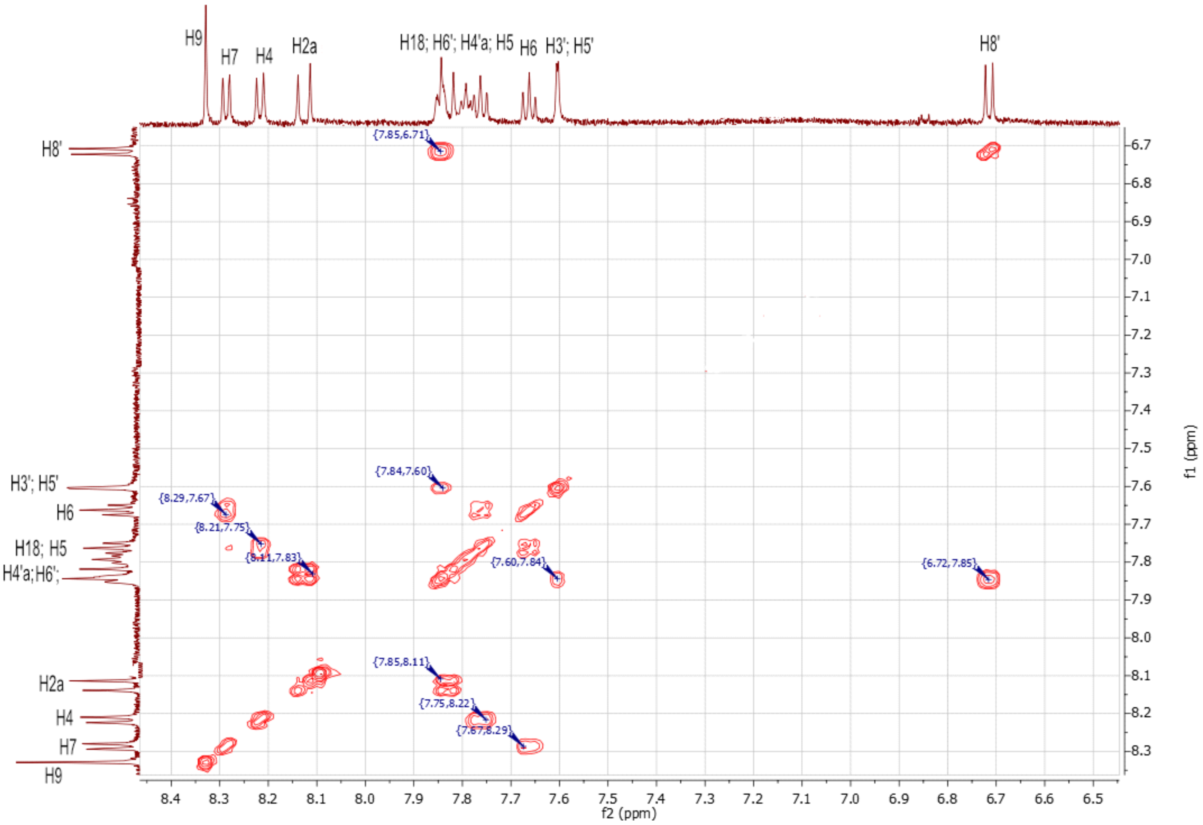

An expansion of the 2D $^{1}$H COSY NMR spectrum (aliphatic region)

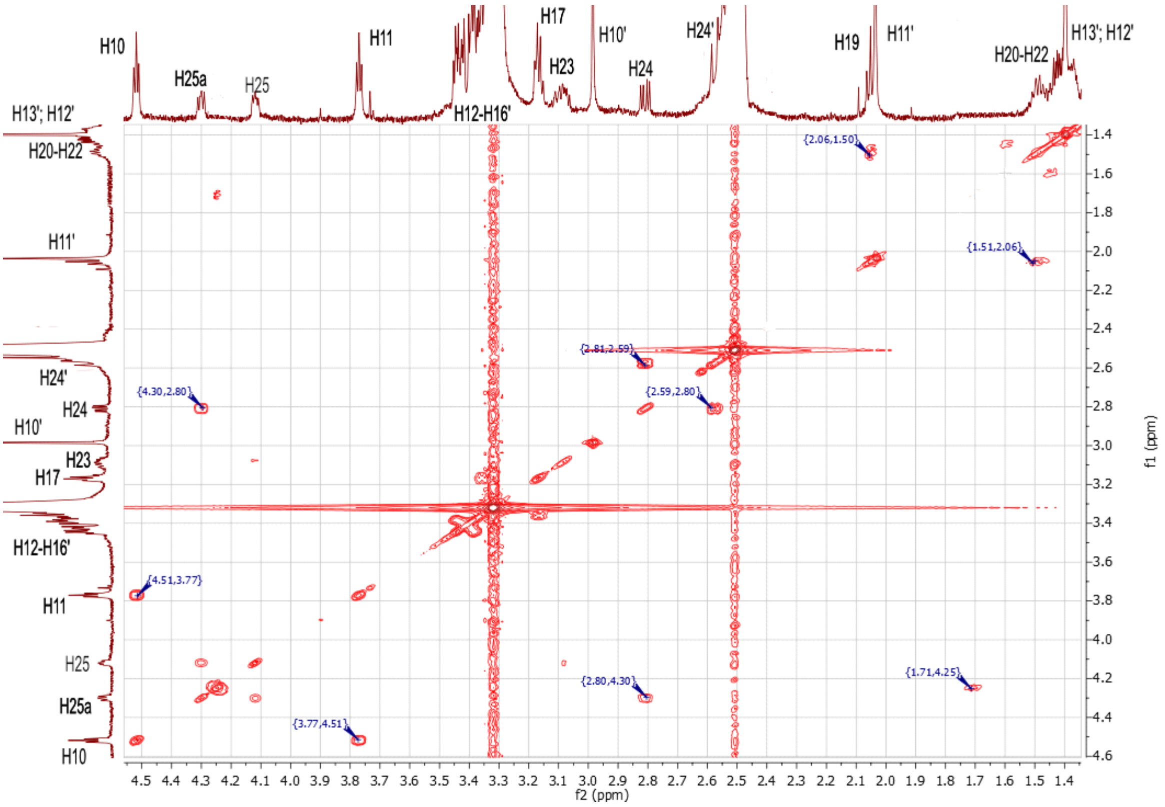

### HPLC and MS data

*(E)-2-((1-methylquinolin-4(1H)-ylidene)methyl)-3-(prop-2-yn-1-yl)benzo[d]thiazol-3-ium bromide* **2a**

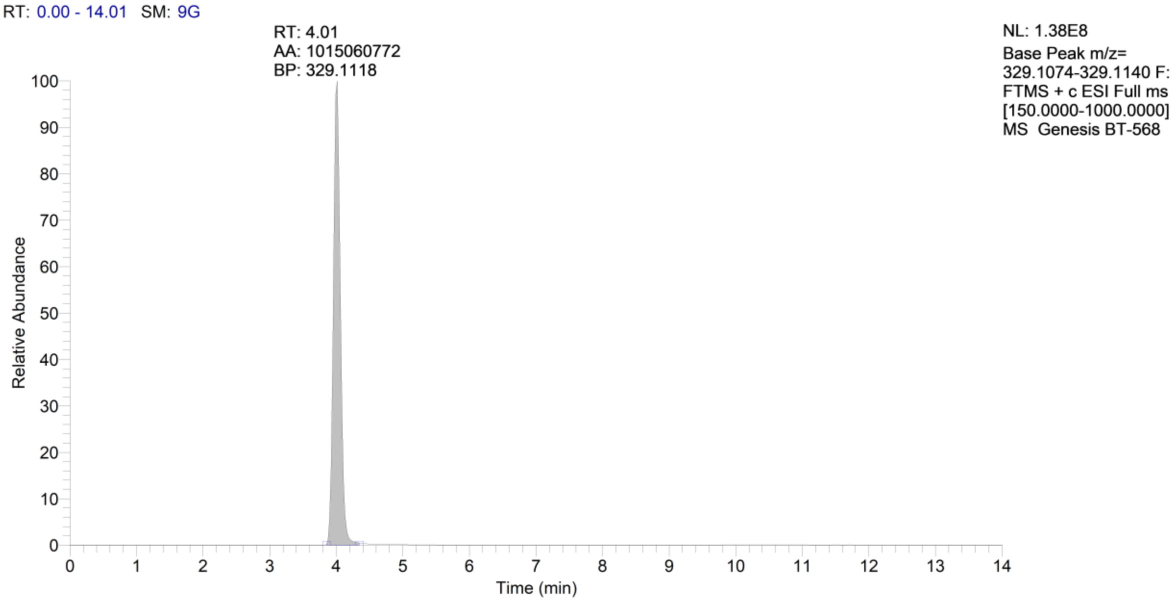

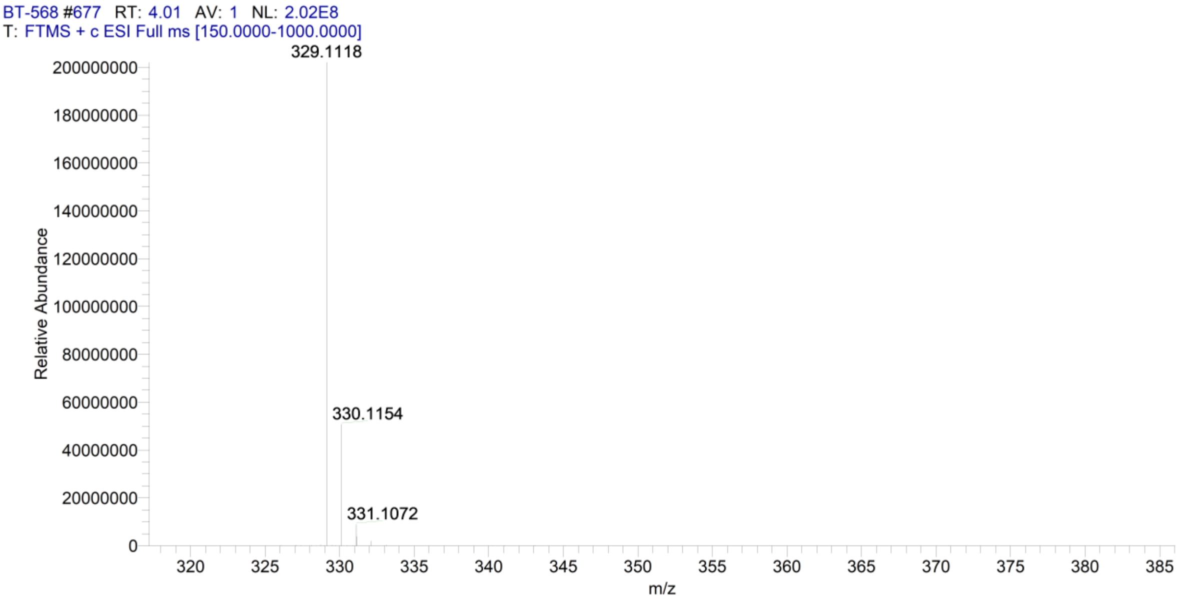

*(E)-2-(2-(4-(Dimethylamino)phenyl)ethenyl)-3-(2-propynyl)benzo[d]thiazolium bromide* **2b**

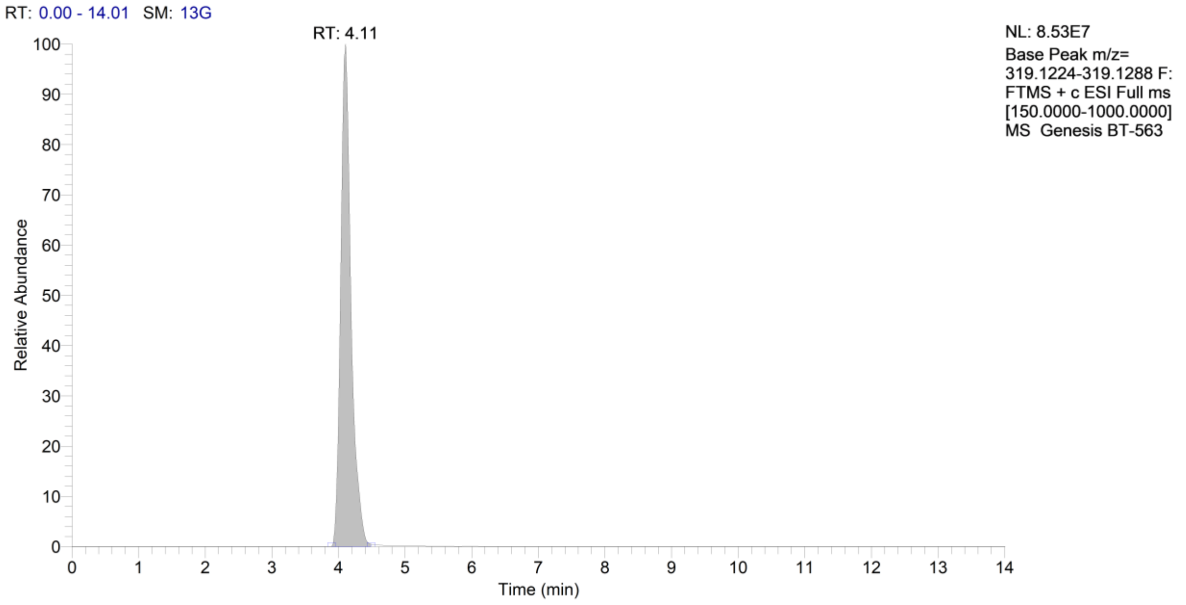

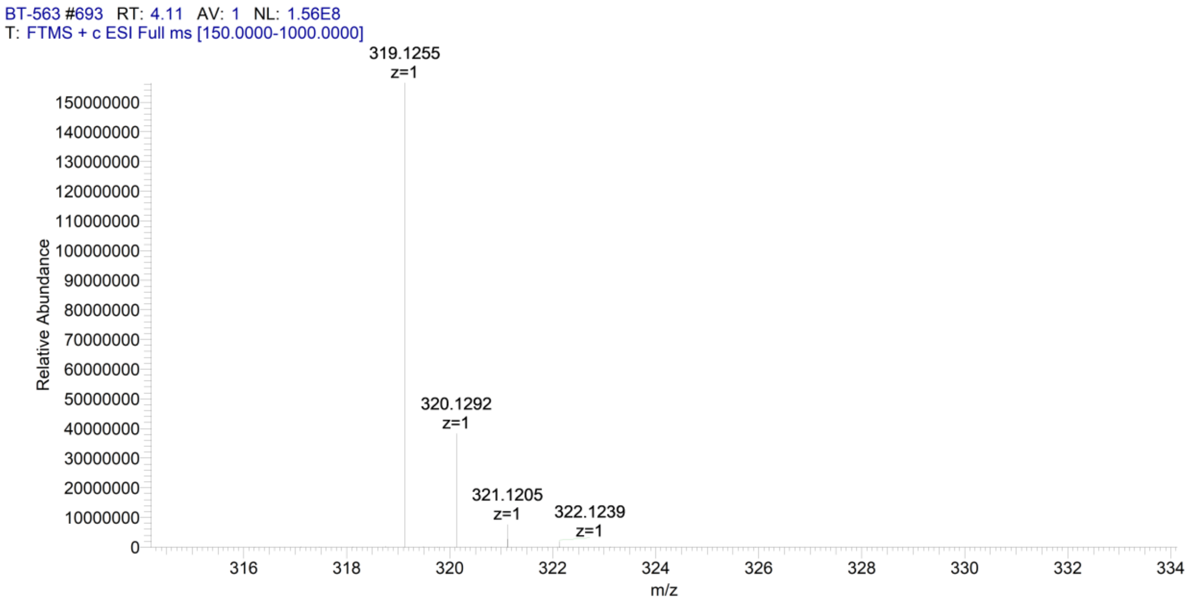

*(E)-3-(prop-2-yn-1-yl)-2-(2-(2,3,6,7-tetrahydro-1H,5H-pyrido[3,2,1-ij]quinolin-9-yl)vinyl)benzo[d]thiazol-3-ium bromide* **2c**

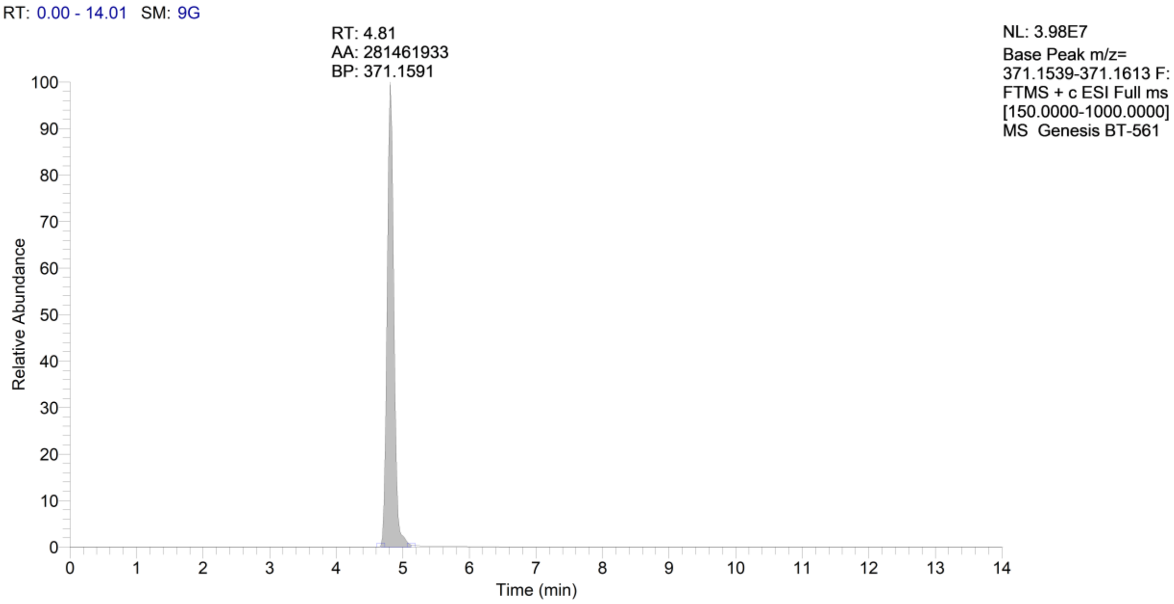

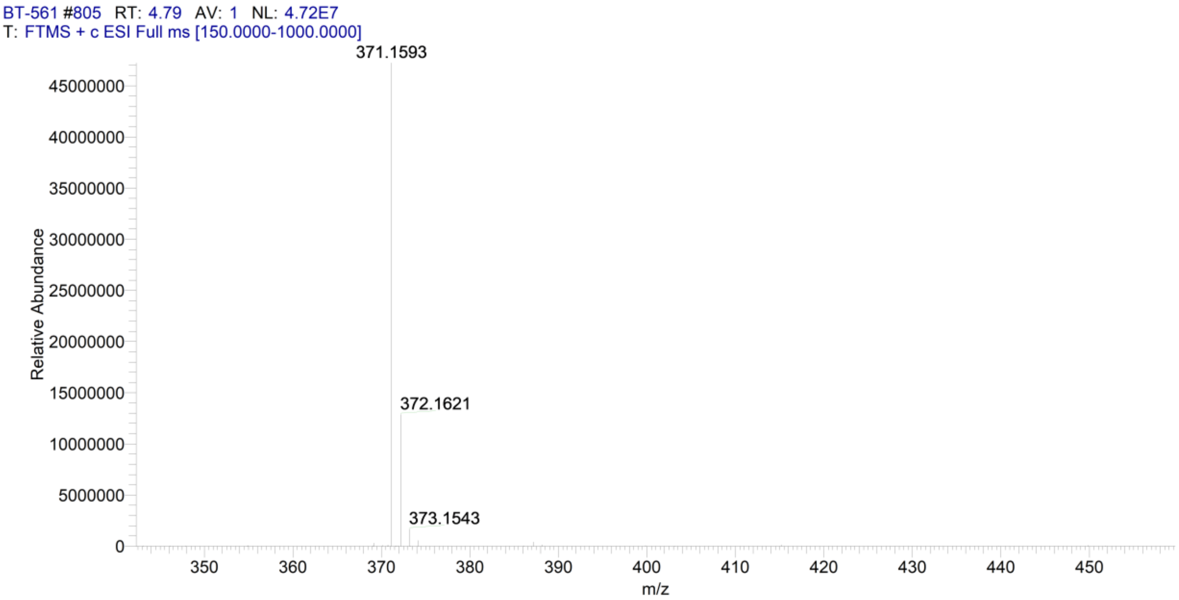

*(E)-3-(prop-2-yn-1-yl)-2-(2-(1,2,2,4-tetramethyl-1,2-dihydroquinolin-6-yl)vinyl)benzo[d]thiazol-3-ium bromide* **2d**

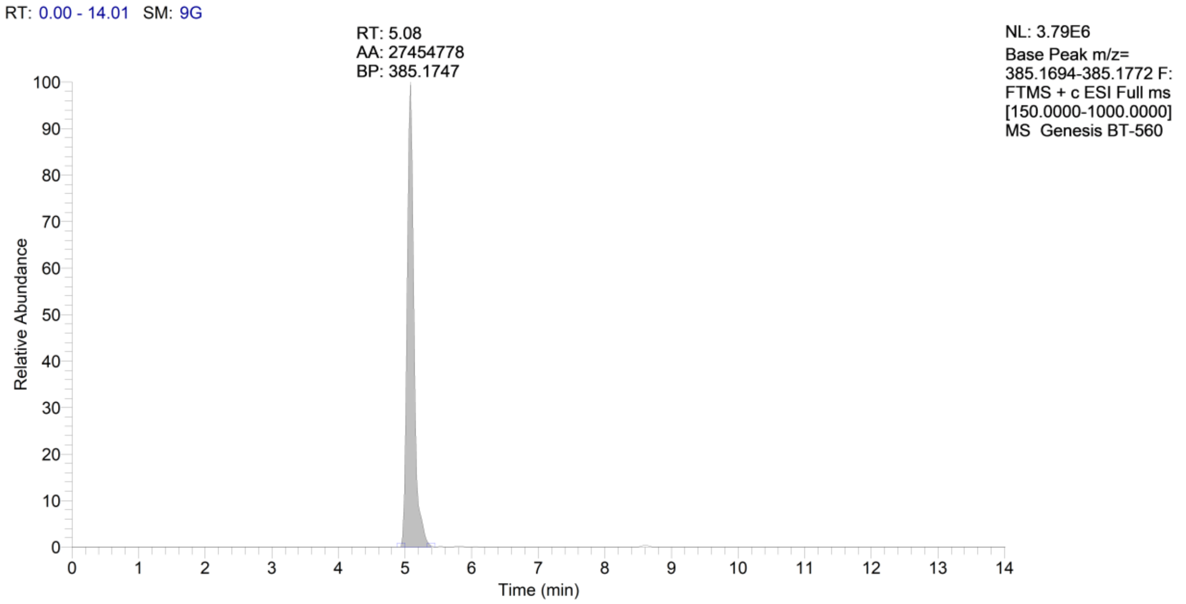

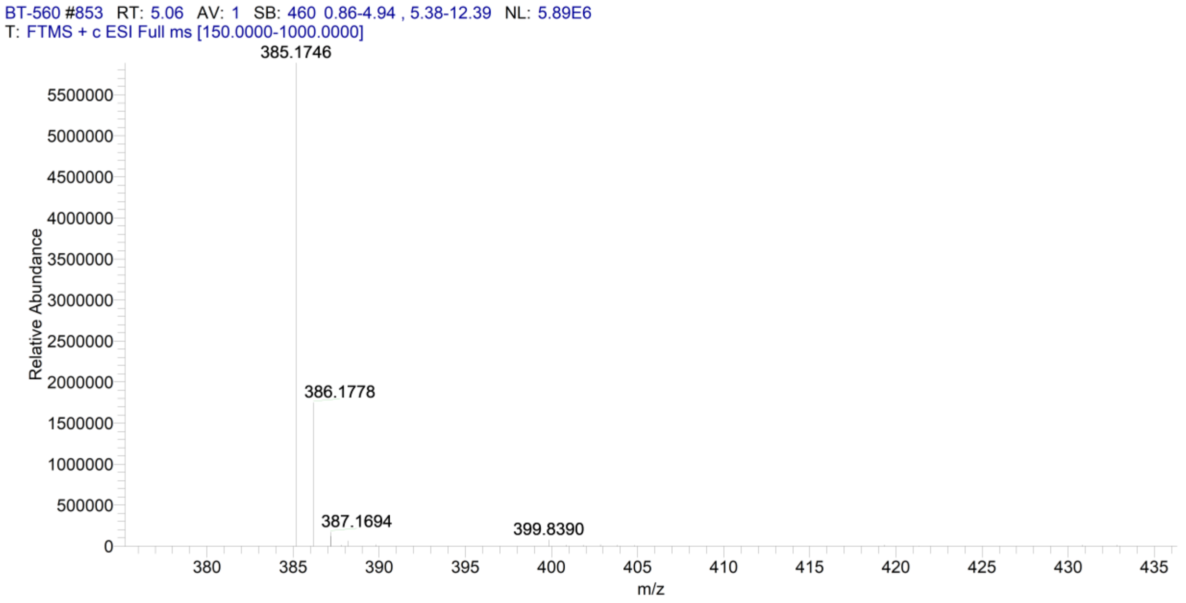

### TO-1 biotin

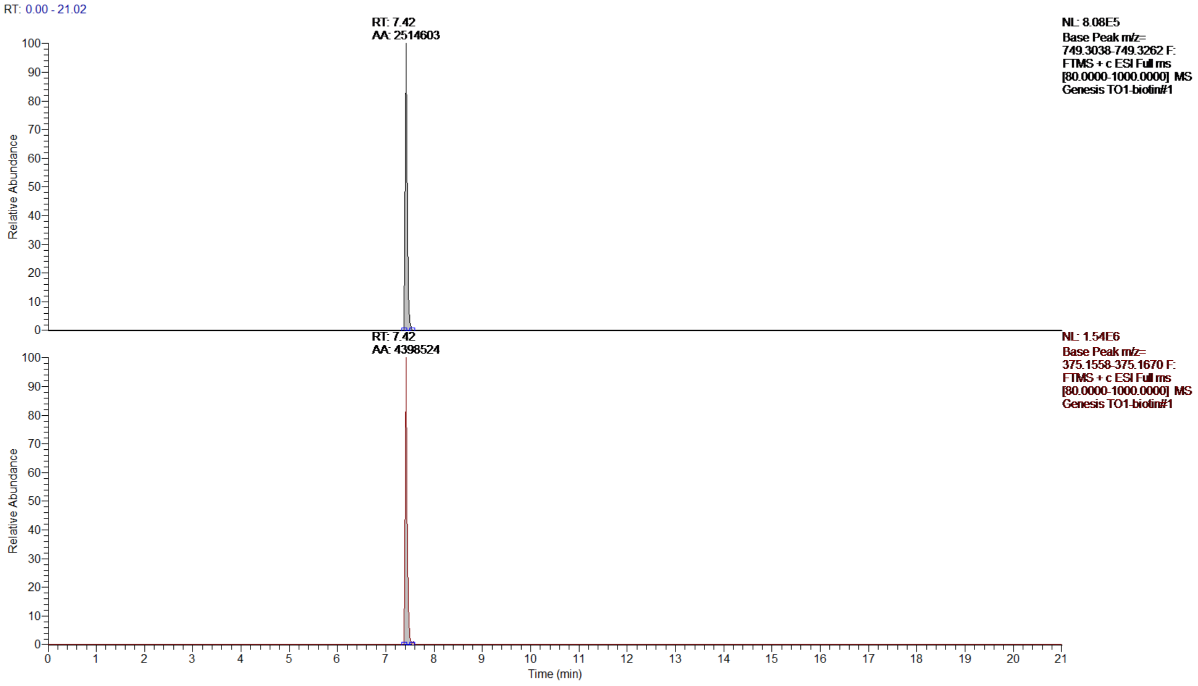

RT: 0.00 - 21.02
RT: 7.42
AA: 2514603
NL: 8.08E5
Base Peak m/z= 749.3038-749.3262 F: FTMS + c ESI Full ms [80.0000-1000.0000] MS Genesis TO1-biotin#1
RT: 7.42
AA: 4398524
NL: 1.54E6
Base Peak m/z= 375.1558-375.1670 F: FTMS + c ESI Full ms [80.0000-1000.0000] MS Genesis TO1-biotin#1
Relative Abundance
Time (min)

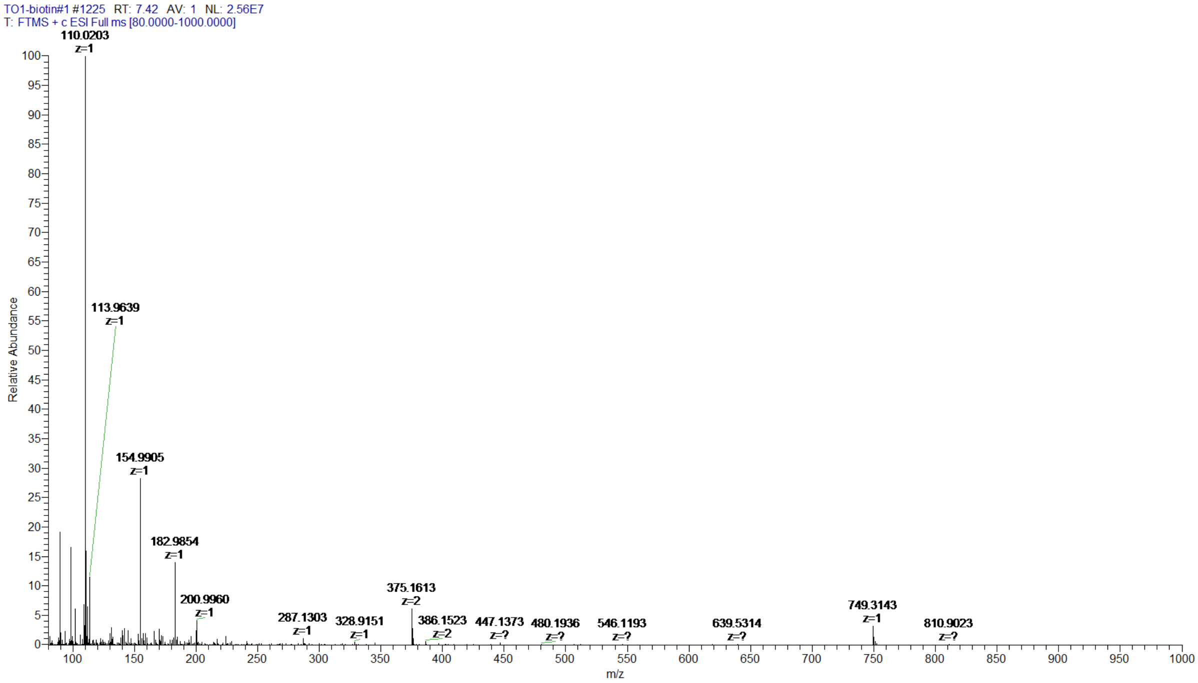

TO1-biotin#1 #1225 RT: 7.42 AV: 1 NL: 2.56E7
T: FTMS + c ESI Full ms [80.0000-1000.0000]
110.0203 z=1
113.9639 z=1
154.9905 z=1
182.9854 z=1
200.9960 z=1
287.1303 z=1
328.9151 z=1
375.1613 z=2
386.1523 z=2
447.1373 z=?
480.1936 z=?
546.1193 z=?
639.5314 z=?
749.3143 z=1
810.9023 z=?
Relative Abundance
m/z

*2-((1-methylquinolin-4(1H)-ylidene)methyl)-3-((1-(13-oxo-17-(2-oxohexahydro-1H-thieno[3,4-d]imidazol-4-yl)-3,6,9-trioxa-12-azaheptadecyl)-1H-1,2,3-triazol-4-yl)methyl)benzo[d]thiazol-3-ium 2,2,2-trifluoroacetate* **4a**

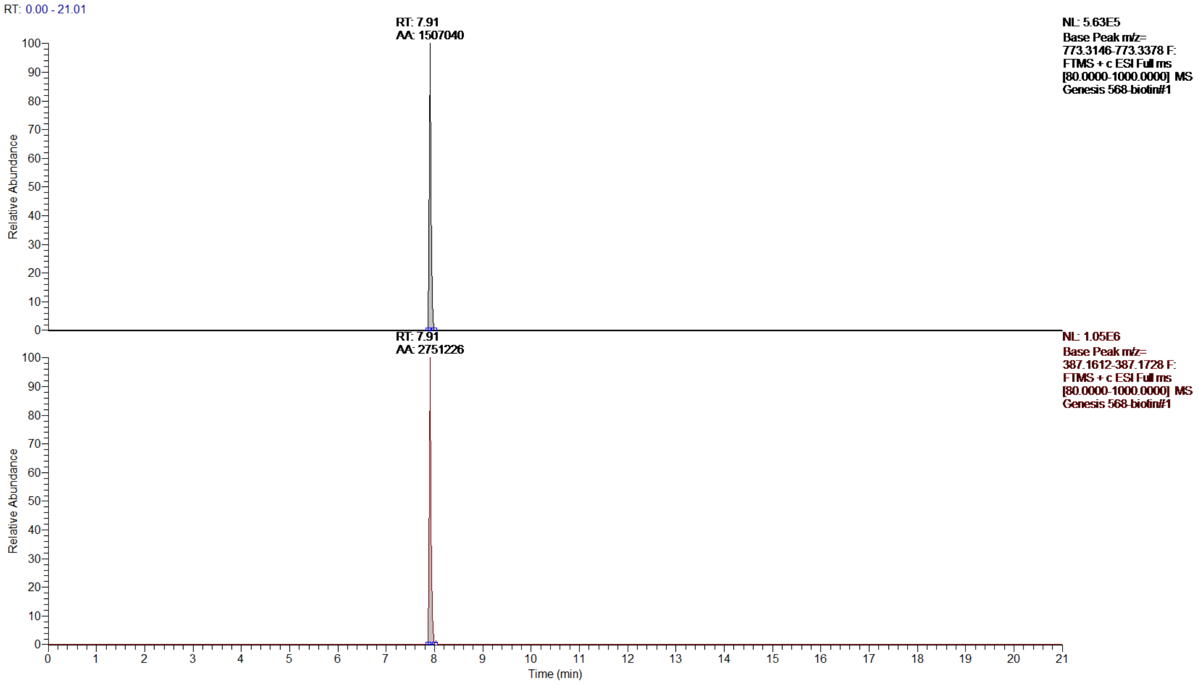

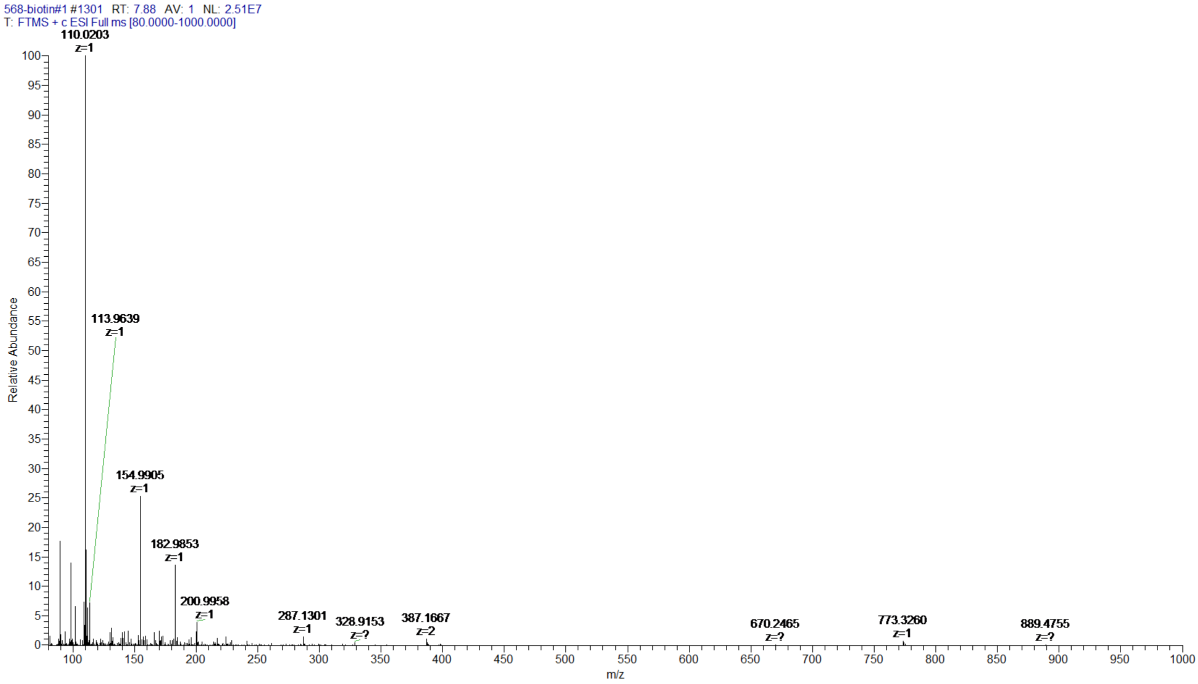

*2-(4-(dimethylamino)styryl)-3-((1-(13-oxo-17-(2-oxohexahydro-1H-thieno[3,4-d]imidazol-4-yl)-3,6,9-trioxa-12-azaheptadecyl)-1H-1,2,3-triazol-4-yl)methyl)benzo[d]thiazol-3-ium 2,2,2-trifluoroacetate* **4b**

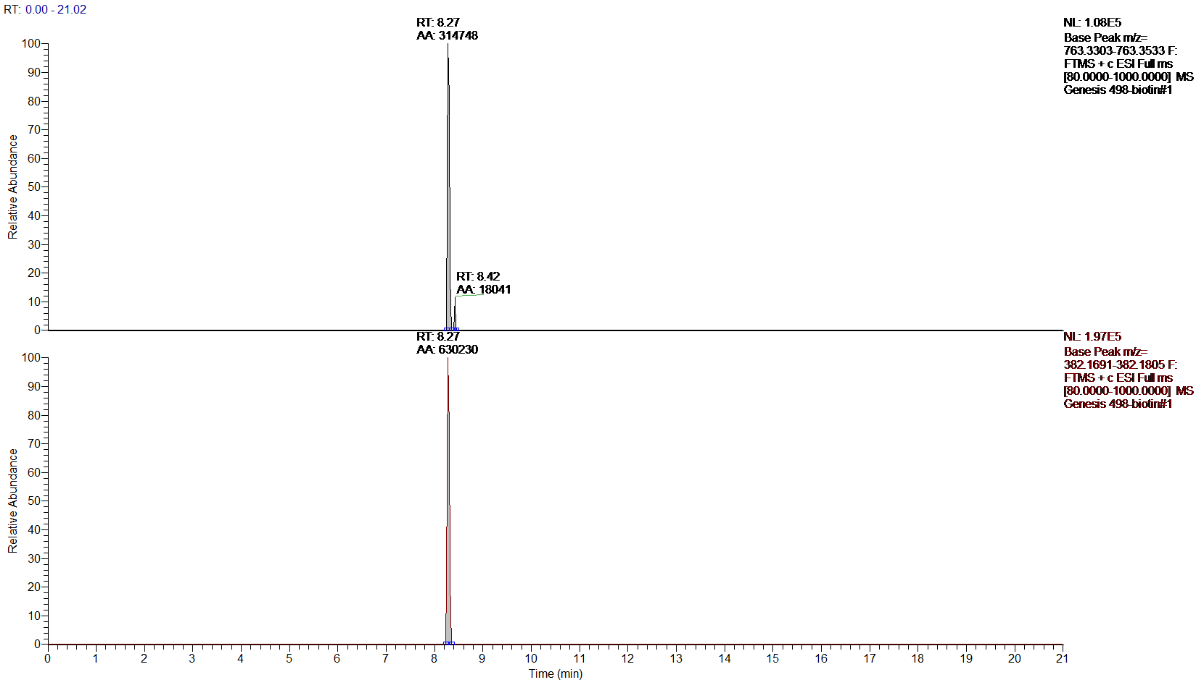

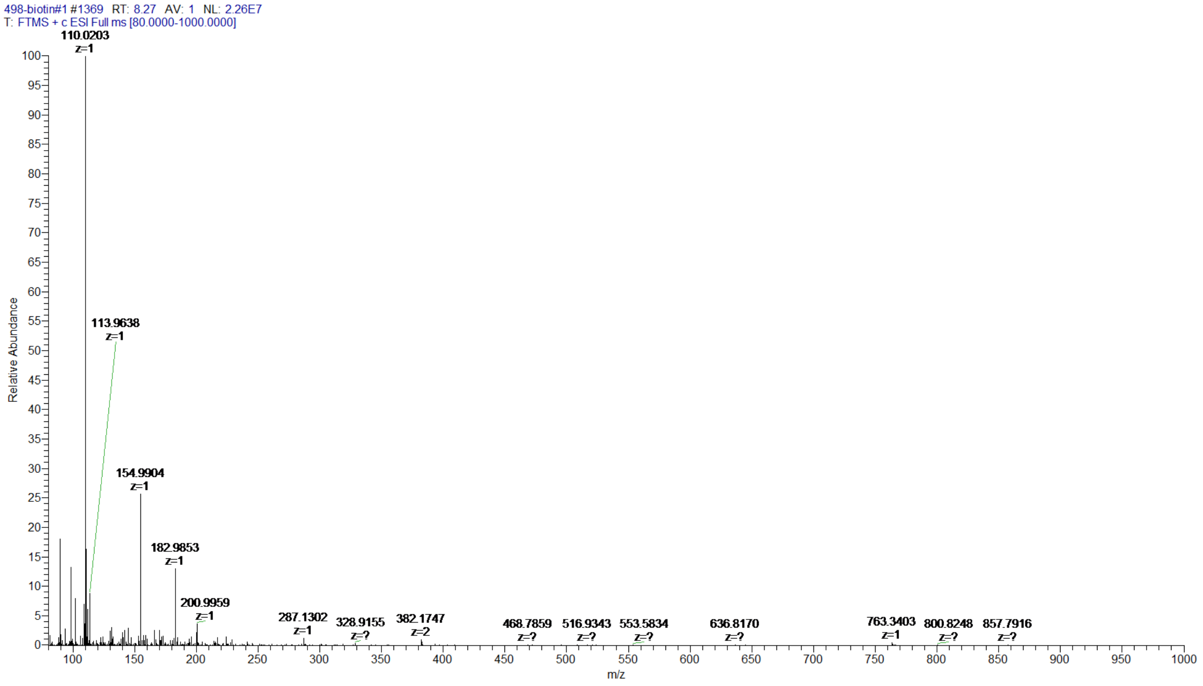

*3-((1-(13-oxo-17-(2-oxohexahydro-1H-thieno[3,4-d]imidazol-4-yl)-3,6,9-trioxa-12-azaheptadecyl)-1H-1,2,3-triazol-4-yl)methyl)-2-(2-(2,3,6,7-tetrahydro-1H,5H-pyrido[3,2,1-ij]quinolin-9-yl)vinyl)benzo[d]thiazol-3-ium 2,2,2-trifluoroacetate* **4c**

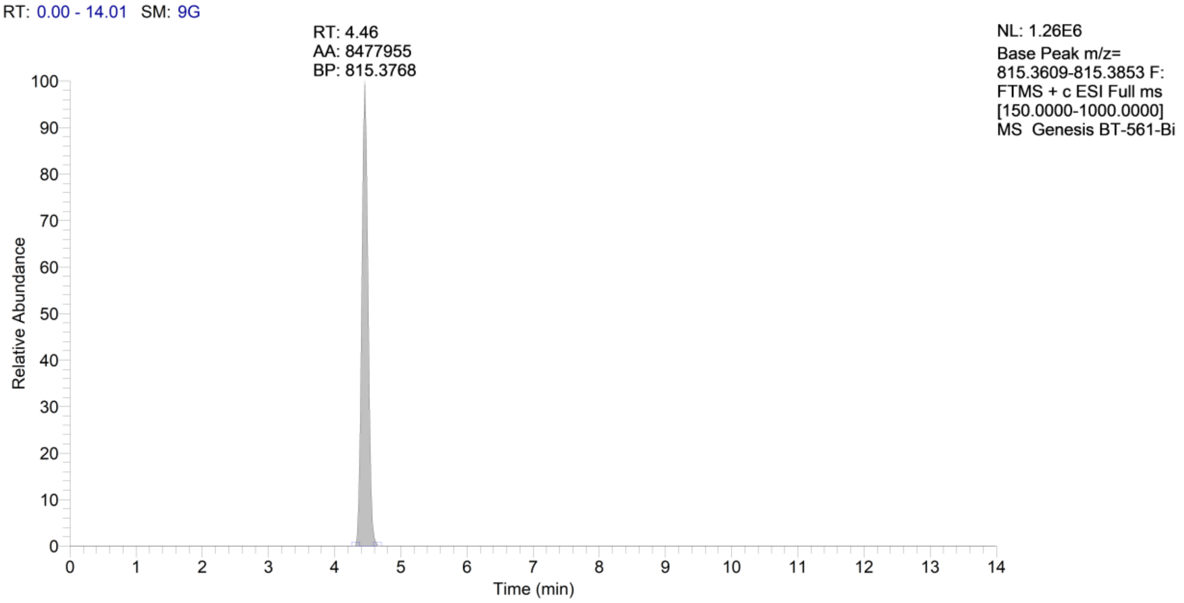

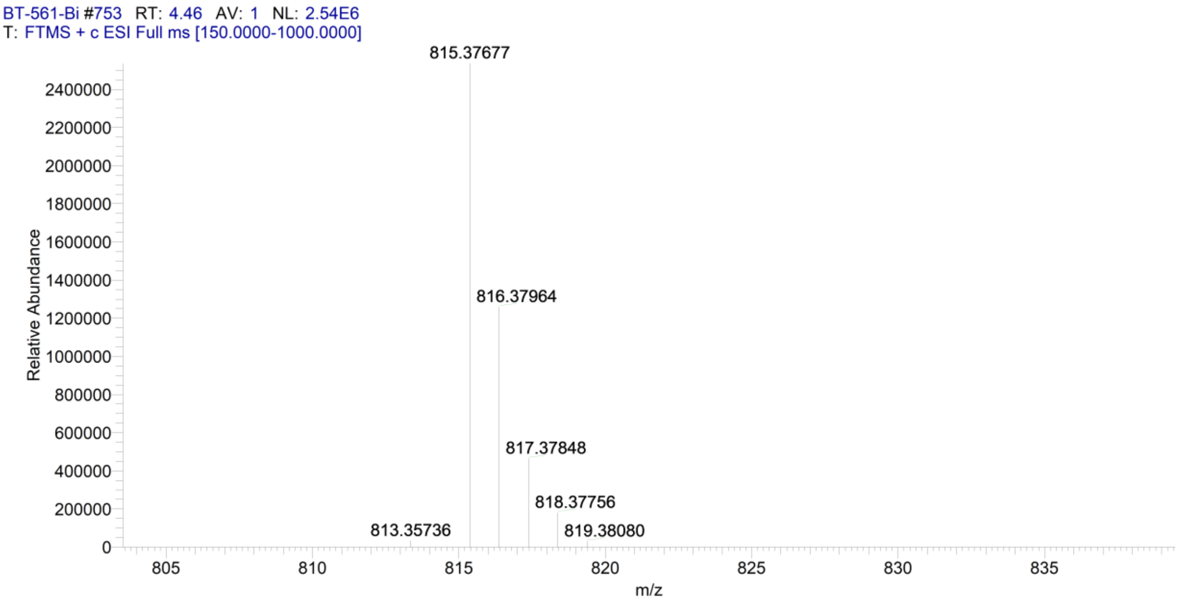

*3-((1-(13-oxo-17-(2-oxohexahydro-1H-thieno[3,4-d]imidazol-4-yl)-3,6,9-trioxa-12-azaheptadecyl)-1H-1,2,3-triazol-4-yl)methyl)-2-(2-(1,2,2,4-tetramethyl-1,2-dihydroquinolin-6-yl)vinyl)benzo[d]thiazol-3-ium 2,2,2-trifluoroacetate* **4d**

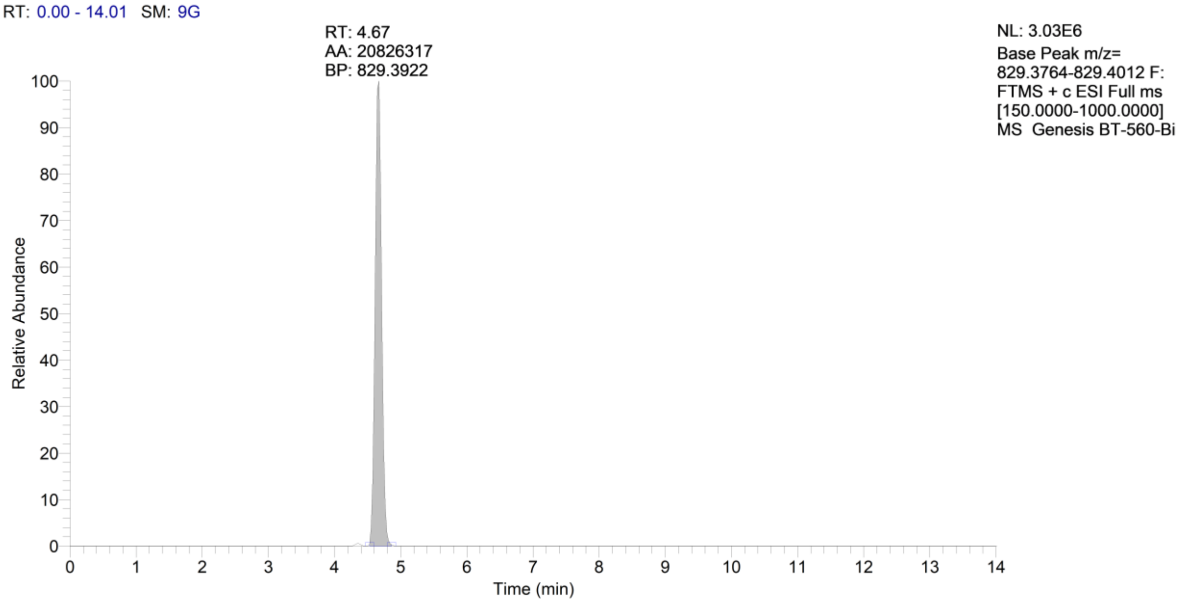

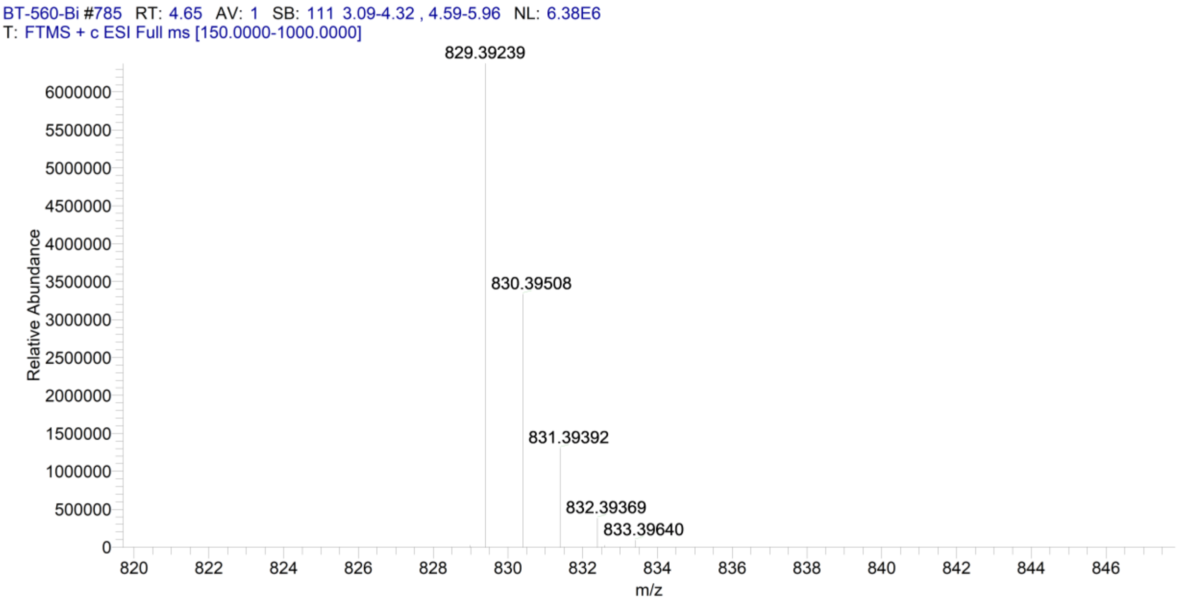

#### References

1. Shen D, *et al.* Design, synthesis and evaluation of a novel fluorescent probe to accurately detect H2S in hepatocytes and natural waters. *Spectrochim Acta A Mol Biomol Spectrosc* **228**, 117690 (2020).

2. Ditmangklo B, Taechalertpaisarn J, Siriwong K, Vilaivan T. Clickable styryl dyes for fluorescence labeling of pyrrolidinyl PNA probes for the detection of base mutations in DNA. *Org Biomol Chem* **17**, 9712-9725 (2019).

3. Sun XL, Stabler CL, Cazalis CS, Chaikof EL. Carbohydrate and protein immobilization onto solid surfaces by sequential Diels-Alder and azide-alkyne cycloadditions. *Bioconjug Chem* **17**, 52-57 (2006).

4. Dolgosheina EV, *et al.* RNA mango aptamer-fluorophore: a bright, high-affinity complex for RNA labeling and tracking. *ACS Chem Biol* **9**, 2412-2420 (2014).

5. Zhao Y, Truhlar DG. The M06 suite of density functionals for main group thermochemistry, thermochemical kinetics, noncovalent interactions, excited states, and transition elements: Two new functionals and systematic testing of four M06-class functionals and 12 other functionals. *Theor Chem Acc.* **120**, 215–241 (2008).

6. Barone V, Cossi M. Quantum Calculation of Molecular Energies and Energy Gradients in Solution by a Conductor Solvent Model. *J Phys Chem A* **102**, 1995–2001 (1998).

7. Singh UC, Kollman PA. An approach to computing electrostatic charges for molecules. *J Comp Chem* **5**, 129-145 (1984).

8. Bayly CI, Cieplak P, Cornell WD, Kollman PA. A Well-Behaved Electrostatic Potential Based Method Using Charge Restraints For Determining Atom-Centered Charges: The RESP Model. *J Phys Chem* **97**, 10269-10280 (1993).

9. Gaussian 16, Revision C.01, Frisch MJ, Trucks GW, Schlegel HB, Scuseria GE, Robb MA, Cheeseman JR, Scalmani G, Barone V, Petersson GA, Nakatsuji H, Li X, Caricato M, Marenich AV, Bloino J, Janesko BG, Gomperts R, Mennucci B, Hratchian HP, Ortiz JV, Izmaylov AF, Sonnenberg JL, Williams-Young D, Ding F, Lipparini F, Egidi F, Goings J, Peng B, Petrone A, Henderson T, Ranasinghe D, Zakrzewski VG, Gao J, Rega N, Zheng G, Liang W, Hada M, Ehara M, Toyota K, Fukuda R, Hasegawa J, Ishida M, Nakajima T, Honda Y, Kitao O, Nakai H, Vreven T, Throssell K, Montgomery JA Jr, Peralta JE, Ogliaro F, Bearpark MJ, Heyd JJ, Brothers EN, Kudin KN, Staroverov VN, Keith TA, Kobayashi R, Normand J, Raghavachari K, Rendell AP, Burant JC, Iyengar SS, Tomasi J, Cossi M, Millam JM, Klene M, Adamo C, Cammi R, Ochterski JW, Martin RL, Morokuma K, Farkas O, Foresman JB, Fox DJ. Gaussian, Inc., Wallingford CT (2016).

10. Abagyan R, Totrov M, Kuznetsov D. ICM, a new method for protein modeling and design: applications to docking and structure prediction from the distorted native conformation. *J Comput Chem* **15**, 488-506 (1994).

11. Case DA, Belfon K, Ben-Shalom IY, Brozell SR., Cerutti DS, Cheatham III TE, Cruzeiro VWD, Darden TA, Duke RE, Giambasu G, Gilson MK, Gohlke H, Goetz AW, Harris R, Izadi S, Kasavajhala K, Kovalenko A, Krasny R, Kurtzman T, Lee TS, LeGrand S, Li P, Lin C, Liu J, Luchko T, Luo R, Man V, Merz KM, Miao Y, Mikhailovskii O, Monard G, Nguyen H, Onufriev A, Pan F, Pantano S, Qi R, Roe DR, Roitberg A, Sagui C, Schott-Verdugo S, Shen J, Simmerling CL, Skrynnikov N, Smith J, Swails J, Walker RC, Wang J, Wilson L, Wolf RM, Wu X, York DM, Kollman PA. AMBER 2020, University of California, San Francisco (2020).

12. Izadi S, Onufriev AV. Accuracy limit of rigid 3-point water models. *J Chem Phys* **145**, 074501 (2016).

13. Perez A, Marchan I, Svozil D, Sponer J, Cheatham TE, Laughton CA, Orozco M. Refinement of the AMBER Force Field for Nucleic Acids: Improving the Description of alpha/gamma Conformers. *Biophys J* **92**, 3817–3829 (2007).

14. Yildirim I, Stern HA, Kennedy SD, Tubbs JD, Turner DH.. Reparameterization of RNA chi Torsion Parameters for the AMBER Force Field and Comparison to NMR Spectra for Cytidine and Uridine *J Chem Theory Comput* **6**, 1520–1531 (2010).

15. Onufriev A, Case DA, Bashford D. Effective Born radii in the generalized Born approximation: the importance of being perfect. *J Comput Chem* **23**, 1297–1304 (2002).

16. Penuelas-Urquides K, Villarreal-Trevino L, Silva-Ramirez B, Rivadeneyra-Espinoza L, Said-Fernandez S, de Leon MB. Measuring of Mycobacterium tuberculosis growth. A correlation of the optical measurements with colony forming units. *Braz J Microbiol* **44**, 287-289 (2013).

17. Aseev LV, Koledinskaya LS, Bychenko OS, Boni IV. Regulation of Ribosomal Protein Synthesis in Mycobacteria: The Autogenous Control of rpsO. *Int J Mol Sci* **22**, (2021).

18. Bychenko OS, Skvortsova YV, Grigorov AS, Azhikina TL. Use of Genetically Encoded Fluorescent Aptamers for Visualization of Mycobacterium tuberculosis Small RNA MTS1338 in Infected Macrophages. *Dokl Biochem Biophys* **493**, 185-189 (2020).

19. Grigorov A, *et al.* Small RNA F6 Provides Mycobacterium smegmatis Entry into Dormancy. *Int J Mol Sci* **22**, (2021).
